# Functional alterations of immune gene expression in ICU and non-ICU patients with Legionnaires’ disease, a prospective observational study

**DOI:** 10.64898/2026.03.06.709905

**Authors:** Camille Allam, William Mouton, Chloé Albert-Vega, Marine Ibranosyan, Christophe Ginevra, Ghislaine Descours, Laetitia Beraud, Annelise Chapalain, Abdelrahim Zoued, Laurent Argaud, Arnaud Friggeri, Vanessa Labeye, Yvan Jamilloux, Anne-Claire Lukaszewicz, Guillaume Monneret, Jonathan Lopez, Nathalie Freymond, Gerard Lina, Patricia Doublet, Jean-Christophe Richard, Fabienne Venet, Florence Ader, Sophie Trouillet-Assant, Sophie Jarraud

## Abstract

Legionnaires’ disease (LD), a pneumonia caused by *Legionella pneumophila* intracellular bacterium, leads to intensive care unit (ICU) admission in 20-40% of cases. While these ICU-LD patients display severe lung injury or septic shock, their functional immune response remains poorly understood. The present study aimed, through a large immune gene expression assessment, to improve the understanding of immune cell functionality after whole blood LPS *ex vivo* stimulation in ICU-LD patients compared with non-ICU.

Both ICU and non-ICU-LD displayed altered gene expression indicating that both patients’ immune cells are less able to respond to the LPS *ex vivo* stimulus than a healthy population. ICU-LD patients had 1.6-fold greater number of less-expressed genes (35/93 vs 22/93, p=0.039), and lower Log2(FC) of these genes (median [IQR]: -1.9 [-2.6;-1.5] *vs* -1.2 [-1.7;-0.9], p=0.0011) than non-ICU-LD. Seven genes were significantly less expressed by ICU-LD patients (*IRF7*, *MX1*, *NFKBI2*, *NFKBIA*, *RELB*, *SRC*, *TIM3*; p-value range: 0.029-0.0080).

Top five gene ontology biological processes, subcellular localisations, and reactome pathways (STRING database) uniquely enriched in ICU-LD-patients and related less-expressed genes were cellular response to LPS (*CCL2*, *NFKBIA*, *IRAK2*, *TIM3, SRC*, *NFKB1*), regulation of IFN-β production (*IRF7*, *RIG1*, *OAS2*, *RELB*), I-κB/NF-κB complex (*NFKBIA*, *NFKB1*, *NFKB2*), IFN regulatory factor complex (*RIG1*, *IRF7*), and TRAF6-mediated NF-κB activation pathway (*NFKBIA*, *NFKB1*, *NFKB2*, *RIG1*).

Immune gene expression alterations in LD after LPS stimulation were found herein, with more pronounced alterations in ICU-LD patients. A reduced expression of key genes and pathways involved in controlling *Legionella* proliferation in ICU-LD patients may contribute to increased disease severity.

## Introduction

Legionnaires’ disease (LD), a pneumonia caused by *Legionella pneumophila* intracellular bacterium, leads to intensive care unit (ICU) admission in 20-40% of cases [1,2]. The case fatality ratio is around 25% [3,4] in these ICU-LD patients, while this is around 8% in the general population [5]. Among ICU-LD patients, 47-77% suffer from severe lung injury that necessitates mechanical ventilation (MV) [3,6–8] and 33-53% develop multi-organ failure or septic shock (SS) [3,9]. As the clinical course of ICU-LD patients is difficult to predict [10], dedicated tools and tailored approaches are needed to better assess prognosis and optimise infection management.

We previously reported that ICU-LD patients on MV or with a high Sepsis-related Organ Failure Assessment (SOFA) score had high initial concentration of seven bloodstream pro-inflammatory mediators [11]. Moreover, after leukocyte *ex vivo* stimulation with a non-specific mitogen, ICU-LD patients had functional alterations in the release of pro-inflammatory cytokines, TH-1 and TH-2 cytokines, chemokines, as well as growth factors, and, after lipopolysaccharide (LPS) stimulation, TNF-α. Although these results open ways to better understand the functional immune alterations in ICU-LD patients, a more comprehensive understanding of the transcriptional immunological mechanisms and pathways underlying the progression toward severe LD is needed. Furthermore, we did not compare this functional response to that of patients showing mild LD, or other critically ill populations unrelated to LD (*e.g.* SS or ICU-Coronavirus Disease-19 patients), that are known to display deep functional alterations in anti-microbial immunological pathways, linked to disease worsening [12].

The present study aimed, through a large immune gene expression assessment, to improve the understanding of immune cell functionality after whole blood LPS *ex vivo* stimulation in ICU-LD patients compared with non-ICU and LD-unrelated-SS patients.

## Methods

### Study design and patients

This study included 14 LD patients from the ProgLegio prospective observational study (NCT03064737) who were, at inclusion (September 2019 and July 2020), admitted to either conventional wards (non-ICU-LD, n=8) or ICUs (ICU-LD, n=6) of the Hospices Civils de Lyon (Lyon, France). They were enrolled a median [interquartile range, IQR] 2 [0;4] days post- LD-diagnosis. Whole blood was taken on day-2 after enrolment for immune functional assay (IFA). These were compared to healthy volunteers (HVs) and LD-unrelated-SS patients reported by Albert Vega *et al.* [13] (for both of these groups, stimulation and gene expression measurement was similar to that for LD in the ProgLegio study). The median [IQR] age of HVs was 55 [51;57] years, and 50% were male. Samples from these were obtained from the French national blood services (*Etablissement Français du Sang*, EFS). LD-unrelated-SS patients admitted to the ICU of the Edouard Herriot hospital (Hospices Civils de Lyon) were included (June-September 2017) in the Immunosepsis-4 prospective observational study (NCT02803346) [13], within a median [IQR] 1 [0;1] day post SS-diagnosis and were stimulated at day-3 after inclusion. The presence of a SS was defined according to the sepsis-3 definition (*i.e.* persistent hypotension requiring vasopressors to maintain mean an arterial pressure of ≥65mmHg and a serum lactate level >2mmol/L), for both ICU-LD and SS patients.

### Ethics and regulatory issues

LD patients were included in the ProgLegio study; SS patients were included in the Immunosepsis-4 study, with previously described ethical and regulatory considerations ([11,13] and Supplementary Methods). HVs consented for the use of their donated blood for research, following EFS standard procedures.

### IFA

IFA were performed on heparinised whole blood from LD and SS patients, as well as HVs, within 3h after collection. *Ex vivo* stimulation was performed on 1mL of blood incubated for 24h at 37°C in standardised TruCulture® tubes (Rules Based Medicine, IQUVIA Laboratories, Austin, TX, USA) prefilled with ultrapure *E. coli* lipopolysaccharide (LPS; 100ng/mL, *E. coli* O55:B5), or culture medium (NUL; Gibco RPMI 1640 medium, Fisher Scientific, Illkirch-Graffenstaden, France), as described by Albert Vega *et al.* [13]. After 24h incubation, the buffy coat was resuspended in 2mL TRI reagent LS (Sigma-Aldrich, Deisenhofen, Germany) and rested for 10min at room temperature before storage at -80°C for RNA extraction (Supplementary Methods).

### Gene expression (mRNA) nanoString measurement

The expression of 93 immune-related genes (Table S1), and three housekeeping genes used for normalisation, (*HPRT1*, *DECR1*, and *TBP*) were measured using the nCounter Analysis System (nanoString Technologies, Bruker, Bothell WA, USA) following a protocol developed by Mouton *et al.* [14]. The gene panel was chosen by selecting 44 representative genes of the healthy immune response after blood stimulation and 49 genes whose expression vary during SS [13]. Gene expression (mRNA) count was analysed using nSolver software (v4.0, nanoString Technologies; Supplementary Methods). Ratios of each gene expression count between the LPS-stimulated condition and the NUL condition were first calculated for LD patients and HVs. Gene expression count ratios for patients were divided by the geometric mean of those for HVs for each gene after LPS stimulation to assess gene expression fold change (FC) between patients and HVs, as described by Mouton *et al.* [14]. Differentially expressed genes (DEGs) were those with a ≥2-FC between LD patients and HVs.

### Pathway analysis

We used the Search Tool for the Retrieval of Interacting Genes/Proteins (STRING) database (https://string-db.org) to identify gene networks, Gene Ontology (GO) biological processes, subcellular localisation, and reactome pathways in which less-expressed genes were found. STRING analysis is detailed in Supplementary Methods.

### Statistical analysis

Results were expressed as median [IQR] for numerical data, or n (%) otherwise. The false discovery rate (FDR; Benjamini-Hochberg method) was used to adjust p-values in all gene analyses. The FC of genes was log(2) transformed, and FDR was log10-transformed for visualisation in Volcano-plots. The Mann-Whitney, Kruskal-Wallis, and Chi-squared tests were used; accordingly, p-values, and Benjamini-Hochberg adjusted p-values in case of multiple two-by-two comparisons, <0.05 were considered significant. Statistical tests were conducted using GraphPad Prism (v10.6.1; GraphPad software, San Diego, CA, USA). Missing data were removed for statistical analysis.

## Results

### ICU-LD and non-ICU-LD patients clinical and laboratory characteristics

A total of 14 LD patients (n=8 non-ICU, and n=6 ICU) were included. Compared to non-ICU-LD patients, on the day of inclusion ICU-LD patients had a greater SOFA score (median [IQR]: 8 [7;10] *vs* 1 [0;1], p<0.001), lower mean blood pressure (90 [72–98] *vs* 103 [102–110] mmHg, p<0.001), higher *Legionella* DNA load in the lungs (2373 [233;23915] *vs* 60.3 [1.4;214] GU/reaction, p=0.026) and serum (12.0 [1.5;33.0] *vs* 0.1 [0.1-0.1] GU/reaction, p=0.003), and longer hospital length of stay (20 [16;24] *vs* 8 [7;10] days, p=0.018; Table 1 near here). ICU-LD patients’ immune response to LPS stimulation was also compared to a group of patients suffering from an LD-unrelated-SS with a non-pulmonary primary site of infection (n=14; abdominal n=5, skin and soft tissues n=4, urinary tract n=3, catheter infection, and osteo-arthritis n=1). The characteristics of ICU-LD patients, including those meeting the definition of SS (n=3), were similar to LD-unrelated-SS patients (except mean blood pressure, PaO2/FiO2, and lactates that were significantly different; Table S2).

**Table 1:**
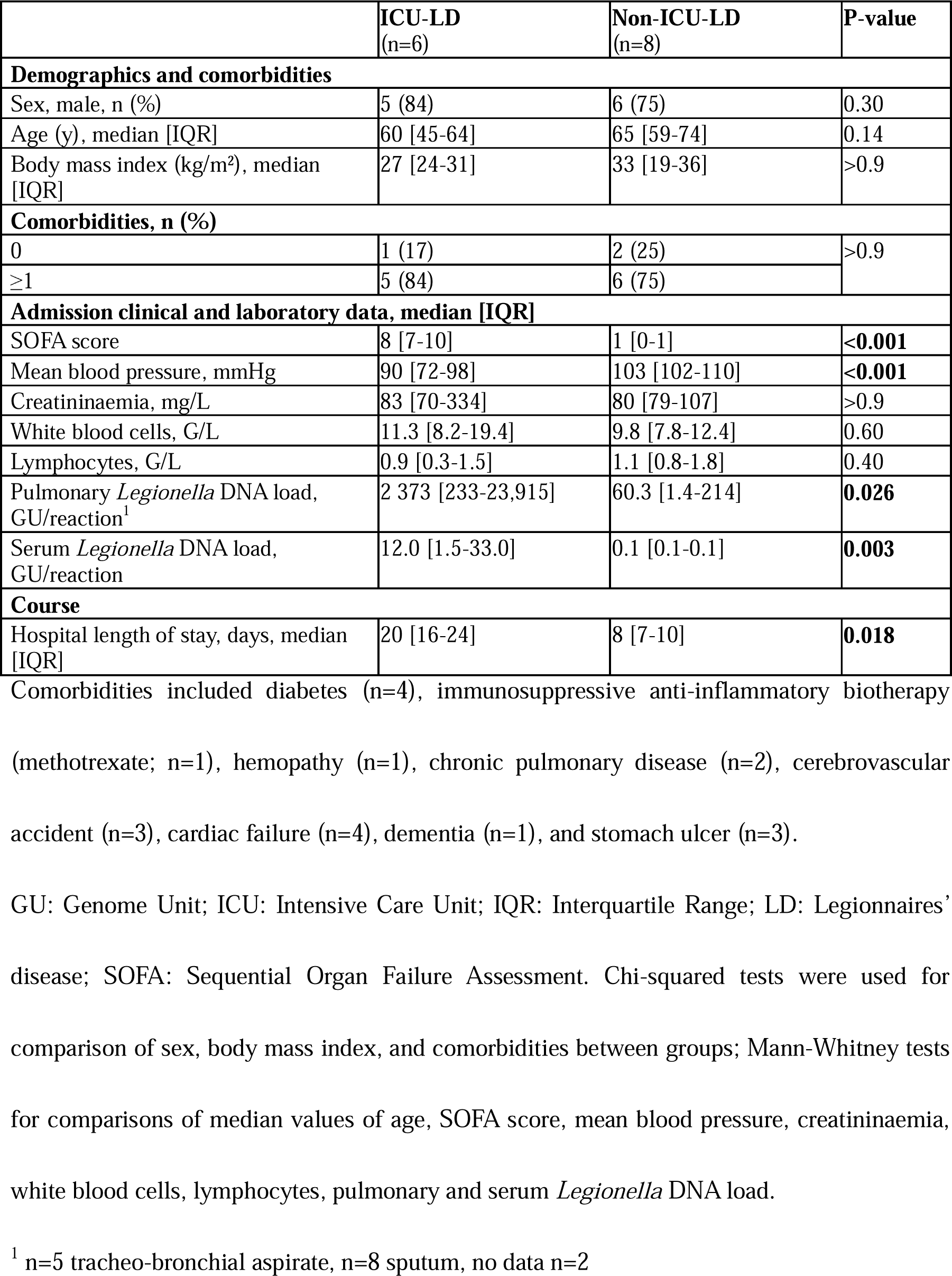
Characteristics of ICU-LD, and non-ICU-LD patients included in the study.

### ICU-LD and non-ICU-LD patients display post-stimulation immune alterations, which are more pronounced in ICU-LD patients

We first assessed the DEGs among the 93 immune gene panel in ICU-LD and non-ICU-LD patients in comparison with HVs (n=9) after LPS stimulation. Among ICU-LD patients, 35/93 (38%) genes were less-expressed while 3/93 (3%; *ADGRE3*, *CX3CR1*, and *IL18R1*) were more-expressed (Figure 1A, Table S3). *IL12B*, *CXCL10*, *IFNG*, *CCL8*, *IDOI1*, *SFAMF7*, and *CXCL9* were the lowest expressed genes by ICU-LD patients, (log2(FC) range: -5.3 to -3.0), five of which were also the least expressed genes by non-ICU-LD patients (Figure 1B). ICU-LD patients had 1.6-fold greater number of less-expressed genes than non-ICU-LD patients (35/93 *vs* 22/93, p=0.039; Figure 1C), and 18/39 were common (Figure 1D, Figure 1 near here). Median gene expression of these 39 less-expressed genes was lower in ICU-LD compared to non-ICU-LD patients (Log2(FC) -1.9 [-2.6;-1.5] *vs* -1.2 [-1.7;-0.9], p=0.0011; Figure 2A and B). When considering individual patients’ gene expression, 7/39 DEG were significantly less expressed by ICU-LD patients (*IRF7*, *MX1*, *NFKBI2*, *NFKBIA*, *RELB*, *SRC*, *TIM3*, p-value range: 0.029 to 0.0080, Figure 2C, Figure 2 near here).

**Figure 1:**
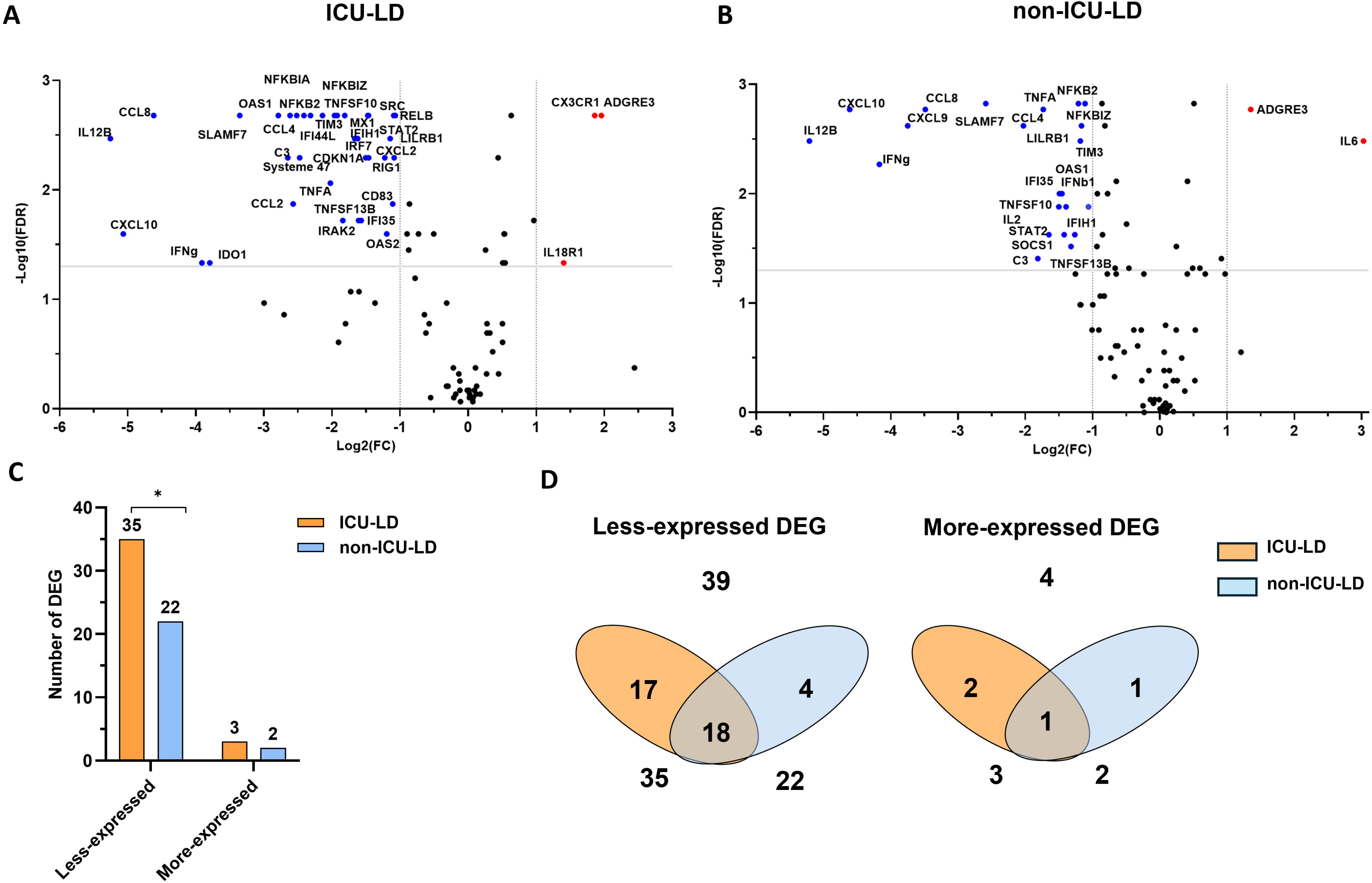
DEGs in ICU-LD and non-ICU-LD patients compared to HVs after LPS stimulation. Volcano plots of the immune gene panel (n=93) in ICU-LD (A) and non-ICU-LD (B) patients against gene expression in HVs. Differential expression (Log2(FC), x axis) is plotted against statistical significance (-log(10)FDR, y axis) for each gene. Black circles represent genes not found to differ significantly between the HVs and LD populations. DEGs with a FC≥2 (log2(FC) ≥1) and a FDR <0.05 (-log10(FDR)>1.33) are in red; DEGs with a FC≥2 and a FDR <0.05 are in blue. Comparison of the number of less-and more-expressed DEGs in ICU-LD and non-ICU-LD patients (C), * p-value of Chi-squared test <0.05. Venn diagrams of less-expressed and more-expressed DEGs in ICU-LD and non-ICU-LD patients (D)

**Figure 2:**
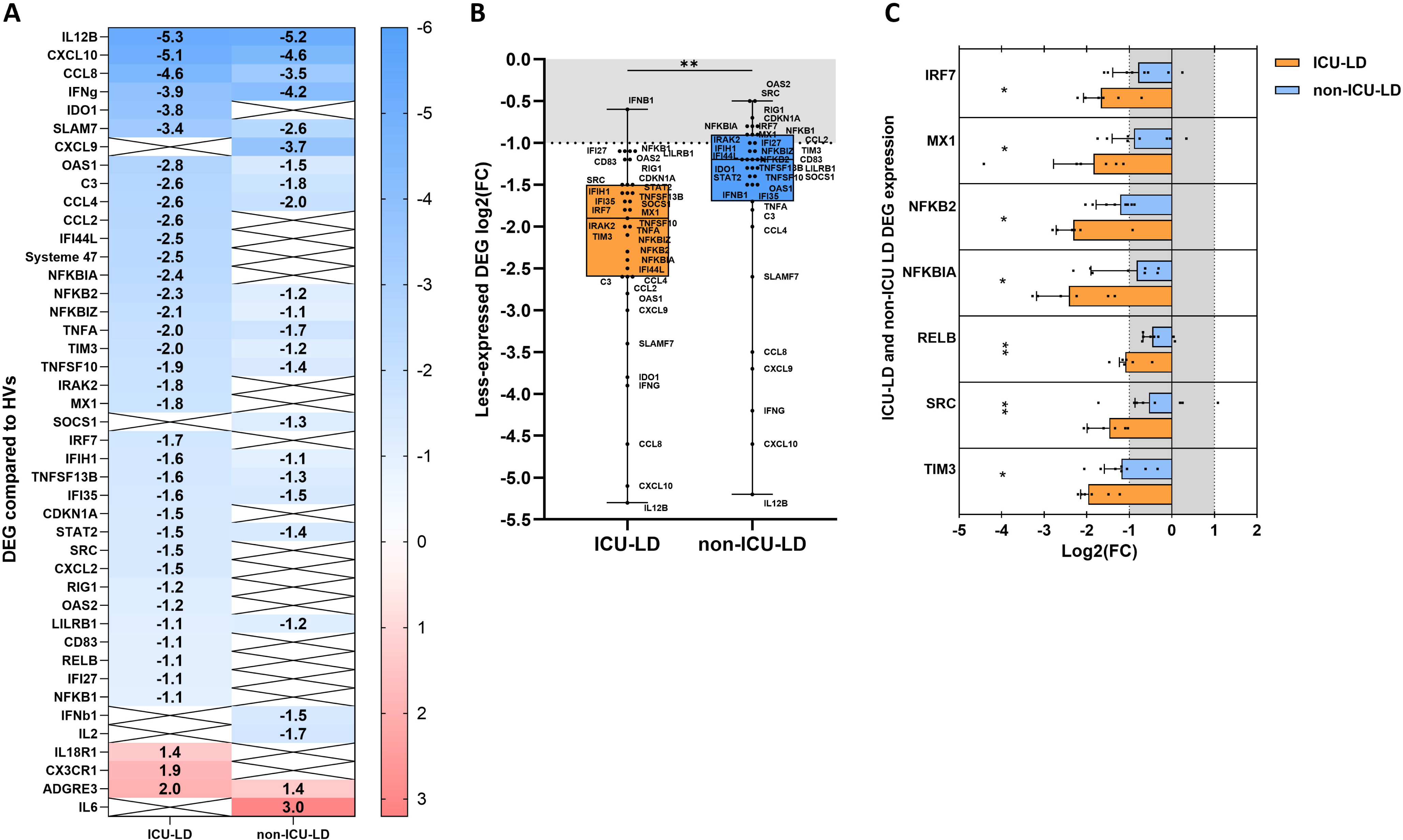
Comparison of DEG between ICU-LD and non-ICU-LD patients after LPS stimulation. List of less- and more-expressed DEGs in ICU-LD and non-ICU-LD patients (A). The blue to red colour gradient indicates the lowest to highest log2(FC) values. White boxes with black crosses indicate genes that were not differentially expressed between non-ICU-LD patients and HVs or ICU-LD patients and HVs. Boxplots presenting the median, IQR, and range gene expressions of the 39 less-expressed genes in ICU-LD compared with non-ICU-LD patients (B). Histogram presenting the 7 significantly less-expressed genes in ICU-LD compared with non-ICU-LD patients, representing median, first quartile, and individual gene expression (C). Mann-Whitney tests were used to compare median values and p-values were adjusted for multiple comparisons using Benjamini-Hochberg correction. * p-adjusted <0.05, ** p-value adjusted <0.01. The grey areas figure the log2(FC) between -1 and 1 (considered as non-significant when comparing to HVs expression).

We then compared the functional alterations found in ICU-LD patients to that of patients with a LD-unrelated-SS. There was no significant difference in the number of less-expressed genes between ICU-LD and LD-unrelated-SS patients (35/93 *vs* 42/93, p=0.30; Figure S1A and Table S3), 33/44 genes were common to both populations (Figure S1B and C), nor in their median expression (Log2(FC) -2.0 [-2.9;-1.6] *vs* -1.9 [-2.6;-1.5], p=0.65; Figure S1D).

### Cellular response to LPS, NF-*κ*B, and type I IFN pathways are more dysregulated in ICU-LD patients

A total of 733 terms and pathways were enriched in ICU-LD and 538 in non-ICU-LD patients from the previously identified less-expressed genes (ICU-LD, n=35, Table S.2 and non-ICU-LD: n=22, Table S.3). We then analysed the top five GO biological processes, subcellular localisation terms, and reactome pathways enriched in ICU-LD (Figure 3A) and the top five enriched in non-ICU-LD patients (Figure 3B) based on their signals. Of these enriched terms or pathways, 10 were common to both ICU-LD and non-ICU-LD patients (Figure 3C, Figure 3 near here). Signals and count of observed genes were greater for 8/10 in ICU-LD compared with non-ICU LD, including cytokine-mediated signalling pathway, defense response and response to virus, (Figure 4A), NF-κB complex (Figure 4B), as well as cytokine signalling in immune system, IFN α/β signalling, IL-10 signalling, and signalling by ILs (Figure 4C, Figure 4 near here). The five unique enriched terms in non-ICU-LD patients included the regulation of adaptive and innate immune response, IgG immunoglobulin and TGF-β complex, and IFN signalling pathway. The five enriched terms and pathways unique to ICU-LD encompassed 11/17 genes uniquely less-expressed in ICU-LD patients (Figure 2A) and 6/7 genes with reduced expression in ICU-LD patients (Figure 2C): cellular response to LPS (*CCL2*, *NFKBIA*, *IRAK2*, *TIM3, SRC* and *NFKB1*), regulation of IFN-β production (*IRF7*, *RIG1*, *OAS2*, and *RELB*), I-κB/NF-κB complex (*NFKBIA*, *NFKB1*, and *NFKB2*), IFN regulatory factor complex (*RIG1* and *IRF7*), and TRAF6-mediated NF-κB activation pathway (*NFKBIA*, *NFKB1*, *NFKB2*, and *RIG1*).

**Figure 3:**
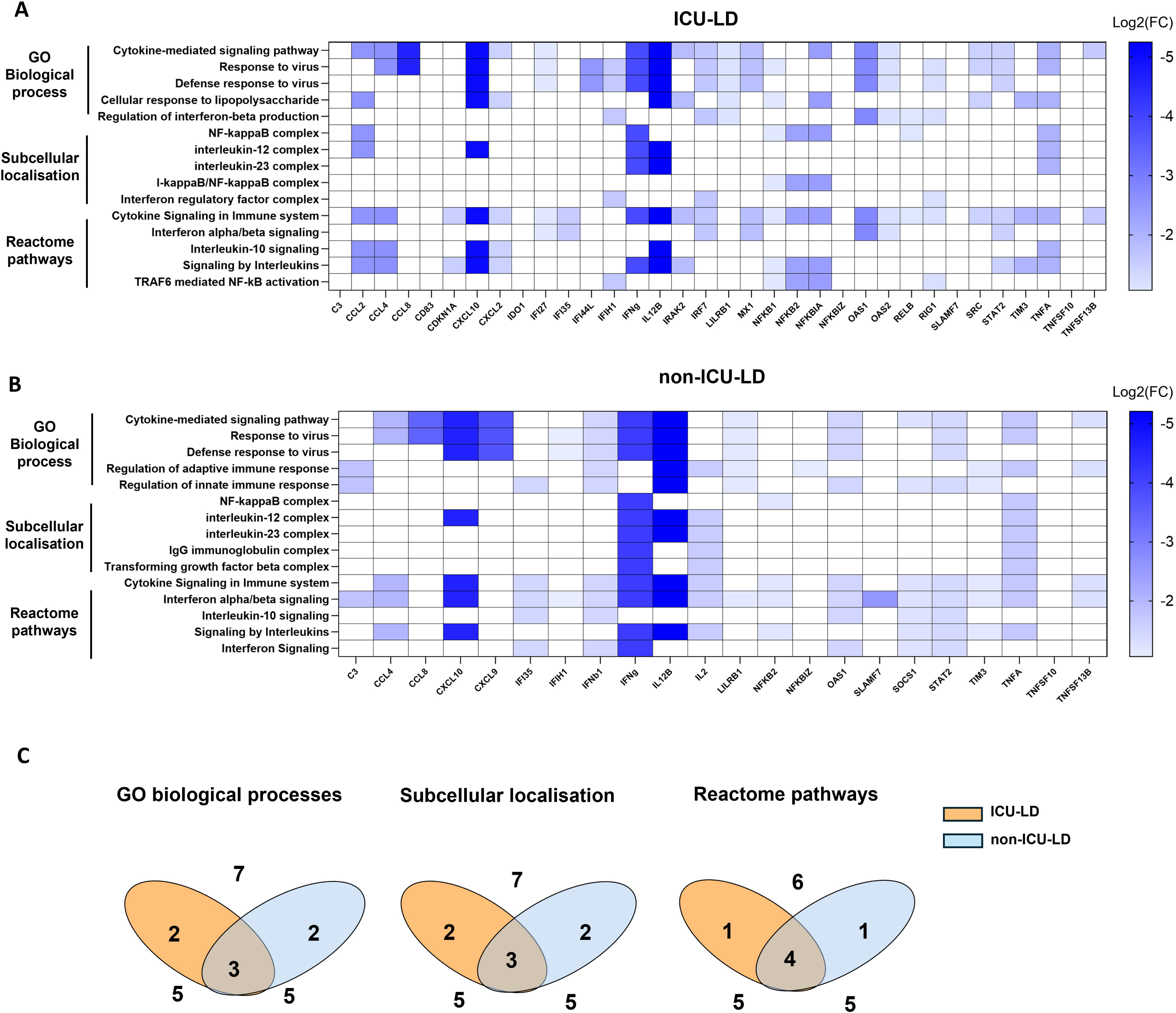
Top five enriched terms and pathways and corresponding genes in ICU-LD and non-ICU-LD. Top five enriched GO biological process, subcellular localisation terms, and reactome pathways in ICU-LD (A) and non-ICU-LD (B) patients according to the gene expression (log2(FC) of less-expressed genes found in these enriched terms and pathways. White boxes indicate genes that are not observed in the enriched terms or pathways. One DEG being a human endogenous retrovirus (System47) was not in the STRING database and was not counted in this analysis. Venn diagram of the top five enriched terms and pathways in ICU-LD and non-ICU-LD patients (C).

**Figure 4:**
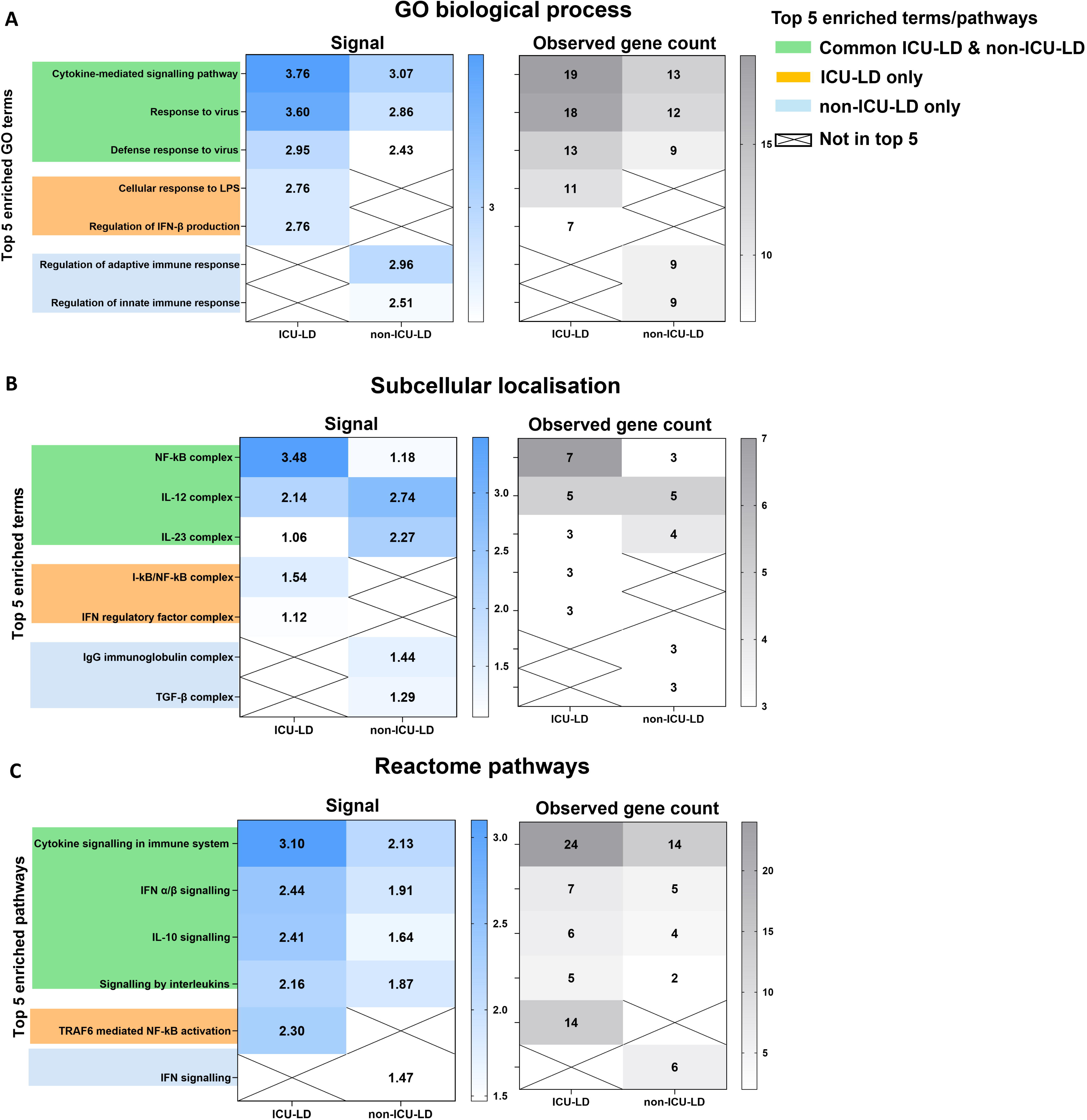
Top five enriched terms and pathways comparison in ICU-LD with non-ICU-LD. Signals and observed gene count of the top five enriched GO biological process (A), subcellular localisation (B) terms, and reactome pathways (C) in ICU-LD and non-ICU-LD patients. One DEG being a human endogenous retrovirus (System47) was not in the STRING database and was not counted in this analysis.

## Discussion

The present study reveals significant immune gene expression alterations in patients with LD following whole blood *ex vivo* LPS stimulation. ICU-LD patients had a greater number of less-expressed genes, a lower expression of these, and a greater number of associated enriched terms and pathways than non-ICU-LD patients. This suggests that ICU-LD display deeper functional immune alterations, that may resemble that of SS.

In ICU-LD patients, LPS stimulation produced a lower expression of more than a third of the selected gene panel compared to HVs while only a few were more expressed. Among them, *IL12B*, *CXCL10*, and *IFNG* from the IFN-γ family, *CCL8* and *CXCL9* coding for chemokines, *IDOI1* coding for indoleamine 2,3-dioxygenase 1, and *SFAMF7* that is involved in the activation of immune cells were the lowest expressed genes (Log2(FC) <-3). Non-ICU-LD patients also displayed altered gene expression, indicating that both ICU and non-ICU-LD patients’ immune cells are less able to respond to the LPS *ex vivo* stimulus than a healthy population. However, the intensity of these functional alterations was associated with disease severity, as ICU-LD patients had 1.6-fold more less-expressed genes and a lower median expression of these compared to non-ICU patients. In particular, seven genes (*IRF7*, *MX1*, *NFKBI2*, *NFKBIA*, *RELB*, *SRC*, and *TIM3*) were significantly less expressed by ICU-patients. ICU-LD patients also had higher *Legionella* DNA loads in the lung and serum, although the causal relationship between immune alterations and bacterial DNA load needs to be further explored.

Among the top five enriched terms and pathways, half were common to both ICU-LD and non-ICU-LD patients, and among these the signals and gene counts were greater in ICU-LD patients, probably explained by the greater gene expression alteration. It is of note that the dysregulated pathways common to both ICU-LD and non-ICU-LD patients belong to critical processes for controlling *Legionella* invasion and replication. Among these, cytokine mediated signalling pathways, including TNF-α (*TNF*, *TNFSF13B*, and *TNFS10*), IFN-γ (*IFNG*, *IL12B*, and *CXCL10*), and chemokines (*CCL2*, *CCL4,* and *CCL8*) families, and genes from the NF-κB complex (*NFKBIA, NFKB1, NFKB2, RELB, CCL2, IFNG,* and *TNF*), all strongly activated in response to LPS, play a critical role in *Legionella* clearance [17]. Response to virus pathways may be common to pathways controlling anti-intracellular bacteria multiplication, such as *Legionella* [18]. Interestingly, MX dynamin-like GTPase 1 (*MX1*), belonging to cytokine signalling and defense to virus pathways, has recently been identified as an effector molecule involved in microRNA-dependent control of *L. pneumophila* replication within human macrophages. Its downregulation enhances *L. pneumophila* replication [19]. In addition, type I IFNs (IFN-α and IFN-β) have been reported to enhance host defences against *Legionella* by limiting intravacuolar multiplication in a mouse model [17].

ICU-LD patients also exhibited five uniquely enriched terms or pathways related to anti-*Legionella* response, encompassing most of the genes uniquely less-expressed in this group. Among them, cellular response to LPS and I-κB/NF-κB complex pathways are closely related to dysregulated pathways common to both ICU-LD and non-ICU-LD patients. Regulation of IFN-β production and IFN regulatory factor complex pathways include *IRF7* coding interferon regulatory factor 7 (IRF-7), and *RIG1* coding RIG-I receptor. While IRF-7 has recently been shown to drive autophagy and to eliminate intra-macrophage pathogens during sepsis, ultimately improving the outcome of sepsis in mice [21], its role in the intracellular replication of *Legionella* has never been described. RIG-I participates in an immune network restricting *Legionella* replication in human macrophages [22]. IFN-β production pathway also included *OAS2* gene coding one of the 2’-5’-oligoadenylate synthetases involved in the restriction of intracellular pathogenic mycobacterial replication [23]. Lastly, the TRAF6-mediated NF-κB activation pathway can be activated by pleiotropic cellular signals important in *Legionella* infection, including the binding of membrane toll-like receptors (TLR) 2 and 5, and intracellular TLR 9 [17,24]. It has been shown that intracellular bacteria such as *Francisella* [25] and *Shigella* [26] inhibit TRAF6-mediated signalling as a strategy to proliferate within the host by limiting the immune response. *M. tuberculosis* also inhibits this pathway to reduce autophagy [27]. The decreased TRAF6 pathway observed in ICU-LD patients could therefore contribute to increased *Legionella* replication in these individuals, perpetuating the severity of the infection.

To our knowledge, such an extended evaluation of immune functionality in LD (ICU and non-ICU) patients has yet to be published. The present findings also confirm our previous observations that ICU-LD patients exhibit lower release of pro-inflammatory cytokines, TH-1 and TH-2 cytokines, chemokines, and growth factors than HVs after whole blood stimulation, and aligns with results of a TNF-α expression alteration after LPS stimulation in LD patients requiring prolonged MV [11]. Furthermore, *IL18R1* was more greatly expressed in ICU-LD patients after LPS stimulation while *IL18* and *IL1B* were not differentially expressed. IL-1β and IL-18 are both NRLP3 inflammasome terminal products that are essential for the control of *Legionella* by promoting cell pyroptosis in a mouse model [28]. These results, together with our previous study reporting increased IL-18 in the supernatant of stimulated blood cells from ICU-LD patients [11], suggest partial preservation of inflammasome activation in human LD.

The immune response of patients with SS is known to be deeply impaired, and associated with secondary nosocomial infections and Day-28 mortality [29]; in particular, SS immune cells fail to respond to LPS stimulus (endotoxin tolerance) [12]. Post-infection altered immune response has also been found in ICU-COVID-19 patients [30], and patients admitted to a general ward for community-acquired pneumonia [31,32]. Frequently found features of endotoxin tolerance in SS, including low expression of *TNFA* [33], *IFNG* [34], *IL12*, and *IL1A* [35], were observed in both ICU-LD and LD-unrelated-SS patients herein. To restore immune function, immunomodulatory drugs have been tested in clinical trials or in *in vitro* assays in the context of SS (GM-CSF, IL-7, anti-PD1) as well as severe COVID-19 (IL-7) [12]. Some of these have shown promising results in lymphopenia restoration or cytokine storm reduction. Given the post-infectious immunosuppression described herein, such immunotherapy may also be beneficial for ICU-LD patients.

The present study does have several limitations. First, the small and heterogeneous patient groups limit the generalisability, which should be validated in a larger, independent cohort. The gene panel design was restricted, and we could not include important markers such as *C5AR1* (coding for CD88, for neutrophils) or lymphocyte exhaustion markers (*PD1* coding for CD279, or *CTLA4* coding for CD152), known to be dysregulated in SS [12]. Additionally, the panel included multiple genes from the same families (*e.g.* NF-κB, IFN-γ), that vary in the same way after LPS stimulation, which may have amplified the observed results; a transcriptomic response assessment could address this bias.

In conclusion, immune gene expression alterations in LD after LPS stimulation were found herein, with more pronounced alterations in ICU-LD patients. These findings suggest that reduced expression of key genes and pathways involved in controlling *Legionella* proliferation in ICU-LD patients may contribute to increased disease severity.

## Acknowledgements

We thank physicians from university hospitals that included patients in the ProgLegio study: F. Ader, L. Argaud, T. Baudry, N. Freymond, A. Frigerri, Y. Jamilloux, V. Labeye, J.C. Richard, (Lyon, infectious disease, emergency, pneumology and hematology departments, ICU), J.C. Navellou (Besançon, ICU), S. Alfantari, L. Thirard, P.Y. Delanoye, O. Robineau (Tourcoing, ICU), F.X. Blanc (Nantes, Pneumology), S. Nseir (Lille, ICU), V. Lemiale, J.F. Timsit, M.S. Fartoukh (ICU, Assistance Publique Hôpitaux de Paris), R. Le Berre (Pneumology and Internal medicine department, Brest), S. Perinel-Ragey (ICU, Saint-Etienne), J.P. Quenot (ICU, Dijon), A. Gacouin (ICU, Rennes). The authors would like to thank technicians from the national reference centre (J. Chastang, L. Chaverot, A. Marie, J. Reboulet, I. Royet, and M. Siffert) for technical support. We are grateful to P. Robinson (DRS, *Hospices Civils de Lyon*) for help in manuscript preparation. We acknowledge the contribution of the EFS—Rhône-Alpes.

## Contributors

The project was conceived, planned, and supervised by SJ and STA. SJ, FA, CA, GD, LB, MI, PD, JG, AC, and GL were involved in the design, implementation, and day-to-day management of the study. WM and STA were involved in the IFA selection and analysis of the results. CA-V brought post-LPS stimulation HVs and SS data. J-CR and FV brought their expertise about clinical and laboratory data interpretation. ZM brought his expertise for immunological gene expression interpretation. LA, J-CR, AF, VL, YJ, NF, and FA included patients in the study. CA, GD, LB, MI, and SJ were responsible for the microbiological analyses; CA and WM were responsible for the immunological analyses. CA and WM were involved in statistical analyses. CA and SJ wrote the original draft of the manuscript, which was reviewed and edited by WM, ST-A, FV, MI, CG, GD, LB, ACL, GM, PD, AC, and FA and reviewed by all co-authors. All authors approve the final version of the manuscript.

## Data availability statement

The datasets analysed in the present study are available on the main manuscript or in the supplement material. Raw gene expression level is available upon reasonable request to the corresponding author.

