## Supplementary Methods for "Functional alterations of immune gene expression in ICU and non-ICU patients with Legionnaires’ disease, a prospective observational study"

***ProgLegio study***

The inclusion criteria of the ProgLegio national prospective cohort study were: (i) patients with clinical and laboratory signs of LD (positive urinary antigen test and/or *L. pneumophila* PCR on respiratory sample for *L. pneumophila serogroup 1* only), and (ii) having provided written informed consent (legal representative could be used as a surrogate). Exclusion criteria were: (i) LD caused by *Legionella* non *pneumophila* or *L. pneumophila* serogroup non-1; (ii) patients for whom respiratory secretions could not be obtained, (iii) cases diagnosed only by serology; (iv) outpatients. Age and immunosuppression status (IS; e.g. long-term corticosteroids, immunosuppressive therapy, including anti-TNF-α and other biotherapies, active solid cancer or hemopathy, and other diseases inducing immunosuppression) were collected for all LD patients but were not considered as exclusion criteria. This study was approved by the regional institutional review board (*Comité de Protection des Personnes Sud-Est IV, France*; ID-RCB 2016-A01021-50). It was registered with the *Ministère de lʼEnseignement supérieur, de la Recherche et de lʼInnovation* (DC-2008–509) and the French data protection agency (*Commission nationale de l’informatique et des libertés*).

***Immunosepsis-4 study***

LD-unrleated-SS patients were included in Immunosepsis-4 prospective study that investigated ICU-induced immune dysfunctions. They were included based on sepsis-3 definition of septic shock: vasopressor requirement and serum lactate >2mmol/L in the absence of hypovolemia in a patient with suspected or proven infection. The exclusion criteria were age <18 years and the presence of aplasia or known immuno-suppressive disease . was approved by the regional ethics committee (*Comité de Protection des Personnes Sud-Est II*, number 11236), which waived the need for written informed consent owing to the observational nature of the study, with a low risk for patients, and no specific procedure was required other than routine blood sampling. Oral information and non-opposition to inclusion in the study were mandatory and were systematically obtained before any blood sample was drawn. This was recorded in the patients’ clinical file. This study was also registered at the French ministry of research (*Ministère de lʼEnseignement supérieur, de la Recherche et de lʼInnovation*, DC-2008–509) and with the national data protection commission (*Commission Nationale de l’Informatique et des Libertés*). The study protocol was designed and conducted in accordance with the Declaration of Helsinki54 and Good Clinical Practice.

***Pulmonary and serum Legionella DNA load***

Pulmonary and serum *Legionella* DNA loads were measured from thawed samples (sputum or tracheal aspirates) taken on the day of enrolment in the ProgLegio study. DNA was extracted from 200 µL of pulmonary, or 180 µL of serum samples, using the MagNA Pure Compact Instrument (Roche Diagnostics, Basel, Switzerland) automated system. *Legionella* DNA was assessed using a 5 µL sample by a quantitative PCR (qPCR) targeting *mip* [31] and the DNA load was quantified using a calibration range based on a *Legionella* DNA standard reference material [32]. Results were expressed in Genome Units (GU) per reaction.

***mHLA-DR measurement***

Expression of surface mHLA-DR was measured by flow cytometry from EDTA samples collected within 2 hours at Day-2 (ICU-LD patients) or Day-3 (SS patients). The anti-HLA-DR/Anti-Monocyte Quantibrite assay (BD Biosciences, San Jose, CA, USA) was performed on a Navios flow cytometer and data were analyzed using Navios software (NAVIOS; Beckman-Coulter, Brea, CA, USA). Monocytes were gated based on CD14 expression. mHLA‐DR expression was measured as the median of fluorescence intensity related to the entire monocyte population, as recommended by the manufacturer. The fluorescence was converted to antibodies bound per cell (Ab/C) using a calibrated standard curve determined with phycoerythrin (PE)‐beads (BD QuantiBrite™ ‐ PE Beads, BD). Results are expressed as number of sites per cell (Ab/C). The usual values are 13 500-45 000 Ab/C and the threshold value for immunosuppression is <8000 Ab/C [33].

***mRNA extraction***

Total RNA was extracted from thawed whole blood buffy coat preserved in Trizol (ThermoFisher Scientific, Waltham, MA, USA) using an on-column method (NucleoSpin miRNA; Macherey-Nagel Gmbh&Co. KG, Düren, Germany), quantified using a Qubit 2.0 Fluorometer (ThermoFisher scientific), and normalised to 200 ng.

***Immune functional assays (IFA)***

LPS was chosen because it induces the most robust and complementary activation of innate and adaptive immunity in healthy individuals[13]. In parallel, a negative control (NUL condition) containing only culture medium (Gibco RPMI 1640 medium, Fisher Scientific SAS, Illkirch-Graffenstaden, France) was carried out for all patient and HV samples.

***nanoString process, data analysis, and normalisation***

Each mRNA extract (see list in Table S1) was analysed in a separate multiplexed reaction each including 8 negative probes and 6 serial concentrations of positive control probes. Data were imported into nSolver analysis software (version 4.0, nanoString Technologies) for quality checking and normalisation of the data. A first normalisation step using the internal negative and positive controls allowed correction of a potential source of variation associated with the technical platform. To do so, we calculated for all the samples the background level as the median +3 standard deviations across the six negative probe counts. Each sample under the background level was fixed to this value. Next, we calculated for each sample the geometric mean of the positive probe counts. A scaling factor for a sample was a ratio of the geometric mean of the sample and the average across all geometric means. For each sample, we divided all gene counts by the corresponding scaling factor. To normalise for differences in RNA input we used the same method as in the positive control normalisation, except that geometric means were calculated over three housekeeping genes (*HPRT1* (NM_000194.1), *DECR1* (NM_001359.1), and *TBP* (NM_001172085.1)). These genes were selected using the NormFinder method, an established approach for identification of stable housekeeping genes within and between groups, from the 6 candidate genes included in the custom panel (*HPRT1*, *DECR1*, *TBP*, *GAPDH*, *PLOR2A*, and *RPL19*).

***STRING database analysis***

We entered the list of proteins coded by DEGs and their respective gene expression FC in the section “Protein with Values/Ranks”. The organism “Homo sapiens” was chosen. One gene from the panel (*SYSTEM47*) were not in the database as they are human endogenous retroviruses. In the analysis section, the top five results of each GO process, subcellular localisation terms, and reactome pathways according to their signal. The signal is defined as a weighted harmonic mean between the observed/expected ratio and -log(FDR). FDR tends to emphasise larger terms due to their potential for achieving lower p-values, while the observed/expected ratio highlights smaller terms, which have a high foreground to background ratio but cannot achieve low FDR values due to their size. The signal measure seeks to balance both metrics for more intuitive ordering of enriched terms.
