## Supplementary material for "Functional alterations of immune gene expression in ICU and non-ICU patients with Legionnaires’ disease, a prospective observational study": Table S1

**Supplementary results**

**Table S1: nanoString nCounter mRNA panel**

| Gene name | nanoString identifier | Accession number | Target sequence |
| --- | --- | --- | --- |
| *ADGRE3* | NM_032571.2:1015 | NM_032571.2 | TGTATCTGAACTCTCAGGTTGTGAGTGCTGCTATTGGACCCAAAAGGAACGTGTCTCTCTCCAAGTCTGTGACGCTGACTTTCCAGCACGTGAAGATGAC |
| *ARL14EP* | NM_152316.1:1650 | NM_152316.1 | AGAAGAAGTCACTGACCATGGGAATGTTGTTCTTGCTGCTGTGTATTCATAGGAGCTTAGTGAAGGCAAACTTACCAACACAAATAAGCAAAGTGGTTGC |
| *BST2* | NM_004335.2:560 | NM_004335.2 | GAAGCTGGCACATCTTGGAAGGTCCGTCCTGCTCGGCTTTTCGCTTGAACATTCCCTTGATCTCATCAGTTCTGAGCGGGTCATGGGGCAACACGGTTAG |
| *C3* | NM_000064.2:4396 | NM_000064.2 | CATCTACCTGGACAAGGTCTCACACTCTGAGGATGACTGTCTAGCTTTCAAAGTTCACCAATACTTTAATGTAGAGCTTATCCAGCCTGGAGCAGTCAAG |
| *CCL2/MCP1* | NM_002982.3:123 | NM_002982.3 | CATTCCCCAAGGGCTCGCTCAGCCAGATGCAATCAATGCCCCAGTCACCTGCTGTTATAACTTCACCAATAGGAAGATCTCAGTGCAGAGGCTCGCGAGC |
| *CCL20* | NM_004591.1:35 | NM_004591.1 | ATCTGTTCTTTGAGCTAAAAACCATGTGCTGTACCAAGAGTTTGCTCCTGGCTGCTTTGATGTCAGTGCTGCTACTCCACCTCTGCGGCGAATCAGAAGC |
| *CCL4* | NM_002984.2:35 | NM_002984.2 | TTCTGCAGCCTCACCTCTGAGAAAACCTCTTTGCCACCAATACCATGAAGCTCTGCGTGACTGTCCTGTCTCTCCTCATGCTAGTAGCTGCCTTCTGCTC |
| *CCL8* | NM_005623.2:689 | NM_005623.2 | AAGGAGAGATGGGTCAGGGATTCCATGAAGCATCTGGACCAAATATTTCAAAATCTGAAGCCATGAGCCTTCATACATGGACTGAGAGTCAGAGCTTGAA |
| *CCNB1IP1* | NM_182849.2:792 | NM_182849.2 | TCGTCAGTATCAAAAGCTCCAAGGCCTCTATGATAGCCTTAGGCTACGAAACATCACTATTGCTAACCATGAAGGCACCCTTGAACCATCCATGATTGCA |
| *CCR1/ RANTES R* | NM_001295.2:535 | NM_001295.2 | CATCATTTGGGCCCTGGCCATCTTGGCTTCCATGCCAGGCTTATACTTTTCCAAGACCCAATGGGAATTCACTCACCACACCTGCAGCCTTCACTTTCCT |
| *CD127/IL7R* | NM_002185.2:1610 | NM_002185.2 | TTGCTTTGACCACTCTTCCTGAGTTCAGTGGCACTCAACATGAGTCAAGAGCATCCTGCTTCTACCATGTGGATTTGGTCACAAGGTTTAAGGTGACCCA |
| *CD209/DC-SIGN* | NM_021155.2:1532 | NM_021155.2 | TTATCTCATACATGCAAACCTACCATCTGTTCAACTTCCACCTACCACCTCCTGCACCCCTTTGATCGGGGACTTACTGGTTGCAAGAGCTCATTTTGCA |
| *CD3D* | NM_000732.4:110 | NM_000732.4 | TATCTACTGGATGAGTTCCGCTGGGAGATGGAACATAGCACGTTTCTCTCTGGCCTGGTACTGGCTACCCTTCTCTCGCAAGTGAGCCCCTTCAAGATAC |
| *CD44* | NM_001001392.1:429 | NM_001001392.1 | ACACCATGGACAAGTTTTGGTGGCACGCAGCCTGGGGACTCTGCCTCGTGCCGCTGAGCCTGGCGCAGATCGATTTGAATATAACCTGCCGCTTTGCAGG |
| *CD74* | NM_001025159.1:964 | NM_001025159.1 | TTCAGCCCCCAGCCCCTCCCCCATCTCCCACCCTGTACCTCATCCCATGAGACCCTGGTGCCTGGCTCTTTCGTCACCCTTGGACAAGACAAACCAAGTC |
| *CD83* | NM_004233.3:1960 | NM_004233.3 | CTGTTCTTGAAGCAGTAGCCTAACACACTCCAAGATATGGACACACGGGAGCCGCTGGCAGAAGGGACTTCACGAAGTGTTGCATGGATGTTTTAGCCAT |
| *CDKN1A* | NM_000389.2:1975 | NM_000389.2 | CATGTGTCCTGGTTCCCGTTTCTCCACCTAGACTGTAAACCTCTCGAGGGCAGGGACCACACCCTGTACTGTTCTGTGTCTTTCACAGCTCCTCCCACAA |
| *CLEC7A/DECTIN-1* | NM_197954.2:55 | NM_197954.2 | TGTTAAACTCCGGTAAGTACCTAGCCCACATGATTTGACTCAGAGATTCTCTTTTGTCCACAGACAGTCATCTCAGGAGCAGAAAGAAAAGAGCTCCCAA |
| *CX3CR1* | NM_001337.3:1040 | NM_001337.3 | GGGCGCTCAGTCCACGTTGATTTCTCCTCATCTGAATCACAAAGGAGCAGGCATGGAAGTGTTCTGAGCAGCAATTTTACTTACCACACGAGTGATGGAG |
| *CXCL10/IP10* | NM_001565.1:40 | NM_001565.1 | GCAGAGGAACCTCCAGTCTCAGCACCATGAATCAAACTGCGATTCTGATTTGCTGCCTTATCTTTCTGACTCTAAGTGGCATTCAAGGAGTACCTCTCTC |
| *CXCL2/MIP2alpha* | NM_002089.3:854 | NM_002089.3 | ATCACATGTCAGCCACTGTGATAGAGGCTGAGGAATCCAAGAAAATGGCCAGTGAGATCAATGTGACGGCAGGGAAATGTATGTGTGTCTATTTTGTAAC |
| *CXCL9* | NM_002416.1:1975 | NM_002416.1 | CACCATCTCCCATGAAGAAAGGGAACGGTGAAGTACTAAGCGCTAGAGGAAGCAGCCAAGTCGGTTAGTGGAAGCATGATTGGTGCCCAGTTAGCCTCTG |
| *DECR1* | NM_001359.1:835 | NM_001359.1 | GAATGCGATTCAATGTGATTCAACCAGGGCCTATAAAAACCAAAGGTGCCTTTAGCCGTCTGGACCCAACTGGAACATTTGAGAAAGAAATGATTGGCAG |
| *DYRK2* | NM_003583.3:1922 | NM_003583.3 | CCGGTGCTATCACATCTATATCCAAGTTACCTCCACCTTCTAGCTCAGCTTCCAAACTGAGGACTAATTTGGCGCAGATGACAGATGCCAATGGGAATAT |
| *EIF2AK4* | NM_001013703.2:2715 | NM_001013703.2 | TGATTAAGTCAGACCCTTCAGGTCACTTAACTGGGATGGTTGGCACTGCTCTCTATGTAAGCCCAGAGGTCCAAGGAAGCACCAAATCTGCATACAACCA |
| *FAM89A* | NM_198552.2:750 | NM_198552.2 | GCGGATGCATGGTGGCAGTCTGCTTTGATGGCAGCAGTTTCTGCTTAGGTGACCTAGAGGTCCTCAGCAGTATCCTCCACACCTATTTATTGAGGTGCAC |
| *FOXP3* | NM_014009.3:1230 | NM_014009.3 | GGGCCATCCTGGAGGCTCCAGAGAAGCAGCGGACACTCAATGAGATCTACCACTGGTTCACACGCATGTTTGCCTTCTTCAGAAACCATCCTGCCACCTG |
| *GAPDH* | NM_002046.5:350 | NM_002046.5 | AAATTCCATGGCACCGTCAAGGCTGAGAACGGGAAGCTTGTCATCAATGGAAATCCCATCACCATCTTCCAGGAGCGAGATCCCTCCAAAATCAAGTGGG |
| *GATA3* | NM_001002295.1:2835 | NM_001002295.1 | AAGAGTCCGGCGGCATCTGTCTTGTCCCTATTCCTGCAGCCTGTGCTGAGGGTAGCAGTGTATGAGCTACCAGCGTGCATGTCAGCGACCCTGGCCCGAC |
| *HLA-DMB* | NM_002118.3:20 | NM_002118.3 | CCCGTGAGCTGGAAGGAACAGATTTAATATCTAGGGGCTGGGTATCCCCACATCACTCATTTGGGGGGTCAAGGGACCCGGGCAATATAGTATTCTGCTC |
| *HLA-DPA1* | NM_033554.2:857 | NM_033554.2 | GGAGAGATCTGAACTCCAGCTGCCCTACAAACTCCATCTCAGCTTTTCTTCTCACTTCATGTGAAAACTACTCCAGTGGCTGACTGAATTGCTGACCCTT |
| *HLA-DPB1* | NM_002121.4:931 | NM_002121.4 | TCCAAATTGGATACTGCTGCCAAGAAGTTGCTCTGAAGTCAGTTTCTATCATTCTGCTCTTTGATTCAAAGCACTGTTTCTCTCACTGGGCCTCCAACCA |
| *HLA-DRA* | NM_019111.3:335 | NM_019111.3 | GGCCAACATAGCTGTGGACAAAGCCAACCTGGAAATCATGACAAAGCGCTCCAACTATACTCCGATCACCAATGTACCTCCAGAGGTAACTGTGCTCACG |
| *HPRT1* | NM_000194.1:240 | NM_000194.1 | TGTGATGAAGGAGATGGGAGGCCATCACATTGTAGCCCTCTGTGTGCTCAAGGGGGGCTATAAATTCTTTGCTGACCTGCTGGATTACATCAAAGCACTG |
| *IDO1* | NM_002164.3:50 | NM_002164.3 | CTATTATAAGATGCTCTGAAAACTCTTCAGACACTGAGGGGCACCAGAGGAGCAGACTACAAGAATGGCACACGCTATGGAAAACTCCTGGACAATCAGT |
| *IFI27* | NM_005532.3:390 | NM_005532.3 | TCACTGGGAGCAACTGGACTCTCCGGATTGACCAAGTTCATCCTGGGCTCCATTGGGTCTGCCATTGCGGCTGTCATTGCGAGGTTCTACTAGCTCCCTG |
| *IFI35* | NM_005533.3:415 | NM_005533.3 | TGCCCTCTGCTTGCGGGCTCTGCTCTGATCACCTTTGATGACCCCAAAGTGGCTGAGCAGGTGCTGCAACAAAAGGAGCACACGATCAACATGGAGGAGT |
| *IFI44L* | NM_006820.2:940 | NM_006820.2 | ATCTCTGCCATTTATGTTGTGTGACACTATGGGGCTAGATGGGGCAGAAGGAGCAGGACTGTGCATGGATGACATTCCCCACATCTTAAAAGGTTGTATG |
| *IFIH1/MDA5* | NM_022168.2:185 | NM_022168.2 | GCTTGGGAGAACCCTCTCCCTTCTCTGAGAAAGAAAGATGTCGAATGGGTATTCCACAGACGAGAATTTCCGCTATCTCATCTCGTGCTTCAGGGCCAGG |
| *IFITM1* | NM_003641.3:482 | NM_003641.3 | CCTGTTACTGGTATTCGGCTCTGTGACAGTCTACCATATTATGTTACAGATAATACAGGAAAAACGGGGTTACTAGTAGCCGCCCATAGCCTGCAACCTT |
| *IFNb1* | NM_002176.2:610 | NM_002176.2 | ACAGACTTACAGGTTACCTCCGAAACTGAAGATCTCCTAGCCTGTGCCTCTGGGACTGGACAATTGCTTCAAGCATTCTTCAACCAGCAGATGCTGTTTA |
| *IFNg* | NM_000619.2:970 | NM_000619.2 | ATACTATCCAGTTACTGCCGGTTTGAAAATATGCCTGCAATCTGAGCCAGTGCTTTAATGGCATGTCAGACAGAACTTGAATGTGTCAGGTGACCCTGAT |
| *IL10* | NM_000572.2:230 | NM_000572.2 | AAGGATCAGCTGGACAACTTGTTGTTAAAGGAGTCCTTGCTGGAGGACTTTAAGGGTTACCTGGGTTGCCAAGCCTTGTCTGAGATGATCCAGTTTTACC |
| *IL12B* | NM_002187.2:1435 | NM_002187.2 | GCAAGGCTGCAAGTACATCAGTTTTATGACAATCAGGAAGAATGCAGTGTTCTGATACCAGTGCCATCATACACTTGTGATGGATGGGAACGCAAGAGAT |
| *IL18* | NM_001562.2:48 | NM_001562.2 | GACAGTCAGCAAGGAATTGTCTCCCAGTGCATTTTGCCCTCCTGGCTGCCAACTCTGGCTGCTAAAGCGGCTGCCACCTGCTGCAGTCTACACAGCTTCG |
| *IL18R1* | NM_003855.2:2025 | NM_003855.2 | GAATGAGGGGATTTTAAGTGTCTGAAGAGGCATTTTCTAGGGACCAGTGGGTGACTGAGTAACTGAAATGCTGCTTTCACTCCCTAACACCATGGATCTG |
| *IL1A* | NM_000575.3:1085 | NM_000575.3 | ACTCCATGAAGGCTGCATGGATCAATCTGTGTCTCTGAGTATCTCTGAAACCTCTAAAACATCCAAGCTTACCTTCAAGGAGAGCATGGTGGTAGTAGCA |
| *IL1B* | NM_000576.2:840 | NM_000576.2 | GGGACCAAAGGCGGCCAGGATATAACTGACTTCACCATGCAATTTGTGTCTTCCTAAAGAGAGCTGTACCCAGAGAGTCCTGTGCTGAATGTGGACTCAA |
| *IL1R2* | NM_004633.3:270 | NM_004633.3 | CACCCTTCAGCCTGCGGCACACACAGGGGCTGCCAGAAGCTGCCGGTTTCGTGGGAGGCATTACAAGCGGGAGTTCAGGCTGGAAGGGGAGCCTGTAGCC |
| *IL2* | NM_000586.2:300 | NM_000586.2 | AGGATGCAACTCCTGTCTTGCATTGCACTAAGTCTTGCACTTGTCACAAACAGTGCACCTACTTCAAGTTCTACAAAGAAAACACAGCTACAACTGGAGC |
| *IL6* | NM_000600.1:220 | NM_000600.1 | TGACAAACAAATTCGGTACATCCTCGACGGCATCTCAGCCCTGAGAAAGGAGACATGTAACAAGAGTAACATGTGTGAAAGCAGCAAAGAGGCACTGGCA |
| *IRAK2* | NM_001570.3:1285 | NM_001570.3 | GTGTTGGCCGAGGTCCTCACGGGCATCCCTGCAATGGATAACAACCGAAGCCCGGTTTACCTGAAGGACTTACTCCTCAGTGATATTCCAAGCAGCACCG |
| *IRF3* | NM_001571.5:1303 | NM_001571.5 | TCATGGCCCCAGGACCAGCCGTGGACCAAGAGGCTCGTGATGGTCAAGGTTGTGCCCACGTGCCTCAGGGCCTTGGTAGAAATGGCCCGGGTAGGGGGTG |
| *IRF7* | NM_001572.3:1763 | NM_001572.3 | CGCAGCGTGAGGGTGTGTCTTCCCTGGATAGCAGCAGCCTCAGCCTCTGCCTGTCCAGCGCCAACAGCCTCTATGACGACATCGAGTGCTTCCTTATGGA |
| *JAK2* | NM_004972.2:455 | NM_004972.2 | CTCCTCCCGCGACGGCAAATGTTCTGAAAAAGACTCTGCATGGGAATGGCCTGCCTTACGATGACAGAAATGGAGGGAACATCCACCTCTTCTATATATC |
| *LILRB1* | NM_001081637.1:2332 | NM_001081637.1 | AGCTGAGAAAACTAAGTCAGAAAGTGCATTAAACTGAATCACAATGTAAATATTACACATCAAGCGATGAAACTGGAAAACTACAAGCCACGAATGAATG |
| *MDC1* | NM_014641.2:6719 | NM_014641.2 | TACCCTTTTCCCTCCCAGACCACGAATTAGAAGATATGTGGAAGAAAGAACTCAGGGCGTTAGAAAGGATTGGGGTATATTGATACAACTTGTCCTGGAA |
| *MERTK* | NM_006343.2:665 | NM_006343.2 | GAAGAGATCGTGTCTGATCCCATCTACATCGAAGTACAAGGACTTCCTCACTTTACTAAGCAGCCTGAGAGCATGAATGTCACCAGAAACACAGCCTTCA |
| *MX1* | NM_002462.2:1485 | NM_002462.2 | GCCTTTAATCAGGACATCACTGCTCTCATGCAAGGAGAGGAAACTGTAGGGGAGGAAGACATTCGGCTGTTTACCAGACTCCGACACGAGTTCCACAAAT |
| *NFKB1* | NM_003998.2:1675 | NM_003998.2 | AGGGTATAGCTTCCCACACTATGGATTTCCTACTTATGGTGGGATTACTTTCCATCCTGGAACTACTAAATCTAATGCTGGGATGAAGCATGGAACCATG |
| *NFKB2* | NM_002502.2:825 | NM_002502.2 | ATCTCCGGGGGCATCAAACCTGAAGATTTCTCGAATGGACAAGACAGCAGGCTCTGTGCGGGGTGGAGATGAAGTTTATCTGCTTTGTGACAAGGTGCAG |
| *NFKBIA* | NM_020529.1:945 | NM_020529.1 | GGATGAGGAGAGCTATGACACAGAGTCAGAGTTCACGGAGTTCACAGAGGACGAGCTGCCCTATGATGACTGTGTGTTTGGAGGCCAGCGTCTGACGTTA |
| *NFKBIZ* | NM_001005474.1:2030 | NM_001005474.1 | ATTTGGTTCCCGATGGCCCTGTGGGAGAACAGATCCGACGTATCCTGAAGGGAAAGTCCATTCAGCAGAGAGCTCCACCGTATTAGCTCCATTAGCTTGG |
| *OAS1* | NM_001032409.1:805 | NM_001032409.1 | CTCCTGACGGTCTATGCTTGGGAGCGAGGGAGCATGAAAACACATTTCAACACAGCCCAGGGATTTCGGACGGTCTTGGAATTAGTCATAAACTACCAGC |
| *OAS2* | NM_016817.2:480 | NM_016817.2 | TGAAAAACAATTTCGAGATCCAGAAGTCCCTTGATGGGTTCACCATCCAGGTGTTCACAAAAAATCAGAGAATCTCTTTCGAGGTGCTGGCCGCCTTCAA |
| *POLR2A* | NM_000937.2:3775 | NM_000937.2 | TTCCAAGAAGCCAAAGACTCCTTCGCTTACTGTCTTCCTGTTGGGCCAGTCCGCTCGAGATGCTGAGAGAGCCAAGGATATTCTGTGCCGTCTGGAGCAT |
| *POU2F2* | NM_002698.2:908 | NM_002698.2 | GACGCAAGAAGAGGACCAGCATCGAGACAAACGTCCGCTTCGCCTTAGAGAAGAGTTTTCTAGCGAACCAGAAGCCTACCTCAGAGGAGATCCTGCTGAT |
| *PPIB* | NM_000942.4:272 | NM_000942.4 | AGAAGAAGAAGGGGCCCAAAGTCACCGTCAAGGTGTATTTTGACCTACGAATTGGAGATGAAGATGTAGGCCGGGTGATCTTTGGTCTCTTCGGAAAGAC |
| *PTGS2* | NM_000963.1:495 | NM_000963.1 | GCTACAAAAGCTGGGAAGCCTTCTCTAACCTCTCCTATTATACTAGAGCCCTTCCTCCTGTGCCTGATGATTGCCCGACTCCCTTGGGTGTCAAAGGTAA |
| *PTX3* | NM_002852.3:1152 | NM_002852.3 | ATATCTGGGATAGTGTTCTTAGCAATGAAGAGATAAGAGAGACCGGAGGAGCAGAGTCTTGTCACATCCGGGGGAATATTGTTGGGTGGGGAGTCACAGA |
| *RARRES3* | NM_004585.3:640 | NM_004585.3 | CTGACCCTCGTGCCCTGTCTCAGGCGTTCTCTAGATCCTTTCCTCTGTTTCCCTCTCTCGCTGGCAAAAGTATGATCTAATTGAAACAAGACTGAAGGAT |
| *RELB* | NM_006509.2:250 | NM_006509.2 | CACTCTCGCTCGCCGTTTCCAGGAGCACAGATGAATTGGAGATCATCGACGAGTACATCAAGGAGAACGGCTTCGGCCTGGACGGGGGACAGCCGGGCCC |
| *RIG1/DDX58* | NM_014314.3:2130 | NM_014314.3 | CTGGCATATTGACTGGACGTGGCAAAACAAATCAGAACACAGGAATGACCCTCCCGGCACAGAAGTGTATATTGGATGCATTCAAAGCCAGTGGAGATCA |
| *RPL19* | NM_000981.3:315 | NM_000981.3 | CCAATGCCCGAATGCCAGAGAAGGTCACATGGATGAGGAGAATGAGGATTTTGCGCCGGCTGCTCAGAAGATACCGTGAATCTAAGAAGATCGATCGCCA |
| *RPLP0* | NM_001002.3:250 | NM_001002.3 | CGAAATGTTTCATTGTGGGAGCAGACAATGTGGGCTCCAAGCAGATGCAGCAGATCCGCATGTCCCTTCGCGGGAAGGCTGTGGTGCTGATGGGCAAGAA |
| *S100A9* | NM_002965.2:75 | NM_002965.2 | AACATAGAGACCATCATCAACACCTTCCACCAATACTCTGTGAAGCTGGGGCACCCAGACACCCTGAACCAGGGGGAATTCAAAGAGCTGGTGCGAAAAG |
| *SLAM7* | NM_021181.3:215 | NM_021181.3 | GGGCACTATCATAGTGACCCAAAATCGTAATAGGGAGAGAGTAGACTTCCCAGATGGAGGCTACTCCCTGAAGCTCAGCAAACTGAAGAAGAATGACTCA |
| *SOCS1* | NM_003745.1:1025 | NM_003745.1 | TTAACTGTATCTGGAGCCAGGACCTGAACTCGCACCTCCTACCTCTTCATGTTTACATATACCCAGTATCTTTGCACAAACCAGGGGTTGGGGGAGGGTC |
| *SOCS3* | NM_003955.3:1870 | NM_003955.3 | GGAGGATGGAGGAGACGGGACATCTTTCACCTCAGGCTCCTGGTAGAGAAGACAGGGGATTCTACTCTGTGCCTCCTGACTATGTCTGGCTAAGAGATTC |
| *SRC* | NM_005417.3:1410 | NM_005417.3 | GGCATGAGAAGCTGGTGCAGTTGTATGCTGTGGTTTCAGAGGAGCCCATTTACATCGTCACGGAGTACATGAGCAAGGGGAGTTTGCTGGACTTTCTCAA |
| *STAT2* | NM_005419.2:1965 | NM_005419.2 | CCGTACACGAAGGAGGTGCTGCAGTCACTCCCGCTGACTGAAATCATCCGCCATTACCAGTTGCTCACTGAGGAGAATATACCTGAAAACCCACTGCGCT |
| *STING/TMEM173* | NM_198282.1:725 | NM_198282.1 | CTGGCATGGTCATATTACATCGGATATCTGCGGCTGATCCTGCCAGAGCTCCAGGCCCGGATTCGAACTTACAATCAGCATTACAACAACCTGCTACGGG |
| *Systeme 25 (Human endogenous retrovirus)* | Systeme_25.1:74 | Systeme_25.1 | AACGCGGGGGGCTCTGGGTGCTGTTAACCGGGCGAATTCCTGGGAACTGCGGGTATGGCTTGCCACAGTACCTTATCAGTTAATTGCATTCTTGGATGTG |
| *Systeme 47 (Human endogenous retrovirus)* | Systeme_47.1:0 | Systeme_47.1 | TAGACCAATGCAGTTAGGTGGCTCTTTCCAAGACTCTGGGGAAAAAAGTAGTAAAAAGCTAAATGCAATCAATCAGCAATTGAAAGCTAAGTGAGAGAGC |
| *TBET/TBX21* | NM_013351.1:890 | NM_013351.1 | ACACAGGAGCGCACTGGATGCGCCAGGAAGTTTCATTTGGGAAACTAAAGCTCACAAACAACAAGGGGGCGTCCAACAATGTGACCCAGATGATTGTGCT |
| *TBP* | NM_001172085.1:587 | NM_001172085.1 | ACAGTGAATCTTGGTTGTAAACTTGACCTAAAGACCATTGCACTTCGTGCCCGAAACGCCGAATATAATCCCAAGCGGTTTGCTGCGGTAATCATGAGGA |
| *TDRD9* | NM_153046.2:4211 | NM_153046.2 | CAGGAAGCTGTGGAGGCTGGATTCCAGGCTCCCTCCGCAGACTGACTTTCCTCTGTGTCTGGGTGTTACAGTCTGTGCCCACTGCATCCTAAAGGCCTTT |
| *TGFB1* | NM_000660.3:1260 | NM_000660.3 | TATATGTTCTTCAACACATCAGAGCTCCGAGAAGCGGTACCTGAACCCGTGTTGCTCTCCCGGGCAGAGCTGCGTCTGCTGAGGCTCAAGTTAAAAGTGG |
| *TIM3/HAVCR2* | NM_032782.3:955 | NM_032782.3 | TATATGAAGTGGAGGAGCCCAATGAGTATTATTGCTATGTCAGCAGCAGGCAGCAACCCTCACAACCTTTGGGTTGTCGCTTTGCAATGCCATAGATCCA |
| *TNFA* | NM_000594.2:1010 | NM_000594.2 | AGCAACAAGACCACCACTTCGAAACCTGGGATTCAGGAATGTGTGGCCTGCACAGTGAAGTGCTGGCAACCACTAAGAATTCAAACTGGGGCCTCCAGAA |
| *TNFAIP3* | NM_006290.2:260 | NM_006290.2 | CAAAGCCCTCATCGACAGAAACATCCAGGCCACCCTGGAAAGCCAGAAGAAACTCAACTGGTGTCGAGAAGTCCGGAAGCTTGTGGCGCTGAAAACGAAC |
| *TNFSF10* | NM_003810.2:115 | NM_003810.2 | GGGGGGACCCAGCCTGGGACAGACCTGCGTGCTGATCGTGATCTTCACAGTGCTCCTGCAGTCTCTCTGTGTGGCTGTAACTTACGTGTACTTTACCAAC |
| *TNFSF13B* | NM_006573.4:1430 | NM_006573.4 | CTTGGAGGAAGGACACAATTCAAAGGGGCAGTAAGGATTTTGTAAAACGTGGCATCCATAATTTACTATGGAGCAAGTGCCCACATCTCTAGGACATTAA |
| *ZAP70* | NM_001079.3:1175 | NM_001079.3 | GGAGCTCAAGGACAAGAAGCTCTTCCTGAAGCGCGATAACCTCCTCATAGCTGACATTGAACTTGGCTGCGGCAACTTTGGCTCAGTGCGCCAGGGCGTG |
| *ZBP1* | NM_001160419.2:680 | NM_001160419.2 | ATGAGGACAGCAAAAGATGTGAACCGAGACTTGTACAGGATGAAGAGCAGGCACCTTCTGGACATGGATGAGCAGTCCAAAGCATGGACGATTTACCGCC |
| *ZBTB16* | NM_006006.4:1585 | NM_006006.4 | TCCTGGATAGTTTGCGGCTGAGAATGCACTTACTGGCTCATTCAGCGGGTGCCAAAGCCTTTGTCTGTGATCAGTGCGGTGCACAGTTTTCGAAGGAGGA |

The grey filling indicates housekeeping genes (*HPRT1*, *DECR1*, *TBP*).
