## Supplementary material for "Functional alterations of immune gene expression in ICU and non-ICU patients with Legionnaires’ disease, a prospective observational study": Table S2

**Table S2: Characteristics of ICU-LD (with or without SS) and LD-unrelated-SS patients**

|  | **ICU-LD non-SS**  **(n=3)** | **ICU-LD SS**  **(n=3)** | **LD-unrelated-SS**  **(n=14)** | **P-value** |
| --- | --- | --- | --- | --- |
| **Demographics** | | | | |
| Sex, male, n (%) | 2 | 3 | 9 | 0.40 |
| Age (y), median [IQR] | 56 [45-66] | 63 [36-64] | 70 [58–75] | 0.11 |
| Body mass index (kg/m²), median [IQR] | 31 [24-32] | 24 [24-30] | 29 [21-40] | >0.9 |
| **Comorbidities, n (%)** | | | | |
| 0 | 0 | 0 | 1 | 0.51 |
| ≥1 | 2 | 3 | 13 |  |
| **Admission clinical and laboratory data, median [IQR]** | | | | |
| SOFA score | 8 [5-8] | 9 [8-12] | 8 [7-8] | 0.20 |
| Mean blood pressure, mmHg | 91 [89-97] | 74 [64-100] | 59 [54–64] | **0.007** |
| Creatininaemia, mg/L | 88 [60-283] | 78 [73-486] | 157 [111-247] | 0.50 |
| White blood cell count G/L | 11.0 [6.0-11.50] | 16.0 [8.9-29.6] | 9.2 [7.1-12.4] | 0.50 |
| Lymphocytes, G/L | 1.20 [0.27-1.37] | 0.60 [0.20-1.60] | 0.8 [0.5-1.3] | >0.9 |
| **Intensive care, ICU clinical and laboratory criteria** | | | | |
| PaO2/FiO2, median [IQR] | 86 [66-100] | 90 [85-222] | 226 [198–356] | **0.015** |
| Mechanical ventilation (MV), n (%) | 2 | 3 | 10 | 0.8 |
| Mechanical ventilation duration, days, median [IQR] | 8.5 [8.0-9.0] | 16.0 [9.0-44.0] | 3.5 [2.0-7.8] | 0.13 |
| Vasopressor administration, n (%) | 2 | 3 | 14 | 0.30 |
| Vasopressor therapy duration (days), median [IQR] | 1.5 [1.0-2.0] | 10.0 [2.0-25.0] | 3.0 [2.0-6.3] | 0.15 |
| Haemofiltration, n (%) | 0 | 1 | 3 (21) | 0.81 |
| Lactate concentration, mmol/L, median [IQR] | 1.6 [1.4-1.7] | 2.3 [2.2-2.6] | 2.8 [2.5-3.5] | **0.022** |
| Day2-Day3 mHLA-DR (Ab/C) | 10 905 [5 421-19 823] | 8 929 [2 122- 15 736] | 9 133 [4 734–11 449] | 0.70 |
| ICU length of stay, days, median [IQR] | 15.0 [8.0-20.0] | 11.0 [9.0-21.0] | 6.0 [4.0-13.0] | 0.90 |
| **Course** | | | | |
| Hospital length of stay, days, median [IQR] | 18.0 [10.0-20.0] | 21.0 [20.0-32.0] | 17.0^a^ [7.0–31.0] | 0.40 |
| Day 28 mortality, n (%) | 0 | 0 | 1 | 0.68 |

Comorbidities included diabetes (n=4), immunosuppressive anti-inflammatory biotherapy (methotrexate, n=1), hemopathy (n=1), chronic pulmonary disease (n=2), cerebrovascular accident (n=3), cardiac failure (n=4), dementia (n=1), and stomach ulcer (n=3).

FiO2: fraction of inspired oxygen; GU: Genome Unit; ICU: Intensive Care Unit; IQR: Interquartile Range; LD: Legionnaires’ disease; mHLA-DR: monocyte human leukocyte antigen DR expressed as number of sites per cell (Ab/C), usual values are 13 500 – 45 000Ab/C and the threshold value for immunosuppression is <8 000Ab/C [33]; MV: mechanical ventilation; NA: not applicable; PaO2: partial pressure of arterial oxygen; SOFA: Sequential Organ Failure Assessment; SS: Septic Shock. Chi-squared tests were used for comparison of sex, body mass index, primary site of infection, comorbidities, invasive ventilation, vasopressor administration, hemofiltration, SS, and Day-28 mortality between groups; the Mann-Whitney test was used for comparison of median comparisons of age, SOFA score, mean blood pressure, creatininaemia, white blood cells, lymphocytes, and PaO2/FiO2, lactate, and hospital and ICU length of stay. ^a^ n=12
