## Supplementary material for "Functional alterations of immune gene expression in ICU and non-ICU patients with Legionnaires’ disease, a prospective observational study": Table S3

**Table S3: Details of gene expression of all LD, non-ICU-LD, ICU-LD, and LD-unrelated-SS patients after LPS stimulation.**

| Gene | All LD (n=14) | | Non-ICU-LD (n=8) | | ICU-LD (n=6) | | LD-unrelated-SS (n=14) | |
| --- | --- | --- | --- | --- | --- | --- | --- | --- |
|  | Log(10)  FDR | Log2  (FC) | Log(10)  FDR | Log2  (FC) | Log(10)  FDR | Log2  (FC) | Log(10)  FDR | Log2  (FC) |
| *ADGRE3* | **4.1** | **1.6** | **2.8** | **1.4** | **2.7** | **2.0** | **4.5** | **1.8** |
| *ARL14EP* | 0.1 | 0.0 | 0.0 | 0.1 | 0.1 | 0.0 | 0.6 | 0.2 |
| *BST2* | 2.3 | -0.9 | 1.6 | -0.9 | 1.6 | -0.7 | **3.1** | **-1.1** |
| *C3* | **2.5** | **-2.1** | **1.4** | **-1.8** | **2.3** | **-2.7** | **2.6** | **-2.2** |
| *CCL2* | **1.9** | **-1.7** | 1.0 | -1.0 | **1.9** | **-2.6** | **3.3** | **-1.9** |
| *CCL20* | 0.5 | -0.6 | 0.1 | -0.3 | 0.9 | -2.7 | **1.6** | **-2.1** |
| *CCL4* | **4.0** | **-2.3** | **2.6** | **-2.0** | **2.7** | **-2.6** | **3.0** | **-2.9** |
| *CCL8* | **4.1** | **-3.7** | **2.8** | **-3.5** | **2.7** | **-4.6** | **4.5** | **-4.0** |
| *CCNB1IP1* | 0.0 | 0.1 | 0.1 | 0.2 | 0.1 | 0.1 | 1.3 | 0.5 |
| *CCR1* | 0.6 | -0.4 | 0.0 | 0.2 | 1.9 | -0.9 | 3.0 | -0.9 |
| *CD127* | 3.3 | 0.5 | 2.8 | 0.5 | 1.6 | 0.5 | 2.0 | 0.4 |
| *CD209* | 0.5 | -0.4 | 1.5 | -0.9 | 0.2 | 0.1 | 0.1 | -0.4 |
| *CD3D* | 2.2 | 0.3 | 1.5 | 0.2 | 1.5 | 0.3 | 1.6 | 0.4 |
| *CD44* | 0.1 | -0.3 | 0.0 | 0.1 | 0.2 | -0.3 | 1.1 | -0.5 |
| *CD74* | 0.1 | 0.1 | 0.1 | -0.1 | 0.4 | 0.1 | 0.1 | 0.0 |
| *CD83* | **1.9** | **-1.2** | 1.0 | -1.2 | **1.9** | **-1.1** | **4.0** | **-1.4** |
| *CDKN1A* | **2.2** | **-1.1** | 1.1 | -0.8 | **2.3** | **-1.5** | **2.3** | **-1.3** |
| *CLEC7A* | 1.0 | 0.8 | 0.2 | 0.4 | 1.7 | 1.0 | 1.1 | 0.5 |
| *CX3CR1* | **2.7** | **1.7** | 1.4 | 0.9 | **2.7** | **1.9** | **3.7** | **1.9** |
| *CXCL10* | **3.2** | **-4.8** | **2.8** | **-4.6** | **1.6** | **-5.1** | **4.5** | **-4.7** |
| *CXCL2* | 1.9 | -0.9 | 0.8 | -0.9 | **2.3** | **-1.5** | **3.2** | **-1.6** |
| *CXCL9* | **2.5** | **-3.5** | **2.6** | **-3.8** | 1.0 | -3.0 | **3.2** | **-3.2** |
| *DYRK2* | 1.1 | 0.1 | 0.8 | 0.1 | 0.8 | 0.3 | 1.0 | 0.2 |
| *EIF2AK4* | 0.9 | 0.3 | 0.5 | 0.3 | 0.8 | 0.5 | 1.6 | 0.5 |
| *FAM89A* | 0.7 | -0.4 | 0.6 | -0.7 | 0.4 | -0.2 | 0.2 | 0.1 |
| *FOXP3* | 0.5 | -0.3 | 0.6 | -0.6 | 0.2 | -0.1 | 1.1 | -0.8 |
| *GAPDH* | 0.8 | -0.4 | 0.1 | -0.1 | 1.6 | -0.9 | 1.8 | -0.9 |
| *GATA3* | 0.3 | 0.3 | 0.0 | 0.1 | 0.6 | 0.5 | 1.5 | 0.6 |
| *HLA-DMB* | 1.3 | -0.4 | 2.0 | -0.8 | 0.1 | -0.2 | 0.2 | 0.1 |
| *HLA-DPA1* | 0.1 | -0.1 | 0.3 | -0.3 | 0.1 | 0.1 | 0.2 | 0.2 |
| *HLA-DPB1* | 1.0 | 0.3 | 0.4 | 0.1 | 1.3 | 0.5 | 1.7 | 0.5 |
| *HLA-DRA* | 0.0 | -0.2 | 0.3 | -0.7 | 0.3 | 0.5 | 2.2 | -0.7 |
| *IDO1* | **1.9** | **-2.0** | 1.3 | -1.3 | **1.3** | **-3.8** | **3.0** | **-3.5** |
| *IFI27* | **1.9** | **-1.1** | 0.8 | -1.0 | **2.3** | **-1.1** | **3.9** | **-1.2** |
| *IFI35* | **2.7** | **-1.6** | **2.0** | **-1.5** | **1.7** | **-1.6** | **4.0** | **-2.1** |
| *IFI44L* | **2.3** | **-2.0** | 1.0 | -1.2 | **2.7** | **-2.5** | **3.2** | **-2.9** |
| *IFIH1* | **3.1** | **-1.5** | **1.9** | **-1.1** | **2.5** | **-1.6** | **3.9** | **-1.8** |
| *IFITM1* | 0.5 | -0.3 | 0.0 | -0.2 | 1.2 | -0.8 | 0.6 | -0.4 |
| *IFNb1* | **2.0** | **-1.0** | **1.9** | **-1.5** | 0.9 | -0.6 | 0.3 | -0.2 |
| *IFNg* | **2.6** | **-4.0** | **2.3** | **-4.2** | **1.3** | **-3.9** | **4.5** | **-3.6** |
| *IL10* | 0.2 | 0.7 | 0.6 | 1.2 | 0.2 | -0.3 | 1.7 | -0.9 |
| *IL12B* | **3.6** | **-5.3** | **2.5** | **-5.2** | **2.5** | **-5.3** | **3.9** | **-4.9** |
| *IL18* | 1.5 | -0.4 | 0.8 | -0.3 | 1.6 | -0.5 | 1.9 | -0.2 |
| *IL18R1* | 1.4 | 0.9 | 0.8 | 0.5 | **1.3** | **1.4** | **2.0** | **0.7** |
| *IL1A* | 0.4 | -0.4 | 0.1 | -0.1 | 0.6 | -1.9 | **2.3** | **-2.7** |
| *IL1B* | 0.9 | -0.9 | 0.4 | -0.2 | 1.0 | -1.4 | **1.5** | **-1.5** |
| *IL1R2* | 1.9 | 0.6 | 1.3 | 1.0 | 1.3 | 0.5 | 1.1 | 0.7 |
| *IL2* | 1.0 | -1.2 | **1.6** | **-1.7** | 0.1 | -0.6 | 0.1 | 0.0 |
| *IL6* | **1.9** | **2.6** | **2.5** | **3.0** | 0.4 | 2.4 | **1.5** | **-2.1** |
| *IRAK2* | **1.3** | **-1.3** | 0.5 | -0.9 | **1.7** | **-1.8** | **2.3** | **-1.6** |
| *IRF3* | 0.3 | 0.3 | 0.3 | 0.3 | 0.1 | 0.2 | 0.5 | 0.2 |
| *IRF7* | **2.4** | **-1.2** | 1.3 | -0.8 | **2.5** | **-1.7** | **3.2** | **-1.9** |
| *JAK2* | 2.3 | -0.4 | 2.1 | -0.7 | 1.0 | -0.6 | 4.0 | -0.8 |
| *LILRB1* | **3.8** | **-1.2** | **2.6** | **-1.2** | **2.5** | **-1.2** | **1.8** | **-1.0** |
| *MDC1* | 0.2 | 0.0 | 0.1 | 0.1 | 0.2 | 0.0 | 0.2 | 0.1 |
| *MERTK* | 0.4 | 0.0 | 0.2 | 0.1 | 0.3 | -0.1 | 0.2 | 0.3 |
| *MX1* | **2.3** | **-1.2** | 1.1 | -0.9 | **2.7** | **-1.8** | **2.2** | **-2.1** |
| *NFKB1* | 4.2 | -1.0 | 2.8 | -0.9 | **2.7** | **-1.1** | **3.9** | **-1.2** |
| *NFKB2* | **4.2** | **-1.7** | **2.8** | **-1.2** | **2.7** | **-2.3** | **4.0** | **-2.3** |
| *NFKBIA* | **4.0** | **-1.7** | 2.6 | -0.8 | **2.7** | **-2.4** | **3.9** | **-2.2** |
| *NFKBIZ* | **4.2** | **-1.9** | **2.8** | **-1.1** | **2.7** | **-2.1** | **3.6** | **-2.1** |
| *OAS1* | **3.3** | **-2.3** | **2.0** | **-1.5** | **2.7** | **-2.8** | **4.0** | **-3.2** |
| *OAS2* | **2.4** | **-1.1** | 1.7 | -0.5 | **1.6** | **-1.2** | **3.2** | **-1.7** |
| *POLR2A* | 0.4 | 0.1 | 0.4 | 0.1 | 0.1 | 0.0 | 0.6 | -0.1 |
| *POU2F2* | 0.4 | -0.4 | 0.5 | -0.7 | 0.1 | -0.2 | 1.0 | -0.5 |
| *PPIB* | 0.1 | 0.0 | 0.0 | 0.0 | 0.2 | 0.0 | 0.0 | 0.1 |
| *PTGS2* | 0.1 | 0.3 | 0.3 | 0.5 | 0.8 | -1.8 | **1.4** | **-1.8** |
| *PTX3* | 1.3 | -0.7 | 1.1 | -0.8 | 0.7 | -0.6 | 0.6 | -0.4 |
| *RARRES3* | 1.5 | -0.3 | 1.3 | -0.2 | 0.8 | -0.6 | 2.8 | -0.4 |
| *RELB* | 2.6 | -0.7 | 1.3 | -0.5 | **2.7** | **-1.1** | 4.5 | -0.9 |
| *RIG1* | 2.4 | -0.9 | 1.3 | -0.7 | **2.3** | **-1.2** | **3.4** | **-1.4** |
| *RPL19* | 3.5 | 0.5 | 2.1 | 0.4 | 2.7 | 0.6 | 3.1 | 0.6 |
| *RPLP0* | 1.9 | 0.4 | 0.8 | 0.2 | 2.3 | 0.4 | 1.6 | 0.4 |
| *S100A9* | 0.9 | 0.4 | 1.3 | 0.6 | 0.1 | 0.1 | 0.2 | 0.4 |
| *SLAM7* | **4.2** | **-2.6** | **2.8** | **-2.6** | **2.7** | **-3.4** | **4.2** | **-2.9** |
| *SOCS1* | **2.0** | **-1.4** | **1.6** | **-1.3** | 1.1 | -1.7 | **4.5** | **-2.8** |
| *SOCS3* | 1.1 | -0.5 | 0.6 | -0.3 | 1.1 | -1.6 | **4.0** | **-1.7** |
| *SRC* | 1.8 | -1.0 | 0.6 | -0.5 | **2.7** | **-1.5** | **3.0** | **-1.5** |
| *STAT2* | **3.0** | **-1.4** | **1.6** | **-1.4** | **2.7** | **-1.5** | **4.0** | **-2.0** |
| *STING* | 0.2 | 0.1 | 0.1 | 0.1 | 0.2 | 0.1 | 0.1 | 0.1 |
| *Systeme 25* | 0.8 | -0.3 | 1.3 | -0.7 | 0.1 | -0.1 | 0.1 | 0.1 |
| *Systeme 47* | **3.1** | **-1.5** | 2.0 | -0.9 | **2.3** | **-2.5** | **3.2** | **-2.4** |
| *TBET* | 2.0 | 0.5 | 1.3 | 0.4 | 1.6 | 0.5 | 1.9 | 0.7 |
| *TDRD9* | 1.4 | 0.4 | 1.3 | 0.7 | 0.7 | 0.3 | 0.1 | 0.1 |
| *TGFB1* | 0.4 | 0.2 | 0.3 | 0.2 | 0.3 | 0.3 | 0.0 | -0.1 |
| *TIM3* | **3.8** | **-1.4** | **2.5** | **-1.2** | **2.7** | **-2.0** | **4.5** | **-1.8** |
| *TNFA* | **3.6** | **-1.8** | **2.8** | **-1.7** | **2.1** | **-2.0** | **3.7** | **-2.5** |
| *TNFAIP3* | 1.5 | -0.5 | 0.8 | -0.4 | 1.5 | -0.9 | 1.1 | -0.4 |
| *TNFSF10* | **3.2** | **-1.6** | **1.9** | **-1.4** | **2.7** | **-1.9** | **3.9** | **-2.2** |
| *TNFSF13B* | **2.3** | **-1.5** | **1.5** | **-1.3** | **1.7** | **-1.6** | **3.9** | **-1.9** |
| *ZAP70* | 0.9 | 0.1 | 0.6 | 0.1 | 0.7 | 0.3 | 0.3 | 0.2 |
| *ZBP1* | 0.3 | 0.0 | 0.1 | 0.0 | 0.3 | -0.1 | 1.3 | -0.7 |
| *ZBTB16* | 1.3 | 0.4 | 1.3 | 0.5 | 0.5 | 0.4 | 1.6 | 0.5 |

Genes were classified by alphabetic number. Values in bold are DEGs. FDR: false discovery rate, FC: fold change
