## Supplementary material for "Functional alterations of immune gene expression in ICU and non-ICU patients with Legionnaires’ disease, a prospective observational study": Table S4

| #category | term description | observed gene count | signal | FDR | matching proteins in your network (labels) |
|---|---|---|---|---|---|
| GO Process | Cytokine-mediated signaling pathway | 19 | 3.76 | 6.80E-21 | NFKBIA,CCL2,IFNG,IL12B,IRAK2,CXCL10,STAT2,LILRB1,OAS2,SRC,TNFSF13B,CCL8,IRF7,MX1,OAS1,TNF,CXCL2,CCL4,IFI27 |
| GO Process | Response to virus | 18 | 3.6 | 1.74E-19 | NFKB1,IFNG,IL12B,CXCL10,STAT2,LILRB1,OAS2,IFI44L,SRC,DDX58,CCL8,IRF7,MX1,OAS1,TNF,CCL4,IFI27,IFIH1 |
| GO Process | Defense response to virus | 13 | 2.95 | 1.95E-13 | IFNG,IL12B,CXCL10,STAT2,LILRB1,OAS2,IFI44L,DDX58,IRF7,MX1,OAS1,IFI27,IFIH1 |
| GO Process | Cellular response to lipopolysaccharide | 11 | 2.76 | 1.59E-11 | NFKBIA,CCL2,NFKB1,IL12B,IRAK2,CXCL10,HAVCR2,LILRB1,SRC,TNF,CXCL2 |
| GO Process | Regulation of interferon-beta production | 7 | 2.76 | 4.94E-09 | RELB,LILRB1,OAS2,DDX58,IRF7,OAS1,IFIH1 |
| GO Process | Pattern recognition receptor signaling pathway | 8 | 2.57 | 3.67E-09 | NFKBIA,IRAK2,HAVCR2,DDX58,IRF7,OAS1,TNF,IFIH1 |
| GO Process | Regulation of type I interferon production | 8 | 2.54 | 4.16E-09 | RELB,HAVCR2,LILRB1,OAS2,DDX58,IRF7,OAS1,IFIH1 |
| GO Process | Innate immune response | 21 | 2.48 | 1.72E-18 | RELB,CCL2,IFNG,IL12B,C3,CXCL10,HAVCR2,STAT2,OAS2,SLAMF7,SRC,DDX58,CCL8,IRF7,MX1,OAS1,IFI35,TNF,CCL4,IFI27,IFIH1 |
| GO Process | Cellular response to cytokine stimulus | 20 | 2.45 | 1.64E-17 | NFKBIA,CCL2,NFKB1,IFNG,IL12B,IRAK2,CXCL10,STAT2,LILRB1,OAS2,SRC,TNFSF13B,CCL8,IRF7,MX1,OAS1,TNF,CXCL2,CCL4,IFI27 |
| GO Process | Response to cytokine | 21 | 2.36 | 5.81E-18 | NFKBIA,RELB,CCL2,NFKB1,IFNG,IL12B,IRAK2,CXCL10,STAT2,LILRB1,OAS2,SRC,TNFSF13B,CCL8,IRF7,MX1,OAS1,TNF,CXCL2,CCL4, |
| GO Process | Defense response to other organism | 24 | 2.35 | 1.21E-20 | RELB,CCL2,IFNG,IL12B,C3,CXCL10,HAVCR2,STAT2,LILRB1,OAS2,SLAMF7,IFI44L,SRC,DDX58,CCL8,IRF7,MX1,OAS1,IFI35,TNF,CXCL2 |
| GO Process | Response to type I interferon | 6 | 2.34 | 1.29E-07 | STAT2,OAS2,IRF7,MX1,OAS1,IFI27 |
| GO Process | Response to lipopolysaccharide | 12 | 2.29 | 5.83E-11 | NFKBIA,CCL2,NFKB1,IL12B,IRAK2,CXCL10,HAVCR2,LILRB1,SRC,TNF,CXCL2,IDO1 |
| GO Process | Cellular response to virus | 7 | 2.28 | 5.76E-08 | NFKB1,IFNG,CXCL10,DDX58,IRF7,OAS1,IFIH1 |
| GO Process | Immune response | 28 | 2.21 | 1.29E-23 | RELB,CCL2,IFNG,IL12B,TNFSF10,C3,CXCL10,HAVCR2,STAT2,LILRB1,OAS2,SLAMF7,NFKB2,IFI44L,SRC,TNFSF13B,CD83,DDX58,CCL8 |
| GO Process | Defense response | 29 | 2.2 | 1.79E-24 | RELB,CCL2,NFKB1,IFNG,IL12B,C3,IRAK2,CXCL10,HAVCR2,STAT2,LILRB1,NFKBIZ,OAS2,SLAMF7,IFI44L,SRC,CD83,DDX58,CCL8,IRF7 |
| GO Process | Response to other organism | 28 | 2.2 | 1.29E-23 | NFKBIA,RELB,CCL2,NFKB1,IFNG,IL12B,C3,IRAK2,CXCL10,HAVCR2,STAT2,LILRB1,OAS2,SLAMF7,IFI44L,SRC,DDX58,CCL8,IRF7,MX1 |
| GO Process | Positive regulation of tumor necrosis factor production | 7 | 2.13 | 1.24E-07 | IFNG,IL12B,HAVCR2,OAS2,DDX58,OAS1,IFIH1 |
| GO Process | Positive regulation of defense response | 11 | 2.12 | 8.20E-10 | NFKBIA,IFNG,IL12B,C3,HAVCR2,NFKBIZ,SRC,IRF7,IFI35,TNF,IDO1 |
| GO Process | Positive regulation of inflammatory response | 8 | 2.11 | 4.57E-08 | NFKBIA,IFNG,IL12B,C3,NFKBIZ,IFI35,TNF,IDO1 |
| GO Process | Positive regulation of cytokine production | 14 | 2.07 | 1.59E-11 | IFNG,IL12B,C3,HAVCR2,LILRB1,OAS2,SRC,CD83,DDX58,IRF7,OAS1,TNF,IDO1,IFIH1 |
| GO Process | Immune response-regulating signaling pathway | 11 | 2.04 | 1.36E-09 | NFKBIA,IRAK2,HAVCR2,LILRB1,NFKBIZ,SRC,DDX58,IRF7,OAS1,TNF,IFIH1 |
| GO Process | Regulation of defense response | 16 | 1.99 | 1.46E-12 | NFKBIA,NFKB1,IFNG,IL12B,C3,HAVCR2,STAT2,LILRB1,NFKBIZ,SRC,DDX58,IRF7,OAS1,IFI35,TNF,IDO1 |
| GO Process | Positive regulation of response to external stimulus | 13 | 1.97 | 1.39E-10 | NFKBIA,IFNG,IL12B,C3,CXCL10,HAVCR2,NFKBIZ,SRC,IRF7,IFI35,TNF,IDO1,CCL4 |
| GO Process | Regulation of tumor necrosis factor production | 8 | 1.97 | 9.77E-08 | IFNG,IL12B,HAVCR2,LILRB1,OAS2,DDX58,OAS1,IFIH1 |
| GO Process | Positive regulation of interferon-beta production | 5 | 1.97 | 2.51E-06 | OAS2,DDX58,IRF7,OAS1,IFIH1 |
| GO Process | Positive regulation of leukocyte cell-cell adhesion | 10 | 1.95 | 9.49E-09 | CCL2,IFNG,IL12B,HAVCR2,LILRB1,NFKBIZ,SRC,TNFSF13B,CD83,TNF |
| GO Process | Type I interferon signaling pathway | 5 | 1.95 | 2.77E-06 | STAT2,OAS2,IRF7,OAS1,IFI27 |
| GO Process | Positive regulation of immune system process | 19 | 1.93 | 2.02E-14 | CCL2,IFNG,IL12B,C3,CXCL10,HAVCR2,LILRB1,NFKBIZ,SRC,TNFSF13B,CD83,DDX58,CCL8,IRF7,CDKN1A,IFI35,TNF,IDO1,CCL4 |
| GO Process | Inflammatory response | 14 | 1.91 | 5.69E-11 | RELB,CCL2,NFKB1,IFNG,C3,IRAK2,CXCL10,HAVCR2,NFKBIZ,CCL8,TNF,CXCL2,IDO1,CCL4 |
| GO Process | Negative regulation of cytokine production | 10 | 1.9 | 1.28E-08 | RELB,NFKB1,IFNG,IL12B,HAVCR2,LILRB1,CD83,OAS1,TNF,IDO1 |
| GO Process | Regulation of immune response | 18 | 1.85 | 2.29E-13 | NFKBIA,IFNG,IL12B,C3,IRAK2,HAVCR2,STAT2,LILRB1,NFKBIZ,SRC,TNFSF13B,DDX58,IRF7,OAS1,IFI35,TNF,IDO1,IFIH1 |
| GO Process | Regulation of innate immune response | 9 | 1.84 | 6.58E-08 | IL12B,C3,HAVCR2,STAT2,LILRB1,SRC,IRF7,OAS1,IFI35 |
| GO Process | Regulation of leukocyte cell-cell adhesion | 11 | 1.81 | 6.86E-09 | CCL2,IFNG,IL12B,HAVCR2,LILRB1,NFKBIZ,SRC,TNFSF13B,CD83,TNF,IDO1 |
| GO Process | Regulation of response to external stimulus | 19 | 1.78 | 1.13E-13 | NFKBIA,CCL2,NFKB1,IFNG,IL12B,C3,CXCL10,HAVCR2,STAT2,LILRB1,NFKBIZ,SRC,DDX58,IRF7,OAS1,IFI35,TNF,IDO1,CCL4 |
| GO Process | Positive regulation of T cell activation | 9 | 1.78 | 9.68E-08 | CCL2,IFNG,IL12B,HAVCR2,LILRB1,NFKBIZ,SRC,TNFSF13B,CD83 |
| GO Process | Regulation of adaptive immune response | 8 | 1.78 | 3.02E-07 | IL12B,C3,HAVCR2,LILRB1,NFKBIZ,TNFSF13B,IRF7,TNF |
| GO Process | Antiviral innate immune response | 4 | 1.78 | 1.52E-05 | CXCL10,DDX58,MX1,OAS1 |
| GO Process | Regulation of cytokine production | 16 | 1.77 | 1.26E-11 | RELB,NFKB1,IFNG,IL12B,C3,HAVCR2,LILRB1,OAS2,SRC,CD83,DDX58,IRF7,OAS1,TNF,IDO1,IFIH1 |
| GO Process | Response to bacterium | 15 | 1.77 | 4.40E-11 | NFKBIA,CCL2,NFKB1,IL12B,C3,IRAK2,CXCL10,HAVCR2,LILRB1,OAS2,SRC,OAS1,TNF,CXCL2,IDO1 |
| GO Process | Positive regulation of leukocyte activation | 11 | 1.76 | 1.01E-08 | CCL2,IFNG,IL12B,HAVCR2,LILRB1,NFKBIZ,SRC,TNFSF13B,CD83,CDKN1A,TNF |
| GO Process | Negative regulation of viral genome replication | 5 | 1.72 | 9.18E-06 | OAS2,MX1,OAS1,TNF,IFIH1 |
| GO Process | Regulation of immune system process | 24 | 1.7 | 3.90E-17 | NFKBIA,CCL2,IFNG,IL12B,C3,IRAK2,CXCL10,HAVCR2,STAT2,LILRB1,NFKBIZ,SRC,TNFSF13B,CD83,DDX58,CCL8,IRF7,CDKN1A,OAS1 |
| GO Process | Positive regulation of lymphocyte activation | 10 | 1.69 | 5.76E-08 | CCL2,IFNG,IL12B,HAVCR2,LILRB1,NFKBIZ,SRC,TNFSF13B,CD83,CDKN1A |
| GO Process | Regulation of calcidiol 1-monooxygenase activity | 3 | 1.68 | 4.74E-05 | NFKB1,IFNG,TNF |
| GO Process | Cellular response to interferon-gamma | 6 | 1.66 | 5.13E-06 | CCL2,IFNG,IL12B,CCL8,TNF,CCL4 |
| GO Process | Toll-like receptor signaling pathway | 5 | 1.65 | 1.34E-05 | NFKBIA,IRAK2,HAVCR2,OAS1,TNF |
| GO Process | Cytoplasmic pattern recognition receptor signaling pathw | 4 | 1.63 | 3.31E-05 | NFKBIA,DDX58,IRF7,IFIH1 |
| GO Process | Positive regulation of immune response | 12 | 1.62 | 8.60E-09 | IFNG,IL12B,C3,HAVCR2,LILRB1,NFKBIZ,SRC,TNFSF13B,IRF7,IFI35,TNF,IDO1 |

| | | | | | |
|---|---|---|---|---|---|
| GO Process | Regulation of interleukin-12 production | 5 | 1.62 | 1.60E-05 | NFKB1,IFNG,IL12B,LILRB1,IDO1 |
| GO Process | Regulation of vitamin D biosynthetic process | 3 | 1.62 | 6.37E-05 | NFKB1,IFNG,TNF |
| GO Process | interleukin-27-mediated signaling pathway | 3 | 1.62 | 6.37E-05 | OAS2,MX1,OAS1 |
| GO Process | Adaptive immune response | 10 | 1.61 | 9.68E-08 | RELB,IFNG,IL12B,C3,HAVCR2,LILRB1,SLAMF7,NFKB2,TNFSF13B,IRF7 |
| GO Process | Regulation of response to biotic stimulus | 10 | 1.61 | 9.77E-08 | IL12B,C3,HAVCR2,STAT2,LILRB1,SRC,DDX58,IRF7,OAS1,IFI35 |
| GO Process | Immune system process | 30 | 1.58 | 7.42E-22 | RELB,CCL2,IFNG,IL12B,TNFSF10,C3,CXCL10,HAVCR2,STAT2,LILRB1,NFKBIZ,OAS2,SLAMF7,NFKB2,IFI44L,SRC,TNFSF13B,CD83,DDX |
| GO Process | Regulation of inflammatory response | 10 | 1.58 | 1.21E-07 | NFKBIA,NFKB1,IFNG,IL12B,C3,NFKBIZ,SRC,IFI35,TNF,IDO1 |
| GO Process | Immune effector process | 10 | 1.57 | 1.29E-07 | RELB,IFNG,IL12B,C3,HAVCR2,LILRB1,SLAMF7,SRC,IRF7,IFI35 |
| GO Process | Regulation of T cell activation | 10 | 1.57 | 1.29E-07 | CCL2,IFNG,IL12B,HAVCR2,LILRB1,NFKBIZ,SRC,TNFSF13B,CD83,IDO1 |
| GO Process | Regulation of interferon-alpha production | 4 | 1.57 | 4.61E-05 | HAVCR2,DDX58,IRF7,IFIH1 |
| GO Process | Cell activation | 14 | 1.56 | 1.29E-09 | RELB,IFNG,IL12B,CXCL10,HAVCR2,LILRB1,SLAMF7,NFKB2,SRC,TNFSF13B,CD83,IFI35,TNF,IDO1 |
| GO Process | Cellular response to tumor necrosis factor | 7 | 1.55 | 3.56E-06 | NFKBIA,CCL2,NFKB1,TNFSF13B,CCL8,TNF,CCL4 |
| GO Process | Positive regulation of CD4-positive, alpha-beta T cell dif | 4 | 1.54 | 5.49E-05 | IFNG,IL12B,NFKBIZ,CD83 |
| GO Process | Toll-like receptor 3 signaling pathway | 3 | 1.54 | 0.0001 | HAVCR2,OAS1,TNF |
| GO Process | Regulation of adaptive immune response based on somati | 7 | 1.53 | 4.10E-06 | IL12B,C3,HAVCR2,LILRB1,NFKBIZ,TNFSF13B,TNF |
| GO Process | Response to interleukin-1 | 6 | 1.5 | 1.29E-05 | CCL2,NFKB1,IRAK2,SRC,CCL8,CCL4 |
| GO Process | Neutrophil chemotaxis | 5 | 1.45 | 4.20E-05 | CCL2,CXCL10,CCL8,CXCL2,CCL4 |
| GO Process | Cytoplasmic pattern recognition receptor signaling pathw | 3 | 1.45 | 0.00016 | DDX58,IRF7,IFIH1 |
| GO Process | Lipopolysaccharide-mediated signaling pathway | 4 | 1.44 | 9.10E-05 | NFKBIA,CCL2,IRAK2,TNF |
| GO Process | Regulation of lymphocyte activation | 11 | 1.43 | 1.29E-07 | CCL2,IFNG,IL12B,HAVCR2,LILRB1,NFKBIZ,SRC,TNFSF13B,CD83,CDKN1A,IDO1 |
| GO Process | Humoral immune response | 8 | 1.43 | 2.93E-06 | CCL2,IFNG,C3,CXCL10,CD83,CCL8,TNF,CXCL2 |
| GO Process | Chemokine-mediated signaling pathway | 5 | 1.43 | 4.61E-05 | CCL2,CXCL10,CCL8,CXCL2,CCL4 |
| GO Process | Regulation of leukocyte activation | 12 | 1.41 | 5.76E-08 | CCL2,IFNG,IL12B,HAVCR2,LILRB1,NFKBIZ,SRC,TNFSF13B,CD83,CDKN1A,TNF,IDO1 |
| GO Process | Regulation of inflammatory response to antigenic stimulu | 4 | 1.41 | 0.00011 | IL12B,C3,SRC,TNF |
| GO Process | Response to external stimulus | 29 | 1.38 | 4.45E-19 | NFKBIA,RELB,CCL2,NFKB1,IFNG,IL12B,C3,IRAK2,CXCL10,HAVCR2,STAT2,LILRB1,OAS2,SLAMF7,IFI44L,SRC,DDX58,CCL8,IRF7,MX1 |
| GO Process | Negative regulation of response to external stimulus | 9 | 1.31 | 2.77E-06 | CCL2,NFKB1,IL12B,HAVCR2,STAT2,LILRB1,SRC,OAS1,TNF |
| GO Process | Cellular response to interleukin-1 | 5 | 1.31 | 9.10E-05 | CCL2,NFKB1,IRAK2,CCL8,CCL4 |
| GO Process | Regulation of lipid storage | 4 | 1.3 | 0.0002 | NFKBIA,NFKB1,C3,TNF |
| GO Process | Leukocyte activation | 11 | 1.29 | 4.12E-07 | RELB,IFNG,IL12B,HAVCR2,LILRB1,SLAMF7,TNFSF13B,CD83,IFI35,TNF,IDO1 |
| GO Process | Negative regulation of defense response | 7 | 1.29 | 1.94E-05 | NFKB1,IL12B,HAVCR2,STAT2,LILRB1,SRC,OAS1 |
| GO Process | Regulation of cell killing | 5 | 1.29 | 0.0001 | IFNG,IL12B,C3,HAVCR2,LILRB1 |
| GO Process | Regulation of NIK/NF-kappaB signaling | 5 | 1.29 | 0.0001 | NFKBIA,IL12B,HAVCR2,IFI35,TNF |
| GO Process | Lymphocyte chemotaxis | 4 | 1.29 | 0.00021 | CCL2,CXCL10,CCL8,CCL4 |
| GO Process | Regulation of response to cytokine stimulus | 6 | 1.28 | 4.79E-05 | IRAK2,STAT2,DDX58,IRF7,OAS1,IFIH1 |
| GO Process | Eosinophil chemotaxis | 3 | 1.27 | 0.00043 | CCL2,CCL8,CCL4 |
| GO Process | Adaptive immune response based on somatic recombinati | 6 | 1.26 | 5.63E-05 | RELB,IL12B,C3,NFKB2,TNFSF13B,IRF7 |
| GO Process | Leukocyte migration | 7 | 1.23 | 3.00E-05 | CCL2,CXCL10,SRC,CCL8,TNF,CXCL2,CCL4 |
| GO Process | Positive regulation of adaptive immune response based o | 5 | 1.23 | 0.00014 | IL12B,C3,NFKBIZ,TNFSF13B,TNF |
| GO Process | Macrophage activation involved in immune response | 3 | 1.23 | 0.00054 | IFNG,HAVCR2,IFI35 |
| GO Process | Positive regulation of programmed cell death | 10 | 1.22 | 2.28E-06 | CCL2,IFNG,IL12B,TNFSF10,C3,LILRB1,SRC,CDKN1A,TNF,IDO1 |
| GO Process | Response to dsRNA | 4 | 1.22 | 0.00031 | NFKBIA,NFKB1,DDX58,IFIH1 |
| GO Process | Negative regulation of immune system process | 9 | 1.2 | 6.42E-06 | CCL2,IL12B,HAVCR2,STAT2,LILRB1,SRC,OAS1,TNF,IDO1 |
| GO Process | Regulation of alpha-beta T cell activation | 5 | 1.2 | 0.00017 | IFNG,IL12B,LILRB1,NFKBIZ,CD83 |
| GO Process | Negative regulation of immune response | 6 | 1.19 | 8.70E-05 | IL12B,HAVCR2,STAT2,LILRB1,SRC,OAS1 |
| GO Process | Regulation of interleukin-10 production | 4 | 1.19 | 0.00037 | IL12B,LILRB1,CD83,IDO1 |
| GO Process | Cellular response to organic substance | 23 | 1.18 | 1.23E-12 | NFKBIA,CCL2,NFKB1,IFNG,IL12B,IRAK2,CXCL10,HAVCR2,STAT2,LILRB1,OAS2,SRC,TNFSF13B,DDX58,CCL8,IRF7,MX1,OAS1,TNF,CX |
| GO Process | Positive regulation of immune effector process | 7 | 1.18 | 4.20E-05 | IFNG,IL12B,C3,LILRB1,NFKBIZ,DDX58,TNF |
| GO Process | Negative regulation of interleukin-10 production | 3 | 1.18 | 0.00069 | IL12B,LILRB1,IDO1 |
| GO Process | Leukocyte activation involved in immune response | 6 | 1.17 | 9.86E-05 | RELB,IFNG,IL12B,HAVCR2,LILRB1,IFI35 |
| GO Process | Macrophage activation | 4 | 1.17 | 0.00041 | IFNG,HAVCR2,IFI35,TNF |
| GO Process | Regulation of leukocyte proliferation | 7 | 1.16 | 4.79E-05 | IL12B,HAVCR2,LILRB1,TNFSF13B,CCL8,CDKN1A,IDO1 |
| GO Process | Regulation of myeloid leukocyte differentiation | 5 | 1.15 | 0.00023 | IFNG,IL12B,LILRB1,IRF7,TNF |
| GO Process | Positive regulation of vitamin D biosynthetic process | 2 | 1.15 | 0.0012 | IFNG,TNF |
| GO Process | Positive regulation of NIK/NF-kappaB signaling | 4 | 1.13 | 0.00052 | IL12B,HAVCR2,IFI35,TNF |

| | | | | | |
|---|---|---|---|---|---|
| GO Process | Cellular response to dsRNA | 3 | 1.13 | 0.00093 | NFKB1,DDX58,IFIH1 |
| GO Process | Cell killing | 5 | 1.12 | 0.00028 | C3,CXCL10,SLAMF7,CCL8,CXCL2 |
| GO Process | I-kappaB kinase/NF-kappaB signaling | 4 | 1.12 | 0.00054 | NFKBIA,RELB,IRAK2,TNF |
| GO Process | Response to organic substance | 27 | 1.11 | 9.48E-15 | NFKBIA,RELB,CCL2,NFKB1,IFNG,IL12B,TNFSF10,IRAK2,CXCL10,HAVCR2,STAT2,LILRB1,OAS2,SRC,TNFSF13B,CD83,DDX58,CCL8,IR |
| GO Process | Positive regulation of interferon-alpha production | 3 | 1.11 | 0.001 | DDX58,IRF7,IFIH1 |
| GO Process | Regulation of immune effector process | 8 | 1.1 | 3.31E-05 | IFNG,IL12B,C3,HAVCR2,LILRB1,NFKBIZ,DDX58,TNF |
| GO Process | Regulation of osteoclast differentiation | 4 | 1.1 | 0.0006 | IFNG,IL12B,LILRB1,TNF |
| GO Process | Positive regulation of osteoclast differentiation | 3 | 1.1 | 0.0011 | IFNG,IL12B,TNF |
| GO Process | Positive regulation of chemokine production | 4 | 1.09 | 0.00063 | IFNG,HAVCR2,OAS1,TNF |
| GO Process | T cell proliferation | 4 | 1.09 | 0.00065 | IL12B,LILRB1,TNFSF13B,IDO1 |
| GO Process | T cell activation | 7 | 1.08 | 7.95E-05 | RELB,IL12B,LILRB1,SLAMF7,TNFSF13B,CD83,IDO1 |
| GO Process | Regulation of myeloid cell differentiation | 6 | 1.08 | 0.00017 | NFKBIA,IFNG,IL12B,LILRB1,IRF7,TNF |
| GO Process | Positive regulation of ERK1 and ERK2 cascade | 6 | 1.08 | 0.00017 | CCL2,HAVCR2,SRC,CCL8,TNF,CCL4 |
| GO Process | Regulation of lymphocyte chemotaxis | 3 | 1.08 | 0.0012 | CCL2,CXCL10,CCL4 |
| GO Process | Positive regulation of calcidiol 1-monooxygenase activity | 2 | 1.08 | 0.0018 | IFNG,TNF |
| GO Process | Negative regulation of T cell activation via T cell recepto | 2 | 1.08 | 0.0018 | HAVCR2,LILRB1 |
| GO Process | Response to stress | 31 | 1.07 | 6.65E-18 | NFKBIA,RELB,CCL2,NFKB1,IFNG,IL12B,C3,IRAK2,CXCL10,HAVCR2,STAT2,LILRB1,NFKBIZ,OAS2,SLAMF7,IFI44L,SRC,CD83,DDX58,C |
| GO Process | Leukocyte differentiation | 8 | 1.07 | 4.18E-05 | RELB,IFNG,IL12B,LILRB1,SRC,TNFSF13B,CD83,TNF |
| GO Process | Negative regulation of innate immune response | 4 | 1.07 | 0.00071 | HAVCR2,STAT2,LILRB1,OAS1 |
| GO Process | Positive regulation of apoptotic process | 9 | 1.06 | 2.09E-05 | CCL2,IFNG,IL12B,TNFSF10,C3,LILRB1,SRC,TNF,IDO1 |
| GO Process | Positive regulation of phagocytosis | 4 | 1.06 | 0.00074 | CCL2,IFNG,C3,TNF |
| GO Process | Regulation of leukocyte differentiation | 7 | 1.05 | 0.0001 | IFNG,IL12B,LILRB1,NFKBIZ,CD83,IRF7,TNF |
| GO Process | Regulation of smooth muscle cell proliferation | 5 | 1.05 | 0.00041 | IFNG,IL12B,SRC,CDKN1A,TNF |
| GO Process | Positive regulation of innate immune response | 5 | 1.05 | 0.00043 | IL12B,HAVCR2,SRC,IRF7,IFI35 |
| GO Process | Cell surface receptor signaling pathway | 21 | 1.04 | 2.09E-10 | NFKBIA,CCL2,IFNG,IL12B,C3,IRAK2,CXCL10,STAT2,LILRB1,NFKBIZ,OAS2,SRC,TNFSF13B,CCL8,IRF7,MX1,OAS1,TNF,CXCL2,CCL4,IF |
| GO Process | Regulation of hemopoiesis | 8 | 1.04 | 5.05E-05 | NFKBIA,IFNG,IL12B,LILRB1,NFKBIZ,CD83,IRF7,TNF |
| GO Process | Positive regulation of interferon-gamma production | 4 | 1.04 | 0.00084 | IL12B,HAVCR2,LILRB1,TNF |
| GO Process | Intracellular receptor signaling pathway | 5 | 1.03 | 0.00049 | NFKBIA,SRC,DDX58,IRF7,IFIH1 |
| GO Process | Myeloid leukocyte activation | 5 | 1.02 | 0.00051 | RELB,IFNG,HAVCR2,IFI35,TNF |
| GO Process | Regulation of interleukin-6 production | 5 | 1.02 | 0.00052 | IFNG,HAVCR2,DDX58,TNF,IFIH1 |
| GO Process | Positive regulation of chronic inflammatory response | 2 | 1.01 | 0.0026 | TNF,IDO1 |
| GO Process | MDA-5 signaling pathway | 2 | 1.01 | 0.0026 | IRF7,IFIH1 |
| GO Process | Regulation of leukocyte mediated immunity | 6 | 1 | 0.0003 | IL12B,C3,HAVCR2,LILRB1,DDX58,TNF |
| GO Process | Negative regulation of type I interferon production | 3 | 1 | 0.0019 | RELB,HAVCR2,LILRB1 |
| GO Process | Positive regulation of response to stimulus | 21 | 0.99 | 4.77E-10 | NFKBIA,CCL2,NFKB1,IFNG,IL12B,TNFSF10,C3,CXCL10,HAVCR2,LILRB1,NFKBIZ,SRC,TNFSF13B,DDX58,CCL8,IRF7,IFI35,TNF,IDO1,CC |
| GO Process | Regulation of lymphocyte proliferation | 6 | 0.99 | 0.00032 | IL12B,HAVCR2,LILRB1,TNFSF13B,CDKN1A,IDO1 |
| GO Process | Cellular response to chemical stimulus | 24 | 0.98 | 1.59E-11 | NFKBIA,RELB,CCL2,NFKB1,IFNG,IL12B,IRAK2,CXCL10,HAVCR2,STAT2,LILRB1,OAS2,SRC,TNFSF13B,DDX58,CCL8,IRF7,MX1,OAS1,T |
| GO Process | Positive regulation of defense response to virus by host | 3 | 0.98 | 0.0021 | IL12B,LILRB1,DDX58 |
| GO Process | Regulation of type III interferon production | 2 | 0.96 | 0.0035 | DDX58,IFIH1 |
| GO Process | Negative regulation of cell population proliferation | 10 | 0.93 | 3.16E-05 | CCL2,IFNG,IL12B,HAVCR2,LILRB1,CCL8,CDKN1A,IFI35,TNF,IDO1 |
| GO Process | Regulation of lymphocyte mediated immunity | 5 | 0.92 | 0.00096 | IL12B,C3,HAVCR2,LILRB1,TNF |
| GO Process | Positive regulation of interleukin-6 production | 4 | 0.91 | 0.0018 | IFNG,DDX58,TNF,IFIH1 |
| GO Process | Regulation of type I interferon-mediated signaling pathwa | 3 | 0.91 | 0.0032 | STAT2,IRF7,OAS1 |
| GO Process | Detection of virus | 2 | 0.91 | 0.0045 | DDX58,IFIH1 |
| GO Process | Positive regulation of NMDA glutamate receptor activity | 2 | 0.91 | 0.0045 | CCL2,IFNG |
| GO Process | Positive regulation of leukocyte differentiation | 5 | 0.9 | 0.0011 | IFNG,IL12B,NFKBIZ,CD83,TNF |
| GO Process | Negative regulation of leukocyte proliferation | 4 | 0.9 | 0.0019 | HAVCR2,LILRB1,CCL8,IDO1 |
| GO Process | Immune system development | 10 | 0.89 | 4.64E-05 | RELB,IFNG,IL12B,HAVCR2,LILRB1,NFKB2,SRC,TNFSF13B,CD83,TNF |
| GO Process | Regulation of T cell proliferation | 5 | 0.88 | 0.0012 | IL12B,HAVCR2,LILRB1,TNFSF13B,IDO1 |
| GO Process | Cellular response to nicotine | 2 | 0.88 | 0.0055 | NFKB1,TNF |
| GO Process | Positive regulation of T-helper 17 cell differentiation | 2 | 0.88 | 0.0055 | IL12B,NFKBIZ |
| GO Process | Negative regulation of gene expression | 11 | 0.87 | 2.71E-05 | RELB,NFKB1,IFNG,IL12B,HAVCR2,LILRB1,CD83,CDKN1A,OAS1,TNF,IDO1 |
| GO Process | Negative regulation of endothelial cell proliferation | 3 | 0.87 | 0.0039 | CCL2,IL12B,TNF |
| GO Process | Monocyte chemotaxis | 3 | 0.87 | 0.0039 | CCL2,CCL8,CCL4 |

| | | | | | |
|---|---|---|---|---|---|
| GO Process | Positive regulation of interleukin-12 production | 3 | 0.87 | 0.0039 | IFNG,IL12B,IDO1 |
| GO Process | Positive regulation of multicellular organismal process | 15 | 0.86 | 1.53E-06 | IFNG,IL12B,C3,HAVCR2,LILRB1,NFKBIZ,OAS2,SRC,CD83,DDX58,IRF7,OAS1,TNF,IDO1,IFIH1 |
| GO Process | Regulation of natural killer cell mediated cytotoxicity dir | 2 | 0.85 | 0.0065 | IL12B,HAVCR2 |
| GO Process | Positive regulation of apoptotic cell clearance | 2 | 0.85 | 0.0065 | CCL2,C3 |
| GO Process | Regulation of natural killer cell chemotaxis | 2 | 0.85 | 0.0065 | CCL2,CCL4 |
| GO Process | Regulation of natural killer cell mediated cytotoxicity | 3 | 0.83 | 0.0048 | IL12B,HAVCR2,LILRB1 |
| GO Process | Negative regulation of lipid localization | 3 | 0.83 | 0.0048 | NFKBIA,NFKB1,TNF |
| GO Process | Regulation of cellular respiration | 3 | 0.83 | 0.005 | IFNG,OAS1,TNF |
| GO Process | Regulation of killing of cells of another organism | 2 | 0.82 | 0.0077 | IFNG,C3 |
| GO Process | Negative regulation of amyloid-beta clearance | 2 | 0.82 | 0.0077 | IFNG,TNF |
| GO Process | Response to exogenous dsRNA | 3 | 0.81 | 0.0055 | NFKBIA,DDX58,IFIH1 |
| GO Process | CD4-positive, alpha-beta T cell differentiation | 3 | 0.81 | 0.0055 | RELB,IL12B,CD83 |
| GO Process | Hematopoietic or lymphoid organ development | 9 | 0.8 | 0.00021 | RELB,IFNG,IL12B,LILRB1,NFKB2,SRC,TNFSF13B,CD83,TNF |
| GO Process | Response to mechanical stimulus | 5 | 0.8 | 0.0021 | NFKBIA,NFKB1,CXCL10,SRC,TNF |
| GO Process | Receptor signaling pathway via JAK-STAT | 3 | 0.8 | 0.0057 | CCL2,IFNG,STAT2 |
| GO Process | Negative regulation of smooth muscle cell proliferation | 3 | 0.79 | 0.0063 | IFNG,IL12B,CDKN1A |
| GO Process | Germinal center formation | 2 | 0.79 | 0.0089 | NFKB2,TNFSF13B |
| GO Process | Regulation of ribonuclease activity | 2 | 0.79 | 0.0089 | OAS2,OAS1 |
| GO Process | Regulation of cell population proliferation | 15 | 0.78 | 5.27E-06 | NFKBIA,CCL2,IFNG,IL12B,CXCL10,HAVCR2,STAT2,LILRB1,SRC,TNFSF13B,CCL8,CDKN1A,IFI35,TNF,IDO1 |
| GO Process | Regulation of mononuclear cell migration | 4 | 0.78 | 0.004 | CCL2,CXCL10,TNF,CCL4 |
| GO Process | Negative regulation of multicellular organismal process | 11 | 0.77 | 8.70E-05 | RELB,NFKB1,IFNG,IL12B,CXCL10,HAVCR2,LILRB1,CD83,OAS1,TNF,IDO1 |
| GO Process | Tumor necrosis factor-mediated signaling pathway | 3 | 0.77 | 0.0071 | NFKBIA,TNFSF13B,TNF |
| GO Process | Negative regulation of leukocyte apoptotic process | 3 | 0.77 | 0.0071 | LILRB1,IRF7,IDO1 |
| GO Process | Positive regulation of podosome assembly | 2 | 0.77 | 0.0102 | SRC,TNF |
| GO Process | Response to oxygen-containing compound | 14 | 0.75 | 1.52E-05 | NFKBIA,CCL2,NFKB1,IL12B,TNFSF10,IRAK2,CXCL10,HAVCR2,STAT2,LILRB1,SRC,TNF,CXCL2,IDO1 |
| GO Process | Regulation of leukocyte migration | 5 | 0.75 | 0.0029 | CCL2,CXCL10,CCL8,TNF,CCL4 |
| GO Process | Myeloid leukocyte differentiation | 4 | 0.75 | 0.005 | RELB,IFNG,SRC,TNF |
| GO Process | Signal transduction | 30 | 0.74 | 2.54E-12 | NFKBIA,RELB,CCL2,NFKB1,IFNG,IL12B,TNFSF10,C3,IRAK2,CXCL10,HAVCR2,STAT2,LILRB1,NFKBIZ,OAS2,NFKB2,SRC,TNFSF13B,CD |
| GO Process | Positive regulation of leukocyte mediated immunity | 4 | 0.74 | 0.0054 | IL12B,C3,DDX58,TNF |
| GO Process | Positive regulation of smooth muscle cell apoptotic proce | 2 | 0.74 | 0.0116 | IFNG,IL12B |
| GO Process | Regulation of vascular associated smooth muscle cell pro | 3 | 0.73 | 0.0087 | SRC,CDKN1A,TNF |
| GO Process | Positive regulation of inflammatory response to antigenic | 2 | 0.72 | 0.0131 | C3,TNF |
| GO Process | Negative regulation of interferon-beta production | 2 | 0.72 | 0.0131 | RELB,LILRB1 |
| GO Process | Positive regulation of lymphocyte proliferation | 4 | 0.71 | 0.0065 | IL12B,HAVCR2,TNFSF13B,CDKN1A |
| GO Process | Positive regulation of response to cytokine stimulus | 3 | 0.71 | 0.0097 | DDX58,IRF7,IFIH1 |
| GO Process | Regulation of neurotransmitter receptor activity | 3 | 0.71 | 0.0097 | CCL2,IFNG,SRC |
| GO Process | Positive regulation of tyrosine phosphorylation of STAT | 3 | 0.71 | 0.01 | IFNG,IL12B,TNF |
| GO Process | Positive regulation of cellular respiration | 2 | 0.71 | 0.0143 | IFNG,OAS1 |
| GO Process | Regulation of programmed cell death | 13 | 0.7 | 6.39E-05 | CCL2,NFKB1,IFNG,IL12B,TNFSF10,C3,CXCL10,LILRB1,SRC,IRF7,CDKN1A,TNF,IDO1 |
| GO Process | Positive regulation of leukocyte migration | 4 | 0.69 | 0.0071 | CXCL10,CCL8,TNF,CCL4 |
| GO Process | Regulation of cytokine-mediated signaling pathway | 4 | 0.69 | 0.0072 | IRAK2,STAT2,IRF7,OAS1 |
| GO Process | Negative regulation of interleukin-17 production | 2 | 0.69 | 0.0159 | IFNG,IL12B |
| GO Process | Positive regulation of granulocyte macrophage colony-sti | 2 | 0.69 | 0.0159 | IL12B,DDX58 |
| GO Process | Positive regulation of macromolecule metabolic process | 23 | 0.68 | 6.58E-08 | NFKBIA,RELB,CCL2,NFKB1,IFNG,IL12B,TNFSF10,C3,CXCL10,HAVCR2,STAT2,LILRB1,OAS2,NFKB2,SRC,CD83,DDX58,IRF7,CDKN1A,C |
| GO Process | Regulation of multicellular organismal process | 19 | 0.67 | 2.28E-06 | NFKBIA,RELB,NFKB1,IFNG,IL12B,C3,CXCL10,HAVCR2,LILRB1,NFKBIZ,OAS2,SRC,CD83,DDX58,IRF7,OAS1,TNF,IDO1,IFIH1 |
| GO Process | Regulation of hydrolase activity | 10 | 0.67 | 0.00044 | CCL2,IFNG,TNFSF10,C3,OAS2,SRC,CCL8,OAS1,TNF,CCL4 |
| GO Process | Killing of cells of another organism | 3 | 0.67 | 0.0123 | CXCL10,CCL8,CXCL2 |
| GO Process | Response to organic cyclic compound | 9 | 0.66 | 0.00084 | NFKBIA,CCL2,NFKB1,CXCL10,SRC,CD83,DDX58,TNF,IFIH1 |
| GO Process | Positive regulation of interleukin-1 production | 3 | 0.66 | 0.0131 | IFNG,HAVCR2,TNF |
| GO Process | Sensory perception of pain | 3 | 0.66 | 0.0135 | CCL2,IL12B,TNF |
| GO Process | Regulation of response to stimulus | 24 | 0.65 | 6.58E-08 | NFKBIA,CCL2,NFKB1,IFNG,IL12B,TNFSF10,C3,IRAK2,CXCL10,HAVCR2,STAT2,LILRB1,NFKBIZ,SRC,TNFSF13B,DDX58,CCL8,IRF7,OAS |
| GO Process | Regulation of lipid localization | 4 | 0.65 | 0.0096 | NFKBIA,NFKB1,C3,TNF |
| GO Process | Negative regulation of T cell proliferation | 3 | 0.65 | 0.014 | HAVCR2,LILRB1,IDO1 |
| GO Process | Positive regulation of carbohydrate metabolic process | 3 | 0.65 | 0.014 | NFKB1,IFNG,SRC |

| | | | | | |
|---|---|---|---|---|---|
| GO Process | Positive regulation of mononuclear cell migration | 3 | 0.65 | 0.014 | CXCL10,TNF,CCL4 |
| GO Process | T cell activation involved in immune response | 3 | 0.65 | 0.0143 | RELB,IL12B,LILRB1 |
| GO Process | Response to UV-B | 2 | 0.65 | 0.0195 | IL12B,CDKN1A |
| GO Process | Astrocyte activation | 2 | 0.65 | 0.0195 | IFNG,TNF |
| GO Process | Positive regulation of membrane protein ectodomain prot | 2 | 0.65 | 0.0195 | IFNG,TNF |
| GO Process | Modulation by host of symbiont process | 3 | 0.64 | 0.0147 | CCL8,CCL4,IFI27 |
| GO Process | Negative regulation of interleukin-12 production | 2 | 0.64 | 0.0213 | NFKB1,LILRB1 |
| GO Process | Positive regulation of nitric-oxide synthase biosynthetic p | 2 | 0.64 | 0.0213 | CCL2,IFNG |
| GO Process | Regulation of chemokine (C-X-C motif) ligand 2 product | 2 | 0.64 | 0.0213 | OAS1,TNF |
| GO Process | Regulation of apoptotic process | 12 | 0.63 | 0.0003 | CCL2,NFKB1,IFNG,IL12B,TNFSF10,C3,CXCL10,LILRB1,SRC,IRF7,TNF,IDO1 |
| GO Process | Regulation of lipid biosynthetic process | 4 | 0.63 | 0.0105 | NFKB1,IFNG,C3,TNF |
| GO Process | Regulation of developmental process | 17 | 0.62 | 2.07E-05 | NFKBIA,CCL2,NFKB1,IFNG,IL12B,C3,CXCL10,STAT2,LILRB1,NFKBIZ,OAS2,SRC,TNFSF13B,CD83,IRF7,CDKN1A,TNF |
| GO Process | Toll-like receptor 4 signaling pathway | 2 | 0.62 | 0.0231 | NFKBIA,OAS1 |
| GO Process | Modulation by host of viral genome replication | 2 | 0.62 | 0.0231 | CCL8,IFI27 |
| GO Process | Negative regulation of natural killer cell mediated cytoto› | 2 | 0.62 | 0.0231 | HAVCR2,LILRB1 |
| GO Process | Negative regulation of type I interferon-mediated signalir | 2 | 0.62 | 0.0231 | STAT2,OAS1 |
| GO Process | Cellular response to exogenous dsRNA | 2 | 0.62 | 0.0231 | DDX58,IFIH1 |
| GO Process | Response to peptide | 6 | 0.61 | 0.0053 | NFKBIA,NFKB1,TNFSF10,STAT2,SRC,TNF |
| GO Process | Regulation of signaling receptor activity | 4 | 0.61 | 0.0119 | CCL2,IFNG,SRC,TNF |
| GO Process | Regulation of tolerance induction | 2 | 0.61 | 0.0249 | HAVCR2,IDO1 |
| GO Process | Intracellular signal transduction | 12 | 0.6 | 0.00041 | NFKBIA,RELB,CCL2,NFKB1,IRAK2,NFKB2,SRC,DDX58,IRF7,CDKN1A,TNF,IFIH1 |
| GO Process | Positive regulation of signal transduction | 12 | 0.6 | 0.00043 | CCL2,NFKB1,IL12B,TNFSF10,C3,HAVCR2,SRC,CCL8,IRF7,IFI35,TNF,CCL4 |
| GO Process | Defense response to bacterium | 5 | 0.6 | 0.0086 | IL12B,HAVCR2,OAS2,OAS1,TNF |
| GO Process | Positive regulation of myeloid leukocyte mediated immur | 2 | 0.6 | 0.0268 | C3,DDX58 |
| GO Process | Negative regulation of lipid storage | 2 | 0.6 | 0.0268 | NFKBIA,TNF |
| GO Process | Positive regulation of amyloid-beta formation | 2 | 0.6 | 0.0268 | IFNG,TNF |
| GO Process | Regulation of transport | 13 | 0.59 | 0.00032 | NFKBIA,CCL2,NFKB1,IFNG,IL12B,C3,CXCL10,LILRB1,OAS2,SRC,TNF,CCL4,IFI27 |
| GO Process | Positive regulation of humoral immune response | 2 | 0.59 | 0.0286 | C3,TNF |
| GO Process | T-helper 1 type immune response | 2 | 0.59 | 0.0286 | RELB,IL12B |
| GO Process | Mononuclear cell differentiation | 5 | 0.58 | 0.0102 | RELB,IL12B,LILRB1,TNFSF13B,CD83 |
| GO Process | Positive regulation of peptidyl-tyrosine phosphorylation | 4 | 0.58 | 0.0149 | IFNG,IL12B,SRC,TNF |
| GO Process | Negative regulation of NF-kappaB transcription factor ac | 3 | 0.58 | 0.0214 | NFKBIA,IRAK2,HAVCR2 |
| GO Process | Regulation of multicellular organismal development | 11 | 0.57 | 0.00096 | NFKBIA,IFNG,IL12B,C3,CXCL10,LILRB1,NFKBIZ,OAS2,CD83,IRF7,TNF |
| GO Process | Positive regulation of intracellular signal transduction | 9 | 0.57 | 0.0023 | CCL2,IL12B,TNFSF10,HAVCR2,SRC,CCL8,IFI35,TNF,CCL4 |
| GO Process | Positive regulation of lipid storage | 2 | 0.56 | 0.0325 | NFKB1,C3 |
| GO Process | Cellular response to angiotensin | 2 | 0.56 | 0.0325 | NFKB1,SRC |
| GO Process | Regulation of molecular function | 18 | 0.54 | 6.39E-05 | NFKBIA,CCL2,NFKB1,IFNG,IL12B,TNFSF10,C3,IRAK2,CXCL10,HAVCR2,OAS2,SRC,DDX58,CCL8,CDKN1A,OAS1,TNF,CCL4 |
| GO Process | Positive regulation of transcription by RNA polymerase I | 10 | 0.54 | 0.0021 | NFKBIA,RELB,NFKB1,CXCL10,STAT2,LILRB1,NFKB2,DDX58,IRF7,TNF |
| GO Process | Regulation of lipid metabolic process | 5 | 0.54 | 0.0136 | NFKB1,IFNG,C3,SRC,TNF |
| GO Process | Response to muscle stretch | 2 | 0.54 | 0.0364 | NFKBIA,NFKB1 |
| GO Process | Positive regulation of macromolecule biosynthetic proces | 13 | 0.53 | 0.00076 | NFKBIA,RELB,CCL2,NFKB1,IFNG,CXCL10,STAT2,LILRB1,NFKB2,SRC,DDX58,IRF7,TNF |
| GO Process | Cellular response to organic cyclic compound | 6 | 0.53 | 0.0101 | CCL2,NFKB1,SRC,DDX58,TNF,IFIH1 |
| GO Process | Gland morphogenesis | 3 | 0.53 | 0.0298 | NFKB1,SRC,TNF |
| GO Process | Positive regulation of acute inflammatory response | 2 | 0.53 | 0.0387 | C3,TNF |
| GO Process | Negative regulation of inflammatory response to antigeni | 2 | 0.53 | 0.0387 | IL12B,SRC |
| GO Process | Response to stimulus | 34 | 0.52 | 1.48E-11 | NFKBIA,RELB,CCL2,NFKB1,IFNG,IL12B,TNFSF10,C3,IRAK2,CXCL10,HAVCR2,STAT2,LILRB1,NFKBIZ,OAS2,SLAMF7,NFKB2,IFI44L,SR |
| GO Process | Negative regulation of myoblast differentiation | 2 | 0.52 | 0.0409 | CXCL10,TNF |
| GO Process | Regulation of gene expression | 24 | 0.51 | 4.44E-06 | NFKBIA,RELB,NFKB1,IFNG,IL12B,C3,IRAK2,CXCL10,HAVCR2,STAT2,LILRB1,NFKBIZ,OAS2,NFKB2,SRC,CD83,DDX58,IRF7,CDKN1A,C |
| GO Process | Regulation of signal transduction | 17 | 0.51 | 0.00019 | NFKBIA,CCL2,NFKB1,IFNG,IL12B,TNFSF10,C3,IRAK2,HAVCR2,STAT2,SRC,CCL8,IRF7,OAS1,IFI35,TNF,CCL4 |
| GO Process | Negative regulation of macromolecule metabolic process | 16 | 0.51 | 0.00034 | RELB,NFKB1,IFNG,IL12B,C3,HAVCR2,LILRB1,NFKB2,SRC,CD83,IRF7,CDKN1A,OAS1,TNF,IDO1,IFI27 |
| GO Process | Cell migration | 8 | 0.51 | 0.0064 | CCL2,IL12B,CXCL10,SRC,CCL8,TNF,CXCL2,CCL4 |
| GO Process | Positive regulation of cell migration | 6 | 0.51 | 0.0121 | IFNG,CXCL10,SRC,CCL8,TNF,CCL4 |
| GO Process | Positive regulation of T cell proliferation | 3 | 0.51 | 0.0325 | IL12B,HAVCR2,TNFSF13B |
| GO Process | Cellular response to interleukin-6 | 2 | 0.51 | 0.0432 | NFKB1,SRC |

| | | | | | |
|---|---|---|---|---|---|
| GO Process | Positive regulation of biological process | 28 | 0.5 | 2.82E-07 | NFKBIA,RELB,CCL2,NFKB1,IFNG,IL12B,TNFSF10,C3,CXCL10,HAVCR2,STAT2,LILRB1,NFKBIZ,OAS2,NFKB2,SRC,TNFSF13B,CD83,DD |
| GO Process | Positive regulation of cellular process | 26 | 0.5 | 1.53E-06 | NFKBIA,RELB,CCL2,NFKB1,IFNG,IL12B,TNFSF10,C3,CXCL10,HAVCR2,STAT2,LILRB1,NFKBIZ,NFKB2,SRC,TNFSF13B,CD83,DDX58,C |
| GO Process | Negative regulation of biological process | 25 | 0.5 | 3.55E-06 | NFKBIA,RELB,CCL2,NFKB1,IFNG,IL12B,C3,CXCL10,HAVCR2,STAT2,LILRB1,OAS2,NFKB2,SRC,CD83,CCL8,IRF7,MX1,CDKN1A,OAS1,I |
| GO Process | Positive regulation of transport | 8 | 0.5 | 0.0068 | NFKBIA,CCL2,IFNG,C3,CXCL10,SRC,TNF,CCL4 |
| GO Process | Regulation of cell migration | 8 | 0.5 | 0.0073 | CCL2,IFNG,CXCL10,SRC,DDX58,CCL8,TNF,CCL4 |
| GO Process | Positive regulation of lipid localization | 3 | 0.5 | 0.0345 | NFKBIA,NFKB1,C3 |
| GO Process | Antimicrobial humoral immune response mediated by ant | 3 | 0.5 | 0.036 | CXCL10,CCL8,CXCL2 |
| GO Process | Extrinsic apoptotic signaling pathway | 3 | 0.5 | 0.036 | IFNG,TNF,IFI27 |
| GO Process | Negative regulation of cell junction assembly | 2 | 0.5 | 0.0456 | SRC,TNF |
| GO Process | Positive regulation of molecular function | 11 | 0.49 | 0.0028 | CCL2,IFNG,IL12B,TNFSF10,IRAK2,SRC,DDX58,CCL8,CDKN1A,TNF,CCL4 |
| GO Process | Negative regulation of molecular function | 9 | 0.49 | 0.0056 | NFKBIA,NFKB1,IFNG,C3,IRAK2,HAVCR2,SRC,CDKN1A,TNF |
| GO Process | Regulation of vesicle-mediated transport | 6 | 0.49 | 0.0143 | CCL2,IFNG,C3,LILRB1,SRC,TNF |
| GO Process | Positive regulation of transmembrane transport | 4 | 0.49 | 0.0274 | CCL2,IFNG,C3,CXCL10 |
| GO Process | Positive regulation of lymphocyte mediated immunity | 3 | 0.49 | 0.0381 | IL12B,C3,TNF |
| GO Process | Microglial cell activation | 2 | 0.49 | 0.0481 | IFNG,TNF |
| GO Process | Regulation of T-helper 1 type immune response | 2 | 0.49 | 0.0481 | IL12B,HAVCR2 |
| GO Process | Regulation of macrophage derived foam cell differentiati | 2 | 0.49 | 0.0481 | NFKBIA,NFKB1 |
| GO Process | Regulation of catalytic activity | 14 | 0.48 | 0.0012 | CCL2,NFKB1,IFNG,IL12B,TNFSF10,C3,CXCL10,OAS2,SRC,CCL8,CDKN1A,OAS1,TNF,CCL4 |
| GO Process | Negative regulation of response to stimulus | 11 | 0.48 | 0.0032 | NFKBIA,CCL2,NFKB1,IL12B,HAVCR2,STAT2,LILRB1,SRC,OAS1,IFI35,TNF |
| GO Process | Positive regulation of cell population proliferation | 8 | 0.48 | 0.0082 | IFNG,IL12B,CXCL10,HAVCR2,SRC,TNFSF13B,CDKN1A,TNF |
| GO Process | Regulation of cytokine production involved in immune re | 3 | 0.48 | 0.0411 | LILRB1,DDX58,TNF |
| GO Process | Positive regulation of catalytic activity | 9 | 0.47 | 0.0072 | CCL2,IFNG,IL12B,TNFSF10,SRC,CCL8,CDKN1A,TNF,CCL4 |
| GO Process | Regulation of metal ion transport | 5 | 0.47 | 0.0245 | CCL2,IFNG,CXCL10,LILRB1,CCL4 |
| GO Process | Negative regulation of immune effector process | 3 | 0.47 | 0.0437 | HAVCR2,LILRB1,TNF |
| GO Process | Positive regulation of hydrolase activity | 6 | 0.46 | 0.0194 | CCL2,IFNG,TNFSF10,CCL8,TNF,CCL4 |
| GO Process | Regulation of DNA biosynthetic process | 3 | 0.46 | 0.0446 | SRC,CDKN1A,TNF |
| GO Process | Positive regulation of transcription, DNA-templated | 11 | 0.45 | 0.0051 | NFKBIA,RELB,NFKB1,CXCL10,STAT2,LILRB1,NFKB2,SRC,DDX58,IRF7,TNF |
| GO Process | Positive regulation of protein kinase activity | 5 | 0.45 | 0.0268 | IFNG,IL12B,SRC,CDKN1A,TNF |
| GO Process | Biological process involved in symbiotic interaction | 4 | 0.45 | 0.0353 | SRC,CCL8,CCL4,IFI27 |
| GO Process | Regulation of macromolecule metabolic process | 26 | 0.44 | 1.54E-05 | NFKBIA,RELB,CCL2,NFKB1,IFNG,IL12B,TNFSF10,C3,IRAK2,CXCL10,HAVCR2,STAT2,LILRB1,NFKBIZ,OAS2,NFKB2,SRC,CD83,DDX58, |
| GO Process | Positive regulation of cellular metabolic process | 16 | 0.44 | 0.0013 | NFKBIA,RELB,NFKB1,IFNG,IL12B,C3,CXCL10,STAT2,LILRB1,NFKB2,SRC,DDX58,IRF7,CDKN1A,OAS1,TNF |
| GO Process | Regulation of calcium ion transport | 4 | 0.44 | 0.039 | CCL2,CXCL10,LILRB1,CCL4 |
| GO Process | Positive regulation of nitrogen compound metabolic proc | 16 | 0.43 | 0.0015 | NFKBIA,RELB,NFKB1,IFNG,IL12B,TNFSF10,C3,CXCL10,STAT2,LILRB1,NFKB2,SRC,DDX58,IRF7,CDKN1A,TNF |
| GO Process | Positive regulation of cellular biosynthetic process | 12 | 0.43 | 0.0052 | NFKBIA,RELB,NFKB1,IFNG,CXCL10,STAT2,LILRB1,NFKB2,SRC,DDX58,IRF7,TNF |
| GO Process | Apoptotic process | 8 | 0.43 | 0.014 | NFKBIA,NFKB1,IFNG,TNFSF10,MX1,CDKN1A,TNF,IFI27 |
| GO Process | Response to nitrogen compound | 8 | 0.43 | 0.0151 | NFKBIA,NFKB1,TNFSF10,STAT2,SRC,DDX58,TNF,IFIH1 |
| GO Process | Regulation of DNA-binding transcription factor activity | 5 | 0.43 | 0.0317 | NFKBIA,IRAK2,HAVCR2,DDX58,TNF |
| GO Process | Rhythmic process | 4 | 0.43 | 0.0409 | RELB,NFKB2,SRC,TNF |
| GO Process | Positive regulation of nucleobase-containing compound r | 12 | 0.42 | 0.0055 | NFKBIA,RELB,NFKB1,IFNG,CXCL10,STAT2,LILRB1,NFKB2,SRC,DDX58,IRF7,TNF |
| GO Process | Lymphocyte differentiation | 4 | 0.42 | 0.0453 | RELB,IL12B,TNFSF13B,CD83 |
| GO Process | Regulation of cell differentiation | 10 | 0.41 | 0.0112 | NFKBIA,NFKB1,IFNG,IL12B,CXCL10,LILRB1,NFKBIZ,CD83,IRF7,TNF |
| GO Process | Positive regulation of developmental process | 9 | 0.41 | 0.0142 | NFKB1,IFNG,IL12B,C3,NFKBIZ,SRC,TNFSF13B,CD83,TNF |
| GO Process | Positive regulation of ion transport | 4 | 0.41 | 0.0485 | CCL2,IFNG,CXCL10,CCL4 |
| GO Process | Negative regulation of cell adhesion | 4 | 0.41 | 0.0499 | HAVCR2,LILRB1,SRC,IDO1 |
| GO Process | Response to abiotic stimulus | 8 | 0.4 | 0.0197 | NFKBIA,RELB,NFKB1,IL12B,CXCL10,SRC,CDKN1A,TNF |
| GO Process | Negative regulation of apoptotic process | 7 | 0.4 | 0.0268 | CCL2,NFKB1,LILRB1,SRC,IRF7,TNF,IDO1 |
| GO Process | Negative regulation of cellular process | 20 | 0.39 | 0.0011 | NFKBIA,RELB,CCL2,NFKB1,IFNG,IL12B,CXCL10,HAVCR2,STAT2,LILRB1,NFKB2,SRC,CCL8,IRF7,CDKN1A,OAS1,IFI35,TNF,IDO1,IFI27 |
| GO Process | Regulation of nucleobase-containing compound metaboli | 18 | 0.39 | 0.0021 | NFKBIA,RELB,NFKB1,IFNG,IRAK2,CXCL10,HAVCR2,STAT2,LILRB1,OAS2,NFKB2,SRC,DDX58,IRF7,CDKN1A,OAS1,TNF,IFI27 |
| GO Process | Regulation of RNA metabolic process | 17 | 0.38 | 0.0029 | NFKBIA,RELB,NFKB1,IFNG,IRAK2,CXCL10,HAVCR2,STAT2,LILRB1,OAS2,NFKB2,SRC,DDX58,IRF7,OAS1,TNF,IFI27 |
| GO Process | Regulation of anatomical structure morphogenesis | 7 | 0.38 | 0.0312 | CCL2,C3,CXCL10,STAT2,SRC,TNFSF13B,TNF |
| GO Process | Regulation of ion transport | 6 | 0.38 | 0.0396 | CCL2,IFNG,CXCL10,LILRB1,TNF,CCL4 |
| GO Process | Regulation of biosynthetic process | 18 | 0.37 | 0.0031 | NFKBIA,RELB,CCL2,NFKB1,IFNG,C3,IRAK2,CXCL10,HAVCR2,STAT2,LILRB1,NFKB2,SRC,DDX58,IRF7,CDKN1A,TNF,IFI27 |
| GO Process | Regulation of intracellular signal transduction | 10 | 0.37 | 0.0198 | NFKBIA,CCL2,IL12B,TNFSF10,HAVCR2,SRC,CCL8,IFI35,TNF,CCL4 |
| GO Process | Cell population proliferation | 6 | 0.37 | 0.0427 | IL12B,LILRB1,SRC,TNFSF13B,TNF,IDO1 |

| | | | | | |
|---|---|---|---|---|---|
| GO Process | Regulation of macromolecule biosynthetic process | 17 | 0.36 | 0.0055 | NFKBIA,RELB,CCL2,NFKB1,IFNG,IRAK2,CXCL10,HAVCR2,STAT2,LILRB1,NFKB2,SRC,DDX58,IRF7,CDKN1A,TNF,IFI27 |
| GO Process | Positive regulation of protein metabolic process | 9 | 0.35 | 0.0303 | NFKBIA,IFNG,IL12B,TNFSF10,C3,SRC,DDX58,CDKN1A,TNF |
| GO Process | Regulation of cellular biosynthetic process | 17 | 0.34 | 0.0084 | NFKBIA,RELB,NFKB1,IFNG,C3,IRAK2,CXCL10,HAVCR2,STAT2,LILRB1,NFKB2,SRC,DDX58,IRF7,CDKN1A,TNF,IFI27 |
| GO Process | Negative regulation of macromolecule biosynthetic proce | 9 | 0.34 | 0.0325 | RELB,NFKB1,IFNG,NFKB2,SRC,IRF7,CDKN1A,TNF,IFI27 |
| GO Process | Regulation of nitrogen compound metabolic process | 21 | 0.33 | 0.0044 | NFKBIA,RELB,NFKB1,IFNG,IL12B,TNFSF10,C3,IRAK2,CXCL10,HAVCR2,STAT2,LILRB1,OAS2,NFKB2,SRC,DDX58,IRF7,CDKN1A,OAS1, |
| GO Process | Regulation of transcription, DNA-templated | 15 | 0.33 | 0.0131 | NFKBIA,RELB,NFKB1,IFNG,IRAK2,CXCL10,HAVCR2,STAT2,LILRB1,NFKB2,SRC,DDX58,IRF7,TNF,IFI27 |
| GO Process | Negative regulation of nucleobase-containing compound | 9 | 0.33 | 0.0364 | RELB,NFKB1,IFNG,NFKB2,SRC,IRF7,CDKN1A,TNF,IFI27 |
| GO Process | Regulation of cellular process | 32 | 0.32 | 0.0001 | NFKBIA,RELB,CCL2,NFKB1,IFNG,IL12B,TNFSF10,C3,IRAK2,CXCL10,HAVCR2,STAT2,LILRB1,NFKBIZ,OAS2,NFKB2,SRC,TNFSF13B,CD |
| GO Process | Negative regulation of cellular biosynthetic process | 9 | 0.32 | 0.0408 | RELB,NFKB1,IFNG,NFKB2,SRC,IRF7,CDKN1A,TNF,IFI27 |
| GO Process | Negative regulation of transcription, DNA-templated | 8 | 0.32 | 0.05 | RELB,NFKB1,IFNG,NFKB2,SRC,IRF7,TNF,IFI27 |
| GO Process | Regulation of primary metabolic process | 21 | 0.31 | 0.0063 | NFKBIA,RELB,NFKB1,IFNG,IL12B,TNFSF10,C3,IRAK2,CXCL10,HAVCR2,STAT2,LILRB1,OAS2,NFKB2,SRC,DDX58,IRF7,CDKN1A,OAS1, |
| GO Process | Animal organ development | 14 | 0.31 | 0.022 | RELB,CCL2,NFKB1,IFNG,IL12B,TNFSF10,CXCL10,LILRB1,NFKB2,SRC,TNFSF13B,CD83,CDKN1A,TNF |
| GO Process | Regulation of transcription by RNA polymerase II | 12 | 0.31 | 0.0313 | NFKBIA,RELB,NFKB1,IFNG,CXCL10,STAT2,LILRB1,NFKB2,DDX58,IRF7,TNF,IFI27 |
| GO Process | Regulation of cellular metabolic process | 20 | 0.3 | 0.0116 | NFKBIA,RELB,NFKB1,IFNG,IL12B,C3,IRAK2,CXCL10,HAVCR2,STAT2,LILRB1,OAS2,NFKB2,SRC,DDX58,IRF7,CDKN1A,OAS1,TNF,IFI27 |
| GO Process | Cellular process | 33 | 0.2 | 0.0286 | NFKBIA,RELB,CCL2,NFKB1,IFNG,IL12B,TNFSF10,C3,IRAK2,CXCL10,HAVCR2,STAT2,LILRB1,NFKBIZ,OAS2,SLAMF7,NFKB2,SRC,TNF |
| GO Function | Cytokine activity | 10 | 1.97 | 3.46E-08 | CCL2,IFNG,IL12B,TNFSF10,CXCL10,TNFSF13B,CCL8,TNF,CXCL2,CCL4 |
| GO Function | Cytokine receptor binding | 10 | 1.83 | 6.75E-08 | CCL2,IFNG,IL12B,TNFSF10,CXCL10,TNFSF13B,CCL8,TNF,CXCL2,CCL4 |
| GO Function | Chemokine activity | 5 | 1.62 | 2.48E-05 | CCL2,CXCL10,CCL8,CXCL2,CCL4 |
| GO Function | Double-stranded RNA binding | 4 | 0.81 | 0.0052 | NFKBIA,RELB,NFKB1,IL12B,TNFSF10,IRAK2,STAT2,LILRB1,SLAMF7,DDX58,MX1,IFI35,TNF,CCL4,IFI27,IFIH1 |
| GO Function | Tumor necrosis factor receptor binding | 3 | 0.74 | 0.0104 | CCL2,IFNG,IL12B,TNFSF10,C3,CXCL10,LILRB1,SRC,TNFSF13B,CCL8,TNF,CXCL2,CCL4 |
| GO Function | 2-5-oligoadenylate synthetase activity | 2 | 0.73 | 0.0142 | CCL2,C3,CXCL10,CCL8,CXCL2,CCL4 |
| GO Function | G protein-coupled receptor binding | 6 | 0.69 | 0.0052 | OAS2,DDX58,OAS1,IFIH1 |
| GO Function | Signaling receptor binding | 13 | 0.64 | 0.00033 | CCL2,IFNG,IL12B,TNFSF10,C3,CXCL10,SRC,TNFSF13B,CCL8,CDKN1A,TNF,CXCL2,CCL4 |
| GO Function | Identical protein binding | 16 | 0.63 | 9.05E-05 | NFKBIA,RELB,CCL2,NFKB1,IFNG,IL12B,TNFSF10,C3,IRAK2,CXCL10,STAT2,LILRB1,SLAMF7,SRC,TNFSF13B,DDX58,CCL8,MX1,CDKN |
| GO Function | CCR chemokine receptor binding | 3 | 0.58 | 0.0266 | TNFSF10,TNFSF13B,TNF |
| GO Function | Molecular function regulator activity | 13 | 0.46 | 0.0052 | OAS2,OAS1 |
| GO Function | Protein binding | 25 | 0.31 | 0.0056 | NFKBIA,RELB,CCL2,NFKB1,IFNG,IL12B,TNFSF10,C3,IRAK2,CXCL10,HAVCR2,STAT2,LILRB1,OAS2,SLAMF7,NFKB2,IFI44L,SRC,TNFSF |
| GO Function | Binding | 32 | 0.22 | 0.0266 | CCL2,CCL8,CCL4 |
| GO Component | I-kappaB/NF-kappaB complex | 3 | 1.37 | 0.00033 | NFKBIA,NFKB1,NFKB2 |
| GO Component | NF-kappaB complex | 3 | 1.33 | 0.0004 | RELB,NFKB1,NFKB2 |
| GO Component | External side of plasma membrane | 6 | 0.46 | 0.0349 | IL12B,CXCL10,LILRB1,SLAMF7,CD83,TNF |
| GO Component | Side of membrane | 7 | 0.42 | 0.0376 | IL12B,CXCL10,LILRB1,SLAMF7,SRC,CD83,TNF |
| GO Component | Cell surface | 8 | 0.37 | 0.0475 | IL12B,C3,CXCL10,HAVCR2,LILRB1,SLAMF7,CD83,TNF |
| STRING clusters | Toll-like Receptor Cascades, and Novel intracellular com | 9 | 2.74 | 1.11E-09 | NFKBIA,RELB,NFKB1,IRAK2,NFKB2,DDX58,IRF7,TNF,IFIH1 |
| STRING clusters | Mixed, incl, Novel intracellular components of RIG-I-like | 8 | 2.85 | 1.44E-09 | NFKBIA,RELB,NFKB1,NFKB2,DDX58,IRF7,TNF,IFIH1 |
| STRING clusters | Mixed, incl, 2-5-oligoadenylate synthetase activity, and In | 5 | 2.46 | 2.63E-07 | OAS2,IFI44L,MX1,OAS1,IFI27 |
| STRING clusters | Mixed, incl, Chemokine-mediated signaling pathway, and | 8 | 1.82 | 4.24E-07 | CCL2,CXCL10,HAVCR2,SLAMF7,CD83,CCL8,CXCL2,CCL4 |
| STRING clusters | Chemokine receptors bind chemokines | 5 | 1.94 | 3.66E-06 | CCL2,CXCL10,CCL8,CXCL2,CCL4 |
| STRING clusters | Mixed, incl, 2-5-oligoadenylate synthetase activity, and T | 4 | 2 | 5.48E-06 | OAS2,IFI44L,MX1,OAS1 |
| STRING clusters | Mixed, incl, TNFR1-induced NFkappaB signaling pathw | 5 | 1.84 | 5.72E-06 | NFKBIA,RELB,NFKB1,NFKB2,TNF |
| STRING clusters | Mixed, incl, NF-kappa-B/Dorsal, and D domain of beta-T | 4 | 1.99 | 5.72E-06 | NFKBIA,RELB,NFKB1,NFKB2 |
| STRING clusters | Mixed, incl, CXC chemokine, and CXC chemokine recep | 3 | 1.25 | 0.00053 | CCL8,CXCL2,CCL4 |
| STRING clusters | TRAF3-dependent IRF activation pathway, and TBD dor | 3 | 1.14 | 0.00099 | DDX58,IRF7,IFIH1 |
| STRING clusters | C-terminal domain of RIG-I, and RIG-I binding | 2 | 0.8 | 0.0092 | DDX58,IFIH1 |
| STRING clusters | Mixed, incl, TLDc domain, and Symbiont intracellular pr | 2 | 0.71 | 0.0152 | IFI44L,MX1 |
| KEGG | Influenza A | 15 | 4.76 | 6.28E-20 | NFKBIA,CCL2,NFKB1,IFNG,IL12B,TNFSF10,CXCL10,STAT2,OAS2,DDX58,IRF7,MX1,OAS1,TNF,IFIH1 |
| KEGG | Hepatitis C | 12 | 3.66 | 8.21E-15 | NFKBIA,NFKB1,IFNG,CXCL10,STAT2,OAS2,DDX58,IRF7,MX1,CDKN1A,OAS1,TNF |
| KEGG | Epstein-Barr virus infection | 12 | 3.27 | 5.41E-14 | NFKBIA,RELB,NFKB1,CXCL10,STAT2,OAS2,NFKB2,DDX58,IRF7,CDKN1A,OAS1,TNF |
| KEGG | Herpes simplex virus 1 infection | 14 | 2.16 | 2.21E-12 | NFKBIA,CCL2,NFKB1,IFNG,IL12B,C3,STAT2,OAS2,SRC,DDX58,IRF7,OAS1,TNF,IFIH1 |
| KEGG | Measles | 10 | 3.15 | 2.92E-12 | NFKBIA,NFKB1,IL12B,STAT2,OAS2,DDX58,IRF7,MX1,OAS1,IFIH1 |
| KEGG | NF-kappa B signaling pathway | 9 | 3.23 | 9.74E-12 | NFKBIA,RELB,NFKB1,NFKB2,TNFSF13B,DDX58,TNF,CXCL2,CCL4 |
| KEGG | RIG-I-like receptor signaling pathway | 8 | 3.32 | 2.97E-11 | NFKBIA,NFKB1,IL12B,CXCL10,DDX58,IRF7,TNF,IFIH1 |
| KEGG | Hepatitis B | 9 | 2.55 | 3.29E-10 | NFKBIA,NFKB1,STAT2,SRC,DDX58,IRF7,CDKN1A,TNF,IFIH1 |
| KEGG | C-type lectin receptor signaling pathway | 8 | 2.78 | 4.00E-10 | NFKBIA,RELB,NFKB1,IL12B,STAT2,NFKB2,SRC,TNF |

| | | | | | |
|---|---|---|---|---|---|
| KEGG | Legionellosis | 7 | 3.08 | 4.00E-10 | NFKBIA,NFKB1,IL12B,C3,NFKB2,TNF,CXCL2 |
| KEGG | NOD-like receptor signaling pathway | 9 | 2.44 | 5.20E-10 | NFKBIA,CCL2,NFKB1,STAT2,OAS2,IRF7,OAS1,TNF,CXCL2 |
| KEGG | Chemokine signaling pathway | 9 | 2.34 | 8.86E-10 | NFKBIA,CCL2,NFKB1,CXCL10,STAT2,SRC,CCL8,CXCL2,CCL4 |
| KEGG | Cytokine-cytokine receptor interaction | 10 | 2.06 | 1.02E-09 | CCL2,IFNG,IL12B,TNFSF10,CXCL10,TNFSF13B,CCL8,TNF,CXCL2,CCL4 |
| KEGG | Osteoclast differentiation | 8 | 2.57 | 1.02E-09 | NFKBIA,RELB,NFKB1,IFNG,STAT2,LILRB1,NFKB2,TNF |
| KEGG | IL-17 signaling pathway | 7 | 2.49 | 6.47E-09 | NFKBIA,CCL2,NFKB1,IFNG,CXCL10,TNF,CXCL2 |
| KEGG | Viral protein interaction with cytokine and cytokine recep | 7 | 2.43 | 8.64E-09 | CCL2,TNFSF10,CXCL10,CCL8,TNF,CXCL2,CCL4 |
| KEGG | Chagas disease | 7 | 2.42 | 8.71E-09 | NFKBIA,CCL2,NFKB1,IFNG,IL12B,C3,TNF |
| KEGG | Toll-like receptor signaling pathway | 7 | 2.39 | 1.01E-08 | NFKBIA,NFKB1,IL12B,CXCL10,IRF7,TNF,CCL4 |
| KEGG | Kaposi sarcoma-associated herpesvirus infection | 8 | 2.01 | 2.03E-08 | NFKBIA,NFKB1,C3,STAT2,SRC,IRF7,CDKN1A,CXCL2 |
| KEGG | Cytosolic DNA-sensing pathway | 6 | 2.45 | 2.80E-08 | NFKBIA,NFKB1,CXCL10,DDX58,IRF7,CCL4 |
| KEGG | Leishmaniasis | 6 | 2.34 | 4.86E-08 | NFKBIA,NFKB1,IFNG,IL12B,C3,TNF |
| KEGG | Tuberculosis | 7 | 1.82 | 2.28E-07 | NFKB1,IFNG,IL12B,C3,IRAK2,SRC,TNF |
| KEGG | Viral carcinogenesis | 7 | 1.71 | 4.33E-07 | NFKBIA,NFKB1,C3,NFKB2,SRC,IRF7,CDKN1A |
| KEGG | TNF signaling pathway | 6 | 1.85 | 6.28E-07 | NFKBIA,CCL2,NFKB1,CXCL10,TNF,CXCL2 |
| KEGG | Human cytomegalovirus infection | 7 | 1.54 | 1.23E-06 | NFKBIA,CCL2,NFKB1,SRC,CDKN1A,TNF,CCL4 |
| KEGG | Rheumatoid arthritis | 5 | 1.68 | 5.11E-06 | CCL2,IFNG,TNFSF13B,TNF,CXCL2 |
| KEGG | African trypanosomiasis | 4 | 1.79 | 7.99E-06 | IFNG,IL12B,TNF,IDO1 |
| KEGG | Amoebiasis | 5 | 1.52 | 1.20E-05 | NFKB1,IFNG,IL12B,TNF,CXCL2 |
| KEGG | Toxoplasmosis | 5 | 1.51 | 1.27E-05 | NFKBIA,NFKB1,IFNG,IL12B,TNF |
| KEGG | Human T-cell leukemia virus 1 infection | 6 | 1.27 | 1.87E-05 | NFKBIA,RELB,NFKB1,NFKB2,CDKN1A,TNF |
| KEGG | Yersinia infection | 5 | 1.36 | 2.86E-05 | NFKBIA,CCL2,NFKB1,SRC,TNF |
| KEGG | Fluid shear stress and atherosclerosis | 5 | 1.33 | 3.34E-05 | CCL2,NFKB1,IFNG,SRC,TNF |
| KEGG | Inflammatory bowel disease | 4 | 1.47 | 4.14E-05 | NFKB1,IFNG,IL12B,TNF |
| KEGG | Epithelial cell signaling in Helicobacter pylori infection | 4 | 1.41 | 5.80E-05 | NFKBIA,NFKB1,SRC,CXCL2 |
| KEGG | Pertussis | 4 | 1.33 | 8.73E-05 | NFKB1,IL12B,C3,TNF |
| KEGG | Th1 and Th2 cell differentiation | 4 | 1.23 | 0.00015 | NFKBIA,NFKB1,IFNG,IL12B |
| KEGG | Pathogenic Escherichia coli infection | 5 | 1.07 | 0.00017 | NFKBIA,NFKB1,TNFSF10,SRC,TNF |
| KEGG | Pathways in cancer | 7 | 0.82 | 0.00022 | NFKBIA,NFKB1,IFNG,IL12B,STAT2,NFKB2,CDKN1A |
| KEGG | T cell receptor signaling pathway | 4 | 1.13 | 0.00026 | NFKBIA,NFKB1,IFNG,TNF |
| KEGG | Allograft rejection | 3 | 1.26 | 0.00029 | IFNG,IL12B,TNF |
| KEGG | Shigellosis | 5 | 0.97 | 0.00031 | NFKBIA,NFKB1,C3,SRC,TNF |
| KEGG | Type I diabetes mellitus | 3 | 1.21 | 0.00038 | IFNG,IL12B,TNF |
| KEGG | Apoptosis | 4 | 0.98 | 0.00064 | NFKBIA,NFKB1,TNFSF10,TNF |
| KEGG | Malaria | 3 | 1.11 | 0.00064 | CCL2,IFNG,TNF |
| KEGG | Necroptosis | 4 | 0.91 | 0.00095 | IFNG,TNFSF10,STAT2,TNF |
| KEGG | JAK-STAT signaling pathway | 4 | 0.87 | 0.0012 | IFNG,IL12B,STAT2,CDKN1A |
| KEGG | Adipocytokine signaling pathway | 3 | 0.93 | 0.0017 | NFKBIA,NFKB1,TNF |
| KEGG | Human papillomavirus infection | 5 | 0.73 | 0.0017 | NFKB1,STAT2,MX1,CDKN1A,TNF |
| KEGG | Chronic myeloid leukemia | 3 | 0.89 | 0.0022 | NFKBIA,NFKB1,CDKN1A |
| KEGG | B cell receptor signaling pathway | 3 | 0.87 | 0.0024 | NFKBIA,NFKB1,LILRB1 |
| KEGG | Proteoglycans in cancer | 4 | 0.76 | 0.0024 | IL12B,SRC,CDKN1A,TNF |
| KEGG | Salmonella infection | 4 | 0.72 | 0.0031 | NFKBIA,NFKB1,TNFSF10,TNF |
| KEGG | PD-L1 expression and PD-1 checkpoint pathway in canc | 3 | 0.82 | 0.0031 | NFKBIA,NFKB1,IFNG |
| KEGG | Small cell lung cancer | 3 | 0.8 | 0.0036 | NFKBIA,NFKB1,CDKN1A |
| KEGG | Systemic lupus erythematosus | 3 | 0.79 | 0.0038 | IFNG,C3,TNF |
| KEGG | AGE-RAGE signaling pathway in diabetic complications | 3 | 0.78 | 0.0039 | CCL2,NFKB1,TNF |
| KEGG | Prostate cancer | 3 | 0.78 | 0.004 | NFKBIA,NFKB1,CDKN1A |
| KEGG | Th17 cell differentiation | 3 | 0.77 | 0.0041 | NFKBIA,NFKB1,IFNG |
| KEGG | HIF-1 signaling pathway | 3 | 0.76 | 0.0044 | NFKB1,IFNG,CDKN1A |
| KEGG | Insulin resistance | 3 | 0.75 | 0.0048 | NFKBIA,NFKB1,TNF |
| KEGG | Neurotrophin signaling pathway | 3 | 0.72 | 0.0056 | NFKBIA,NFKB1,IRAK2 |
| KEGG | Natural killer cell mediated cytotoxicity | 3 | 0.69 | 0.0067 | IFNG,TNFSF10,TNF |
| KEGG | Relaxin signaling pathway | 3 | 0.67 | 0.0075 | NFKBIA,NFKB1,SRC |

| | | | | | |
|---|---|---|---|---|---|
| KEGG | Antifolate resistance | 2 | 0.77 | 0.0077 | NFKB1,TNF |
| KEGG | MAPK signaling pathway | 4 | 0.59 | 0.0077 | RELB,NFKB1,NFKB2,TNF |
| KEGG | Graft-versus-host disease | 2 | 0.72 | 0.0099 | IFNG,TNF |
| KEGG | Bladder cancer | 2 | 0.69 | 0.0119 | SRC,CDKN1A |
| KEGG | Transcriptional misregulation in cancer | 3 | 0.55 | 0.0163 | NFKB1,NFKBIZ,CDKN1A |
| KEGG | Human immunodeficiency virus 1 infection | 3 | 0.48 | 0.0257 | NFKBIA,NFKB1,TNF |
| KEGG | Antigen processing and presentation | 2 | 0.54 | 0.0277 | IFNG,TNF |
| KEGG | Prolactin signaling pathway | 2 | 0.52 | 0.0306 | NFKB1,SRC |
| KEGG | Pancreatic cancer | 2 | 0.51 | 0.0327 | NFKB1,CDKN1A |
| KEGG | ErbB signaling pathway | 2 | 0.47 | 0.0414 | SRC,CDKN1A |
| Reactome | Cytokine Signaling in Immune system | 24 | 3.1 | 6.30E-24 | NFKBIA,RELB,CCL2,NFKB1,IFNG,IL12B,IRAK2,CXCL10,HAVCR2,STAT2,OAS2,NFKB2,SRC,TNFSF13B,DDX58,IRF7,MX1,CDKN1A,OAS |
| Reactome | Interferon alpha/beta signaling | 7 | 2.44 | 3.22E-08 | STAT2,OAS2,IRF7,MX1,OAS1,IFI35,IFI27 |
| Reactome | Interleukin-10 signaling | 6 | 2.41 | 1.06E-07 | CCL2,IL12B,CXCL10,TNF,CXCL2,CCL4 |
| Reactome | TRAF6 mediated NF-kB activation | 5 | 2.3 | 5.03E-07 | NFKBIA,NFKB1,NFKB2,DDX58,IFIH1 |
| Reactome | Signaling by Interleukins | 14 | 2.16 | 9.80E-12 | NFKBIA,CCL2,NFKB1,IFNG,IL12B,IRAK2,CXCL10,HAVCR2,STAT2,NFKB2,CDKN1A,TNF,CXCL2,CCL4 |
| Reactome | Interferon Signaling | 9 | 2.01 | 3.22E-08 | IFNG,STAT2,OAS2,DDX58,IRF7,MX1,OAS1,IFI35,IFI27 |
| Reactome | SARS-CoV-1 activates/modulates innate immune respons | 5 | 1.94 | 3.29E-06 | NFKBIA,NFKB1,IRAK2,DDX58,IFIH1 |
| Reactome | DDX58/IFIH1-mediated induction of interferon-alpha/be | 6 | 1.86 | 2.00E-06 | NFKBIA,NFKB1,NFKB2,DDX58,IRF7,IFIH1 |
| Reactome | Immune System | 28 | 1.55 | 1.13E-19 | NFKBIA,RELB,CCL2,NFKB1,IFNG,IL12B,C3,IRAK2,CXCL10,HAVCR2,STAT2,LILRB1,OAS2,SLAMF7,NFKB2,SRC,TNFSF13B,DDX58,IRF7 |
| Reactome | DEx/H-box helicases activate type I IFN and inflammato | 3 | 1.51 | 0.00013 | NFKB1,NFKB2,IRF7 |
| Reactome | IkBA variant leads to EDA-ID | 3 | 1.51 | 0.00013 | NFKBIA,NFKB1,NFKB2 |
| Reactome | OAS antiviral response | 3 | 1.47 | 0.00015 | OAS2,DDX58,OAS1 |
| Reactome | TAK1-dependent IKK and NF-kappa-B activation | 4 | 1.36 | 0.00015 | NFKBIA,NFKB1,IRAK2,NFKB2 |
| Reactome | TRAF3-dependent IRF activation pathway | 3 | 1.31 | 0.00034 | DDX58,IRF7,IFIH1 |
| Reactome | RIP-mediated NFkB activation via ZBP1 | 3 | 1.28 | 0.00041 | NFKBIA,NFKB1,NFKB2 |
| Reactome | CLEC7A (Dectin-1) signaling | 5 | 1.26 | 0.00014 | NFKBIA,RELB,NFKB1,NFKB2,SRC |
| Reactome | Toll Like Receptor 3 (TLR3) Cascade | 5 | 1.25 | 0.00015 | NFKBIA,NFKB1,IRAK2,NFKB2,IRF7 |
| Reactome | MyD88 dependent cascade initiated on endosome | 5 | 1.25 | 0.00015 | NFKBIA,NFKB1,IRAK2,NFKB2,IRF7 |
| Reactome | TRIF(TICAM1)-mediated TLR4 signaling | 5 | 1.24 | 0.00015 | NFKBIA,NFKB1,IRAK2,NFKB2,IRF7 |
| Reactome | Chemokine receptors bind chemokines | 4 | 1.21 | 0.00034 | CCL2,CXCL10,CXCL2,CCL4 |
| Reactome | Cytosolic sensors of pathogen-associated DNA | 4 | 1.17 | 0.00041 | NFKBIA,NFKB1,NFKB2,IRF7 |
| Reactome | SARS-CoV-2 activates/modulates innate and adaptive im | 5 | 1.14 | 0.00025 | IRAK2,STAT2,DDX58,IRF7,IFIH1 |
| Reactome | Purinergic signaling in leishmaniasis infection | 3 | 1.1 | 0.0011 | NFKB1,C3,NFKB2 |
| Reactome | TRAF6 mediated IRF7 activation | 3 | 1.06 | 0.0013 | DDX58,IRF7,IFIH1 |
| Reactome | Antiviral mechanism by IFN-stimulated genes | 4 | 1.02 | 0.00092 | OAS2,DDX58,MX1,OAS1 |
| Reactome | Interferon gamma signaling | 4 | 0.98 | 0.0012 | IFNG,OAS2,IRF7,OAS1 |
| Reactome | TNFR2 non-canonical NF-kB pathway | 4 | 0.94 | 0.0014 | RELB,NFKB2,TNFSF13B,TNF |
| Reactome | Interleukin-4 and Interleukin-13 signaling | 4 | 0.9 | 0.0018 | CCL2,IL12B,CDKN1A,TNF |
| Reactome | Peptide ligand-binding receptors | 5 | 0.87 | 0.0013 | CCL2,C3,CXCL10,CXCL2,CCL4 |
| Reactome | Interleukin-1 processing | 2 | 0.86 | 0.006 | NFKB1,NFKB2 |
| Reactome | SARS-CoV Infections | 7 | 0.85 | 0.00044 | NFKBIA,NFKB1,IRAK2,STAT2,DDX58,IRF7,IFIH1 |
| Reactome | SUMOylation of immune response proteins | 2 | 0.83 | 0.007 | NFKBIA,NFKB2 |
| Reactome | Nucleotide-binding domain, leucine rich repeat containin | 3 | 0.8 | 0.0055 | NFKB1,IRAK2,NFKB2 |
| Reactome | NF-kB is activated and signals survival | 2 | 0.76 | 0.0105 | NFKBIA,NFKB1 |
| Reactome | NF-kB activation through FADD/RIP-1 pathway mediate | 2 | 0.76 | 0.0105 | DDX58,IFIH1 |
| Reactome | Infectious disease | 10 | 0.73 | 0.00024 | NFKBIA,NFKB1,C3,IRAK2,STAT2,NFKB2,SRC,DDX58,IRF7,IFIH1 |
| Reactome | Death Receptor Signaling | 4 | 0.72 | 0.0057 | NFKBIA,NFKB1,TNFSF10,TNF |
| Reactome | The NLRP3 inflammasome | 2 | 0.7 | 0.0148 | NFKB1,NFKB2 |
| Reactome | Leishmania infection | 4 | 0.68 | 0.0071 | NFKB1,C3,NFKB2,SRC |
| Reactome | Innate Immune System | 10 | 0.66 | 0.00051 | NFKBIA,RELB,NFKB1,C3,IRAK2,NFKB2,SRC,DDX58,IRF7,IFIH1 |
| Reactome | G alpha (i) signalling events | 5 | 0.62 | 0.007 | C3,CXCL10,SRC,CXCL2,CCL4 |
| Reactome | CD209 (DC-SIGN) signaling | 2 | 0.62 | 0.0236 | RELB,NFKB1 |
| Reactome | Disease | 11 | 0.46 | 0.0037 | NFKBIA,NFKB1,C3,IRAK2,STAT2,NFKB2,SRC,DDX58,IRF7,CDKN1A,IFIH1 |
| Reactome | Immunoregulatory interactions between a Lymphoid and | 3 | 0.45 | 0.0481 | C3,LILRB1,SLAMF7 |

| | | | | | |
|---|---|---|---|---|---|
| Reactome | Signaling by GPCR | 6 | 0.39 | 0.0364 | CCL2,C3,CXCL10,SRC,CXCL2,CCL4 |
| WikiPathways | Network map of SARS-CoV-2 signaling pathway | 15 | 3.95 | 9.12E-18 | CCL2,IFNG,TNFSF10,CXCL10,OAS2,NFKB2,IFI44L,DDX58,CCL8,MX1,TNF,CXCL2,CCL4,IFI27,IFIH1 |
| WikiPathways | SARS-CoV-2 innate immunity evasion and cell-specific i | 11 | 4.93 | 1.58E-16 | CCL2,NFKB1,CXCL10,HAVCR2,STAT2,DDX58,IRF7,MX1,TNF,CXCL2,CCL4 |
| WikiPathways | Measles virus infection | 11 | 3.5 | 1.60E-13 | NFKBIA,NFKB1,IL12B,STAT2,OAS2,NFKB2,DDX58,IRF7,MX1,OAS1,IFIH1 |
| WikiPathways | Non-genomic actions of 1,25 dihydroxyvitamin D3 | 9 | 3.62 | 2.48E-12 | RELB,CCL2,NFKB1,IFNG,STAT2,OAS2,NFKB2,IFI44L,TNF |
| WikiPathways | Overview of proinflammatory and profibrotic mediators | 10 | 3.23 | 3.08E-12 | CCL2,NFKB1,IFNG,IL12B,CXCL10,TNFSF13B,CCL8,TNF,CXCL2,CCL4 |
| WikiPathways | Novel intracellular components of RIG-I-like receptor pat | 8 | 3.44 | 2.53E-11 | NFKBIA,NFKB1,IFNG,CXCL10,DDX58,IRF7,TNF,IFIH1 |
| WikiPathways | Host-pathogen interaction of human coronaviruses - inter | 7 | 3.6 | 4.66E-11 | NFKBIA,NFKB1,STAT2,OAS2,DDX58,OAS1,IFIH1 |
| WikiPathways | miRNA role in immune response in sepsis | 7 | 3.47 | 8.34E-11 | NFKBIA,RELB,NFKB1,NFKB2,IRF7,TNF,CCL4 |
| WikiPathways | IL-18 signaling pathway | 11 | 2.33 | 8.34E-11 | NFKBIA,CCL2,NFKB1,IFNG,IL12B,NFKBIZ,NFKB2,CD83,TNF,CXCL2,CCL4 |
| WikiPathways | Fibrin complement receptor 3 signaling pathway | 7 | 3.31 | 1.73E-10 | CCL2,NFKB1,IL12B,IRAK2,CXCL10,SRC,TNF |
| WikiPathways | Type I interferon induction and signaling during SARS-C | 6 | 3.03 | 2.61E-09 | STAT2,OAS2,DDX58,IRF7,OAS1,IFIH1 |
| WikiPathways | Prostaglandin signaling | 6 | 3 | 2.84E-09 | CCL2,NFKB1,IFNG,CXCL10,IRF7,TNF |
| WikiPathways | TNF-related weak inducer of apoptosis (TWEAK) signal | 6 | 2.72 | 1.15E-08 | NFKBIA,RELB,CCL2,NFKB1,NFKB2,TNF |
| WikiPathways | Hepatitis B infection | 8 | 2.2 | 1.20E-08 | NFKB1,STAT2,SRC,DDX58,IRF7,CDKN1A,TNF,IFIH1 |
| WikiPathways | Toll-like receptor signaling pathway | 7 | 2.27 | 3.20E-08 | NFKBIA,NFKB1,IL12B,CXCL10,IRF7,TNF,CCL4 |
| WikiPathways | Cytosolic DNA-sensing pathway | 6 | 2.15 | 2.04E-07 | NFKBIA,NFKB1,CXCL10,DDX58,IRF7,CCL4 |
| WikiPathways | Mitochondrial immune response to SARS-CoV-2 | 5 | 2.33 | 2.47E-07 | NFKB1,NFKB2,DDX58,IRF7,IFIH1 |
| WikiPathways | Photodynamic therapy-induced NF-kB survival signaling | 5 | 2.31 | 2.68E-07 | RELB,NFKB1,NFKB2,TNF,CXCL2 |
| WikiPathways | T-cell activation SARS-CoV-2 | 6 | 1.98 | 4.93E-07 | NFKBIA,NFKB1,IFNG,IL12B,CDKN1A,TNF |
| WikiPathways | Thymic stromal lymphopoietin (TSLP) signaling pathway | 5 | 2.04 | 1.07E-06 | NFKBIA,RELB,NFKB1,NFKB2,SRC |
| WikiPathways | Vitamin B12 metabolism | 5 | 1.99 | 1.36E-06 | CCL2,NFKB1,IFNG,NFKB2,TNF |
| WikiPathways | RANKL/RANK signaling pathway | 5 | 1.92 | 1.95E-06 | NFKBIA,RELB,NFKB1,NFKB2,SRC |
| WikiPathways | Spinal cord injury | 6 | 1.73 | 1.95E-06 | CCL2,IFNG,CXCL10,TNFSF13B,TNF,CXCL2 |
| WikiPathways | Lung fibrosis | 5 | 1.81 | 3.43E-06 | CCL2,IL12B,TNF,CXCL2,CCL4 |
| WikiPathways | Ebola virus infection in host | 6 | 1.62 | 3.43E-06 | RELB,NFKB1,HAVCR2,NFKB2,DDX58,IRF7 |
| WikiPathways | Ebstein-Barr virus LMP1 signaling | 4 | 1.98 | 3.76E-06 | NFKBIA,NFKB1,NFKB2,TNF |
| WikiPathways | Immune response to tuberculosis | 4 | 1.98 | 3.76E-06 | STAT2,MX1,OAS1,IFI35 |
| WikiPathways | Kynurenine pathway and links to cell senescence | 4 | 1.98 | 3.76E-06 | IFNG,CDKN1A,TNF,IDO1 |
| WikiPathways | Folate metabolism | 5 | 1.78 | 3.88E-06 | CCL2,NFKB1,IFNG,NFKB2,TNF |
| WikiPathways | LDL- influence on CD14 and TLR4 | 4 | 1.97 | 3.88E-06 | RELB,CCL2,NFKB1,NFKB2 |
| WikiPathways | Cytokines and inflammatory response | 4 | 1.92 | 4.91E-06 | IFNG,IL12B,TNF,CXCL2 |
| WikiPathways | Extrafollicular and follicular B cell activation by SARS-C | 5 | 1.73 | 4.93E-06 | IFNG,C3,SLAMF7,TNFSF13B,TNF |
| WikiPathways | Interactions of natural killer cells in pancreatic cancer | 4 | 1.9 | 5.29E-06 | CCL2,IFNG,TNF,CCL4 |
| WikiPathways | TLR4 signaling and tolerance | 4 | 1.88 | 5.86E-06 | NFKBIA,NFKB1,IRF7,TNF |
| WikiPathways | Interactions between immune cells and microRNAs in tu | 4 | 1.88 | 5.86E-06 | CCL2,NFKB1,CXCL10,NFKB2 |
| WikiPathways | Toll-like receptor signaling related to MyD88 | 4 | 1.82 | 8.03E-06 | RELB,NFKB1,NFKB2,IRF7 |
| WikiPathways | Antiviral and anti-inflammatory effects of Nrf2 on SARS | 4 | 1.82 | 8.03E-06 | NFKBIA,CCL2,NFKB1,TNF |
| WikiPathways | Apoptosis | 5 | 1.63 | 8.12E-06 | NFKBIA,NFKB1,TNFSF10,IRF7,TNF |
| WikiPathways | Selenium micronutrient network | 5 | 1.62 | 8.37E-06 | CCL2,NFKB1,IFNG,NFKB2,TNF |
| WikiPathways | Acute viral myocarditis | 5 | 1.61 | 8.63E-06 | IFNG,IL12B,NFKB2,SRC,TNF |
| WikiPathways | Resistin as a regulator of inflammation | 4 | 1.79 | 8.87E-06 | NFKBIA,NFKB1,IL12B,TNF |
| WikiPathways | Signal transduction through IL1R | 4 | 1.79 | 8.87E-06 | NFKBIA,NFKB1,IRAK2,TNF |
| WikiPathways | TNF-alpha signaling pathway | 5 | 1.55 | 1.17E-05 | NFKBIA,CCL2,NFKB1,NFKB2,TNF |
| WikiPathways | Type II interferon signaling | 4 | 1.72 | 1.26E-05 | IFNG,CXCL10,STAT2,OAS1 |
| WikiPathways | Immune infiltration in pancreatic cancer | 4 | 1.71 | 1.36E-05 | CCL2,IFNG,IL12B,TNF |
| WikiPathways | Burn wound healing | 5 | 1.49 | 1.62E-05 | NFKBIA,CCL2,NFKB1,NFKBIZ,TNF |
| WikiPathways | Altered glycosylation of MUC1 in tumor microenvironme | 3 | 1.82 | 1.69E-05 | NFKBIA,NFKB1,TNF |
| WikiPathways | mRNA vaccine activation of dendritic cell and induction | 3 | 1.78 | 2.15E-05 | DDX58,IRF7,IFIH1 |
| WikiPathways | Aryl hydrocarbon receptor pathway | 4 | 1.6 | 2.35E-05 | NFKB1,SRC,CDKN1A,TNF |
| WikiPathways | Aryl hydrocarbon receptor pathway | 4 | 1.6 | 2.35E-05 | IFNG,IL12B,SRC,TNF |
| WikiPathways | NO/cGMP/PKG mediated neuroprotection | 4 | 1.59 | 2.45E-05 | NFKBIA,NFKB1,IFNG,TNF |
| WikiPathways | Interleukin-1 (IL-1) structural pathway | 4 | 1.56 | 2.82E-05 | NFKBIA,NFKB1,IRAK2,IRF7 |
| WikiPathways | Photodynamic therapy-induced AP-1 survival signaling | 4 | 1.54 | 3.22E-05 | IFNG,TNFSF10,CDKN1A,TNF |

| | | | | | |
|---|---|---|---|---|---|
| WikiPathways | NRP1-triggered signaling pathways in pancreatic cancer | 4 | 1.51 | 3.66E-05 | RELB,NFKB1,NFKB2,SRC |
| WikiPathways | IL-1 signaling pathway | 4 | 1.48 | 4.44E-05 | NFKBIA,CCL2,NFKB1,IRAK2 |
| WikiPathways | COVID-19 adverse outcome pathway | 3 | 1.6 | 5.23E-05 | CCL2,CXCL10,TNF |
| WikiPathways | Pathways of nucleic acid metabolism and innate immune | 3 | 1.57 | 6.09E-05 | DDX58,OAS1,IFIH1 |
| WikiPathways | Neuroinflammation and glutamatergic signaling | 5 | 1.25 | 6.31E-05 | NFKB1,IFNG,IL12B,NFKB2,TNF |
| WikiPathways | T-cell antigen receptor (TCR) pathway during Staphyloc | 4 | 1.4 | 6.50E-05 | NFKBIA,NFKB1,IFNG,TNF |
| WikiPathways | Oncostatin M signaling pathway | 4 | 1.38 | 7.20E-05 | NFKBIA,CCL2,NFKB1,SRC |
| WikiPathways | Ulcerative colitis signaling | 3 | 1.53 | 7.79E-05 | NFKB1,IFNG,TNF |
| WikiPathways | Vitamin D in inflammatory diseases | 3 | 1.46 | 0.00011 | NFKBIA,NFKB1,TNF |
| WikiPathways | Malignant pleural mesothelioma | 7 | 0.92 | 0.00011 | CCL2,NFKB1,CXCL10,SRC,CDKN1A,IDO1,CCL4 |
| WikiPathways | Chemokine signaling pathway | 5 | 1.13 | 0.00012 | NFKBIA,NFKB1,CXCL10,STAT2,CCL4 |
| WikiPathways | FGF23 signaling in hypophosphatemic rickets and relatec | 3 | 1.41 | 0.00014 | NFKB1,NFKB2,CDKN1A |
| WikiPathways | IL1 and megakaryocytes in obesity | 3 | 1.39 | 0.00016 | CCL2,NFKB1,IFNG |
| WikiPathways | Nuclear receptors meta-pathway | 6 | 0.95 | 0.00018 | CCL2,IFNG,IL12B,NFKB2,SRC,TNF |
| WikiPathways | Allograft rejection | 4 | 1.19 | 0.00021 | IFNG,IL12B,C3,TNF |
| WikiPathways | Selective expression of chemokine receptors during T-cel | 3 | 1.3 | 0.00025 | IFNG,IL12B,CCL4 |
| WikiPathways | miRNA regulation of prostate cancer signaling pathways | 3 | 1.27 | 0.0003 | NFKBIA,NFKB1,CDKN1A |
| WikiPathways | SARS coronavirus and innate immunity | 3 | 1.27 | 0.0003 | STAT2,DDX58,IFIH1 |
| WikiPathways | Development and heterogeneity of the ILC family | 3 | 1.25 | 0.00032 | IFNG,IL12B,TNF |
| WikiPathways | Neovascularisation processes | 3 | 1.18 | 0.00047 | RELB,NFKB1,NFKB2 |
| WikiPathways | Gastrin signaling pathway | 4 | 1.03 | 0.0005 | NFKBIA,NFKB1,SRC,CDKN1A |
| WikiPathways | CKAP4 signaling pathway map | 4 | 1.03 | 0.00051 | NFKB1,CXCL10,SRC,TNF |
| WikiPathways | Oxidative damage response | 3 | 1.16 | 0.00053 | NFKB1,CDKN1A,TNF |
| WikiPathways | PDGF pathway | 3 | 1.15 | 0.00056 | NFKBIA,NFKB1,SRC |
| WikiPathways | Inflammatory bowel disease signaling | 3 | 1.11 | 0.00068 | NFKB1,IFNG,TNF |
| WikiPathways | Netrin-UNC5B signaling pathway | 3 | 1.02 | 0.0011 | CCL2,SRC,TNF |
| WikiPathways | Apoptosis-related network due to altered Notch3 in ovari | 3 | 1.01 | 0.0012 | NFKB1,CDKN1A,TNF |
| WikiPathways | Canonical NF-kB pathway | 2 | 1.12 | 0.0012 | NFKBIA,NFKB1 |
| WikiPathways | TGF-beta receptor signaling | 3 | 1.01 | 0.0012 | NFKB1,IFNG,TNF |
| WikiPathways | Supression of HMGB1 mediated inflammation by THBD | 2 | 1.08 | 0.0015 | NFKBIA,NFKB1 |
| WikiPathways | TGF-beta receptor signaling in skeletal dysplasias | 3 | 0.97 | 0.0015 | NFKB1,IFNG,TNF |
| WikiPathways | Notch signaling pathway | 3 | 0.95 | 0.0017 | NFKB1,SRC,CDKN1A |
| WikiPathways | AGE/RAGE pathway | 3 | 0.92 | 0.002 | NFKBIA,NFKB1,SRC |
| WikiPathways | Inclusion body myositis | 2 | 1.03 | 0.002 | NFKB1,NFKB2 |
| WikiPathways | RAC1/PAK1/p38/MMP2 pathway | 3 | 0.9 | 0.0022 | NFKBIA,NFKB1,SRC |
| WikiPathways | MAPK and NFkB signaling pathways inhibited by Yersir | 2 | 1 | 0.0023 | NFKBIA,NFKB1 |
| WikiPathways | Th17 cell differentiation pathway | 3 | 0.89 | 0.0023 | NFKBIA,NFKB1,IFNG |
| WikiPathways | Head and neck squamous cell carcinoma | 3 | 0.88 | 0.0025 | NFKB1,NFKB2,CDKN1A |
| WikiPathways | Prolactin signaling pathway | 3 | 0.85 | 0.0029 | NFKBIA,NFKB1,SRC |
| WikiPathways | Quercetin and Nf-kB / AP-1 induced apoptosis | 2 | 0.94 | 0.0032 | NFKBIA,NFKB1 |
| WikiPathways | Cells and molecules involved in local acute inflammatory | 2 | 0.9 | 0.004 | C3,TNF |
| WikiPathways | Apoptosis modulation and signaling | 3 | 0.78 | 0.0043 | NFKBIA,NFKB1,TNFSF10 |
| WikiPathways | p53 transcriptional gene network | 3 | 0.78 | 0.0044 | CCL2,CDKN1A,TNF |
| WikiPathways | T-cell receptor signaling pathway | 3 | 0.78 | 0.0044 | NFKBIA,NFKB1,CD83 |
| WikiPathways | TP53 network | 2 | 0.87 | 0.0047 | TNFSF10,CDKN1A |
| WikiPathways | LTF danger signal response pathway | 2 | 0.87 | 0.0047 | NFKB1,TNF |
| WikiPathways | Small cell lung cancer | 3 | 0.76 | 0.005 | NFKBIA,NFKB1,CDKN1A |
| WikiPathways | 17q12 copy number variation syndrome | 3 | 0.74 | 0.0054 | CCL2,CCL8,CCL4 |
| WikiPathways | STING pathway in Kawasaki-like disease and COVID-1 | 2 | 0.84 | 0.0055 | NFKBIA,NFKB1 |
| WikiPathways | MAPK signaling pathway | 4 | 0.63 | 0.0064 | RELB,NFKB1,NFKB2,TNF |
| WikiPathways | Cancer immunotherapy by PD-1 blockade | 2 | 0.81 | 0.0064 | NFKB1,IFNG |
| WikiPathways | DNA damage response (only ATM dependent) | 3 | 0.7 | 0.007 | NFKB1,NFKB2,CDKN1A |
| WikiPathways | Cell interactions of the pancreatic cancer microenvironm | 2 | 0.79 | 0.0073 | CCL2,CXCL10 |
| WikiPathways | SARS-CoV-2 mitochondrial chronic oxidative stress and | 2 | 0.77 | 0.0083 | NFKB1,TNF |

| | | | | | |
|---|---|---|---|---|---|
| WikiPathways | T cell receptor and co-stimulatory signaling | 2 | 0.75 | 0.0088 | NFKBIA,NFKB1 |
| WikiPathways | RAS and bradykinin pathways in COVID-19 | 2 | 0.75 | 0.0088 | NFKB1,TNF |
| WikiPathways | Dravet syndrome | 2 | 0.75 | 0.0088 | NFKB1,TNF |
| WikiPathways | Oligodendrocyte specification and differentiation, leadin | 2 | 0.74 | 0.0097 | TNF,CXCL2 |
| WikiPathways | Adipogenesis | 3 | 0.63 | 0.0107 | STAT2,CDKN1A,TNF |
| WikiPathways | Initiation of transcription and translation elongation at th | 2 | 0.72 | 0.0108 | NFKBIA,NFKB1 |
| WikiPathways | Overview of interferons-mediated signaling pathway | 2 | 0.68 | 0.0133 | IFNG,STAT2 |
| WikiPathways | Brain-derived neurotrophic factor (BDNF) signaling path | 3 | 0.59 | 0.0136 | NFKBIA,NFKB1,SRC |
| WikiPathways | Bladder cancer | 2 | 0.65 | 0.0153 | SRC,CDKN1A |
| WikiPathways | ATM signaling pathway | 2 | 0.65 | 0.0159 | NFKBIA,CDKN1A |
| WikiPathways | Nonalcoholic fatty liver disease | 3 | 0.56 | 0.0163 | CCL2,NFKB1,TNF |
| WikiPathways | Sudden infant death syndrome (SIDS) susceptibility path | 3 | 0.56 | 0.0163 | NFKB1,NFKB2,TNF |
| WikiPathways | T cell modulation in pancreatic cancer | 2 | 0.61 | 0.0193 | HAVCR2,IDO1 |
| WikiPathways | Hepatitis C and hepatocellular carcinoma | 2 | 0.59 | 0.0216 | NFKB1,CDKN1A |
| WikiPathways | TROP2 regulatory signaling | 2 | 0.59 | 0.0223 | NFKB1,SRC |
| WikiPathways | Cardiac hypertrophic response | 2 | 0.56 | 0.0266 | NFKB1,TNF |
| WikiPathways | IL-4 signaling pathway | 2 | 0.56 | 0.0266 | NFKBIA,NFKB1 |
| WikiPathways | MET in type 1 papillary renal cell carcinoma | 2 | 0.54 | 0.0299 | SRC,CDKN1A |
| WikiPathways | Modulators of TCR signaling and T cell activation | 2 | 0.52 | 0.0327 | NFKBIA,NFKB1 |
| WikiPathways | Physico-chemical features and toxicity-associated pathwa | 2 | 0.5 | 0.0365 | SRC,CDKN1A |
| WikiPathways | VEGFA-VEGFR2 signaling | 4 | 0.39 | 0.0381 | NFKBIA,CCL2,NFKB1,SRC |
| WikiPathways | Glucocorticoid receptor pathway | 2 | 0.48 | 0.0413 | CCL2,NFKB2 |
| WikiPathways | Leptin signaling pathway | 2 | 0.46 | 0.0479 | NFKB1,SRC |
| Monarch | Autoimmunity | 5 | 0.55 | 0.0298 | NFKB1,C3,HAVCR2,NFKB2,IFIH1 |
| DISEASES | Primary immunodeficiency disease | 12 | 1.58 | 6.09E-08 | NFKBIA,CCL2,NFKB1,IFNG,IL12B,C3,STAT2,NFKB2,IRF7,OAS1,TNF,IFIH1 |
| DISEASES | Microphthalmia with limb anomalies | 5 | 2.72 | 7.61E-08 | OAS2,DDX58,MX1,OAS1,IFIH1 |
| DISEASES | Autoimmune disease | 9 | 1.4 | 3.44E-06 | CCL2,NFKB1,IFNG,IL12B,NFKB2,IRF7,OAS1,TNF,IFIH1 |
| DISEASES | Respiratory failure | 4 | 1.66 | 3.96E-05 | CCL2,IFNG,CXCL10,TNF |
| DISEASES | Inflammatory bowel disease | 5 | 1.48 | 5.60E-05 | CCL2,NFKB1,IFNG,IL12B,TNF |
| DISEASES | Autoimmune disease of musculoskeletal system | 6 | 0.94 | 0.00083 | CCL2,IFNG,IL12B,IRF7,TNF,IFIH1 |
| DISEASES | Adult respiratory distress syndrome | 3 | 1.19 | 0.00083 | CCL2,CXCL10,TNF |
| DISEASES | Intestinal disease | 6 | 0.91 | 0.001 | CCL2,NFKB1,IFNG,IL12B,CDKN1A,TNF |
| DISEASES | COVID-19 | 3 | 1.03 | 0.002 | IFNG,CXCL10,TNF |
| DISEASES | Influenza | 3 | 1.03 | 0.002 | IFNG,DDX58,TNF |
| DISEASES | Ulcerative colitis | 3 | 1.03 | 0.002 | NFKB1,IFNG,TNF |
| DISEASES | Myositis | 3 | 1.01 | 0.0022 | CCL8,TNF,IFIH1 |
| DISEASES | Respiratory system disease | 6 | 0.76 | 0.0026 | CCL2,NFKB1,IFNG,CXCL10,DDX58,TNF |
| DISEASES | Crohns disease | 3 | 0.91 | 0.0038 | CCL2,IL12B,TNF |
| DISEASES | Lupus nephritis | 2 | 0.92 | 0.0047 | C3,TNFSF13B |
| DISEASES | Autoimmune disease of skin and connective tissue | 4 | 0.76 | 0.0064 | CCL2,IFNG,IL12B,TNF |
| DISEASES | Hypersensitivity reaction disease | 3 | 0.75 | 0.0094 | IFNG,C3,TNF |
| DISEASES | Glomerulonephritis | 3 | 0.75 | 0.0094 | C3,TNFSF13B,TNF |
| DISEASES | Pulmonary tuberculosis | 2 | 0.8 | 0.0094 | IFNG,TNF |
| DISEASES | Viral infectious disease | 4 | 0.61 | 0.0159 | IFNG,CXCL10,DDX58,TNF |
| DISEASES | Alopecia areata | 3 | 0.66 | 0.0159 | CCL2,IFNG,TNF |
| DISEASES | Sarcoidosis | 2 | 0.69 | 0.0175 | IFNG,TNF |
| DISEASES | Acquired immunodeficiency syndrome | 2 | 0.59 | 0.0309 | IFNG,TNF |
| DISEASES | Psoriatic arthritis | 2 | 0.56 | 0.0352 | IL12B,TNF |
| DISEASES | Skin disease | 6 | 0.42 | 0.0362 | CCL2,IFNG,IL12B,C3,TNF,IFIH1 |
| DISEASES | Upper respiratory tract disease | 3 | 0.52 | 0.0362 | NFKB1,IFNG,TNF |
| DISEASES | Meningitis | 2 | 0.55 | 0.0372 | IFNG,TNF |
| DISEASES | Common variable immunodeficiency | 2 | 0.54 | 0.0403 | NFKB1,NFKB2 |
| TISSUES | Hematopoietic system | 23 | 0.86 | 3.45E-09 | NFKBIA,RELB,CCL2,NFKB1,IFNG,TNFSF10,C3,HAVCR2,LILRB1,NFKBIZ,OAS2,SLAMF7,NFKB2,SRC,TNFSF13B,CD83,IRF7,OAS1,IFI35,T |
| TISSUES | Blood | 19 | 0.98 | 1.42E-08 | NFKBIA,RELB,CCL2,NFKB1,IFNG,C3,HAVCR2,LILRB1,NFKBIZ,OAS2,SRC,TNFSF13B,CD83,IRF7,OAS1,IFI35,TNF,IDO1,CCL4 |

| | | | | | |
|---|---|---|---|---|---|
| TISSUES | Immune system | 16 | 0.83 | 2.88E-06 | NFKBIA,RELB,CCL2,NFKB1,IFNG,LILRB1,SLAMF7,NFKB2,TNFSF13B,CD83,IRF7,OAS1,TNF,IDO1,CCL4,IFIH1 |
| TISSUES | Lymphoid tissue | 15 | 0.78 | 1.17E-05 | NFKBIA,RELB,NFKB1,IFNG,LILRB1,SLAMF7,NFKB2,TNFSF13B,CD83,IRF7,OAS1,TNF,IDO1,CCL4,IFIH1 |
| TISSUES | Hematopoietic cell | 12 | 0.85 | 3.55E-05 | NFKBIA,RELB,CCL2,NFKB1,IFNG,LILRB1,NFKBIZ,TNFSF13B,OAS1,TNF,IDO1,CCL4 |
| TISSUES | Peripheral blood | 7 | 1.23 | 4.45E-05 | IFNG,NFKBIZ,TNFSF13B,CD83,IRF7,TNF,IDO1 |
| TISSUES | Mononuclear cell | 6 | 1.36 | 4.45E-05 | NFKBIA,CCL2,IFNG,NFKBIZ,TNFSF13B,TNF |
| TISSUES | Leukocyte | 11 | 0.82 | 8.10E-05 | NFKBIA,RELB,NFKB1,IFNG,LILRB1,NFKBIZ,TNFSF13B,OAS1,TNF,IDO1,CCL4 |
| TISSUES | Monocyte | 5 | 1.25 | 0.00017 | NFKBIA,IFNG,NFKBIZ,TNFSF13B,TNF |
| TISSUES | Monocytic leukemia cell line | 3 | 1.22 | 0.00063 | NFKBIA,IFNG,TNF |
| TISSUES | Inflammatory cell | 3 | 1.22 | 0.00063 | CCL2,IFNG,TNF |
| TISSUES | Regulatory T-lymphocyte | 3 | 1.21 | 0.00065 | IFNG,TNF,IDO1 |
| TISSUES | Lymph node | 6 | 0.91 | 0.00067 | NFKBIA,IFNG,SLAMF7,TNFSF13B,CD83,IRF7 |
| TISSUES | Macrophage cell line | 3 | 1.2 | 0.00067 | NFKBIA,IFNG,TNF |
| TISSUES | Lymphocyte | 8 | 0.62 | 0.0026 | RELB,NFKB1,IFNG,LILRB1,OAS1,TNF,IDO1,CCL4 |
| TISSUES | Macrophage | 4 | 0.88 | 0.0026 | CCL2,IFNG,SLAMF7,TNF |
| TISSUES | Resting cell | 2 | 0.98 | 0.0033 | NFKBIA,RELB |
| TISSUES | Leukemia cell line | 4 | 0.79 | 0.0044 | NFKBIA,IFNG,TNFSF13B,TNF |
| TISSUES | Dendritic cell | 4 | 0.79 | 0.0045 | IFNG,TNFSF13B,CD83,IRF7 |
| TISSUES | B-lymphocyte cell line | 4 | 0.75 | 0.0058 | IFNG,LILRB1,CD83,MX1 |
| TISSUES | Neutrophil | 3 | 0.81 | 0.0061 | IFNG,TNFSF13B,TNF |
| TISSUES | Peritoneal macrophage | 2 | 0.84 | 0.0072 | IFNG,TNF |
| TISSUES | Bronchoalveolar lavage fluid | 2 | 0.8 | 0.0089 | IFNG,TNF |
| TISSUES | T-lymphocyte | 5 | 0.6 | 0.0099 | RELB,IFNG,TNF,IDO1,CCL4 |
| TISSUES | Alveolar macrophage | 2 | 0.77 | 0.0104 | IFNG,TNF |
| TISSUES | RAW-264,7 cell | 2 | 0.77 | 0.0104 | NFKBIA,TNF |
| TISSUES | Bone marrow-derived macrophage | 2 | 0.77 | 0.0104 | IFNG,TNF |
| TISSUES | Whole body | 32 | 0.23 | 0.0113 | NFKBIA,RELB,CCL2,NFKB1,IFNG,TNFSF10,C3,IRAK2,CXCL10,HAVCR2,STAT2,LILRB1,NFKBIZ,OAS2,SLAMF7,NFKB2,IFI44L,SRC,TNF |
| TISSUES | M1 macrophage | 2 | 0.76 | 0.0113 | IFNG,TNF |
| TISSUES | Spleen | 6 | 0.47 | 0.0205 | IFNG,SLAMF7,TNFSF13B,CD83,IRF7,IFIH1 |
| TISSUES | Peritoneum | 2 | 0.61 | 0.0253 | IFNG,TNF |
| TISSUES | Respiratory system | 10 | 0.35 | 0.0289 | C3,HAVCR2,STAT2,NFKBIZ,OAS2,SLAMF7,SRC,CDKN1A,TNF,IDO1 |
| TISSUES | Parenchyma | 2 | 0.57 | 0.0334 | CCL2,TNF |
| TISSUES | Chronic lymphocytic leukemia cell | 4 | 0.47 | 0.036 | RELB,NFKB1,SRC,CD83 |
| TISSUES | Adult | 2 | 0.52 | 0.0422 | IFNG,TNF |
| Subcellular localis | NF-kappaB complex | 7 | 3.48 | 2.22E-10 | NFKBIA,RELB,CCL2,NFKB1,IFNG,NFKB2,TNF |
| Subcellular localis | interleukin-12 complex | 5 | 2.14 | 1.48E-06 | CCL2,IFNG,IL12B,CXCL10,TNF |
| Subcellular localis | I-kappaB/NF-kappaB complex | 3 | 1.54 | 0.00012 | NFKBIA,NFKB1,NFKB2 |
| Subcellular localis | Interferon regulatory factor complex | 3 | 1.12 | 0.0012 | DDX58,IRF7,IFIH1 |
| Subcellular localis | interleukin-23 complex | 3 | 1.06 | 0.0017 | IFNG,IL12B,TNF |
| Subcellular localis | IKKalpha-IKKalpha complex | 2 | 0.8 | 0.0094 | RELB,NFKB2 |
| Subcellular localis | Interferon regulatory factor 3 complex | 2 | 0.62 | 0.0263 | DDX58,IFIH1 |
| Subcellular localis | STING complex | 2 | 0.57 | 0.0356 | DDX58,IFIH1 |
| Subcellular localis | IkappaB kinase complex | 2 | 0.51 | 0.0496 | NFKBIA,RELB |
| Subcellular localis | Extracellular region | 14 | 0.5 | 0.0017 | CCL2,NFKB1,IFNG,IL12B,TNFSF10,C3,CXCL10,SRC,TNFSF13B,CCL8,IFI35,TNF,CXCL2,CCL4 |
| Subcellular localis | Extracellular space | 9 | 0.46 | 0.0134 | CCL2,IFNG,IL12B,TNFSF10,C3,CXCL10,SRC,IFI35,TNF |
| Subcellular localis | Cytosol | 14 | 0.28 | 0.0499 | NFKBIA,RELB,NFKB1,STAT2,NFKB2,SRC,DDX58,IRF7,MX1,CDKN1A,OAS1,IFI35,IDO1,IFIH1 |
| Subcellular localis | Membrane | 20 | 0.25 | 0.0499 | NFKBIA,IFNG,IL12B,TNFSF10,C3,IRAK2,HAVCR2,STAT2,LILRB1,SLAMF7,SRC,TNFSF13B,CD83,DDX58,IRF7,MX1,IFI35,TNF,IFI27,IFIH1 |
| Subcellular localis | Cellular anatomical entity | 33 | 0.21 | 0.0356 | NFKBIA,RELB,CCL2,NFKB1,IFNG,IL12B,TNFSF10,C3,IRAK2,CXCL10,HAVCR2,STAT2,LILRB1,NFKBIZ,OAS2,SLAMF7,NFKB2,SRC,TNF[illegible] |
| UniProt Keywords | Antiviral defense | 10 | 3.08 | 1.43E-11 | IFNG,STAT2,OAS2,IFI44L,DDX58,IRF7,MX1,OAS1,IFI27,IFIH1 |
| UniProt Keywords | Immunity | 14 | 1.92 | 4.16E-11 | C3,HAVCR2,LILRB1,OAS2,SLAMF7,SRC,TNFSF13B,DDX58,IRF7,MX1,OAS1,IDO1,IFI27,IFIH1 |
| UniProt Keywords | Cytokine | 10 | 2.54 | 1.86E-10 | CCL2,IFNG,IL12B,TNFSF10,CXCL10,TNFSF13B,CCL8,TNF,CXCL2,CCL4 |
| UniProt Keywords | Innate immunity | 10 | 1.74 | 3.00E-08 | C3,HAVCR2,OAS2,SLAMF7,DDX58,IRF7,MX1,OAS1,IFI27,IFIH1 |
| UniProt Keywords | Inflammatory response | 7 | 1.65 | 1.85E-06 | CCL2,C3,CXCL10,HAVCR2,CCL8,CXCL2,CCL4 |
| UniProt Keywords | Chemotaxis | 5 | 1.33 | 7.97E-05 | CCL2,CXCL10,CCL8,CXCL2,CCL4 |
| UniProt Keywords | Secreted | 13 | 0.55 | 0.00057 | CCL2,IFNG,IL12B,TNFSF10,C3,CXCL10,LILRB1,TNFSF13B,CCL8,OAS1,TNF,CXCL2,CCL4 |

| | | | | | |
|---|---|---|---|---|---|
| UniProt Keywords | Host-virus interaction | 7 | 0.64 | 0.0029 | NFKBIA,STAT2,SRC,DDX58,IRF7,IFI27,IFIH1 |
| UniProt Keywords | Pyrrolidone carboxylic acid | 3 | 0.63 | 0.0177 | CCL2,IFNG,CCL8 |
| Pfam | Rel homology DNA-binding domain | 3 | 0.97 | 0.0031 | RELB,NFKB1,NFKB2 |
| Pfam | 2-5-oligoadenylate synthetase 1, domain 2, C-terminus | 2 | 0.58 | 0.0332 | OAS2,OAS1 |
| Pfam | C-terminal domain of RIG-I | 2 | 0.58 | 0.0332 | DDX58,IFIH1 |
| Pfam | Caspase recruitment domain | 2 | 0.58 | 0.0332 | DDX58,IFIH1 |
| InterPro | Chemokine interleukin-8-like domain | 5 | 1.4 | 0.00014 | CCL2,CXCL10,CCL8,CXCL2,CCL4 |
| InterPro | Chemokine interleukin-8-like superfamily | 5 | 1.4 | 0.00014 | CCL2,CXCL10,CCL8,CXCL2,CCL4 |
| InterPro | NF-kappa-B/Dorsal | 3 | 1.26 | 0.00062 | RELB,NFKB1,NFKB2 |
| InterPro | Rel homology domain, conserved site | 3 | 1.26 | 0.00062 | RELB,NFKB1,NFKB2 |
| InterPro | NFkappaB IPT domain | 3 | 1.26 | 0.00062 | RELB,NFKB1,NFKB2 |
| InterPro | Death-like domain superfamily | 5 | 1.02 | 0.00099 | NFKB1,IRAK2,NFKB2,DDX58,IFIH1 |
| InterPro | p53-like transcription factor, DNA-binding domain super | 4 | 1.04 | 0.0014 | RELB,NFKB1,STAT2,NFKB2 |
| InterPro | Rel homology domain, DNA-binding domain | 3 | 1.1 | 0.0014 | RELB,NFKB1,NFKB2 |
| InterPro | Rel homology dimerisation domain | 3 | 1.1 | 0.0014 | RELB,NFKB1,NFKB2 |
| InterPro | Rel homology domain (RHD), DNA-binding domain sup | 3 | 1.1 | 0.0014 | RELB,NFKB1,NFKB2 |
| InterPro | Immunoglobulin-like fold | 9 | 0.65 | 0.0021 | RELB,NFKB1,IL12B,C3,HAVCR2,LILRB1,SLAMF7,NFKB2,CD83 |
| InterPro | Tumour necrosis factor domain | 3 | 0.93 | 0.0036 | TNFSF10,TNFSF13B,TNF |
| InterPro | CC chemokine, conserved site | 3 | 0.87 | 0.0051 | CCL2,CCL8,CCL4 |
| InterPro | Chemokine beta/gamma/delta | 3 | 0.78 | 0.0085 | CCL2,CCL8,CCL4 |
| InterPro | IPT domain | 3 | 0.73 | 0.0118 | RELB,NFKB1,NFKB2 |
| InterPro | RIG-I-like receptor, C-terminal regulatory domain | 2 | 0.75 | 0.0129 | DDX58,IFIH1 |
| InterPro | Caspase recruitment domain | 2 | 0.75 | 0.0129 | DDX58,IFIH1 |
| InterPro | RIG-I-like receptor, C-terminal domain superfamily | 2 | 0.75 | 0.0129 | DDX58,IFIH1 |
| InterPro | RIG-I-like receptor, C-terminal | 2 | 0.75 | 0.0129 | DDX58,IFIH1 |
| InterPro | Death domain | 3 | 0.7 | 0.0136 | NFKB1,IRAK2,NFKB2 |
| InterPro | 2-5OAS/ClassI-CCAase, nucleotidyltransferase domain | 2 | 0.72 | 0.0147 | OAS2,OAS1 |
| InterPro | 2-5-oligoadenylate synthetase, C-terminal conserved site | 2 | 0.72 | 0.0147 | OAS2,OAS1 |
| InterPro | 2-5-oligoadenylate synthetase 1, domain 2/C-terminal | 2 | 0.72 | 0.0147 | OAS2,OAS1 |
| InterPro | 2-5-oligoadenylate synthetase, N-terminal conserved site | 2 | 0.72 | 0.0147 | OAS2,OAS1 |
| InterPro | Tumour necrosis factor-like domain superfamily | 3 | 0.54 | 0.035 | TNFSF10,TNFSF13B,TNF |
| InterPro | Polymerase, nucleotidyl transferase domain | 2 | 0.53 | 0.0434 | OAS2,OAS1 |
| SMART | Intercrine alpha family (small cytokine C-X-C) (chemoki | 5 | 1.73 | 1.34E-05 | CCL2,CXCL10,CCL8,CXCL2,CCL4 |
| SMART | Tumour necrosis factor family, | 3 | 0.99 | 0.0025 | TNFSF10,TNFSF13B,TNF |
| SMART | Ig-like, plexins, transcription factors | 3 | 0.86 | 0.005 | RELB,NFKB1,NFKB2 |
