## Supplementary material for "Functional alterations of immune gene expression in ICU and non-ICU patients with Legionnaires’ disease, a prospective observational study": Table S5

| #category | term ID | term description | observed gene count | signal | FDR | matching proteins in your network (labels) |
|---|---|---|---|---|---|---|
| GO Process | GO:0019221 | Cytokine-mediated signaling pathway | 13 | 3.07 | 9.80E-14 | IFNG,IL12B,CXCL10,STAT2,LILRB1,CXCL9,TNFSF13B,IFNB1,CCL8,OAS1,TNF,CCL4,SOCS1 |
| GO Process | GO:0002822 | Regulation of adaptive immune response based on somatic recombinatio | 9 | 2.96 | 2.09E-10 | IL2,IL12B,C3,HAVCR2,LILRB1,NFKBIZ,TNFSF13B,IFNB1,TNF |
| GO Process | GO:0009615 | Response to virus | 12 | 2.86 | 1.43E-12 | IFNG,IL12B,CXCL10,STAT2,LILRB1,CXCL9,IFNB1,CCL8,OAS1,TNF,CCL4,IFIH1 |
| GO Process | GO:0045088 | Regulation of innate immune response | 9 | 2.51 | 1.78E-09 | IL12B,C3,HAVCR2,STAT2,LILRB1,IFNB1,OAS1,IFI35,SOCS1 |
| GO Process | GO:0051607 | Defense response to virus | 9 | 2.43 | 2.70E-09 | IFNG,IL12B,CXCL10,STAT2,LILRB1,CXCL9,IFNB1,OAS1,IFIH1 |
| GO Process | GO:0050729 | Positive regulation of inflammatory response | 7 | 2.36 | 7.23E-08 | IL2,IFNG,IL12B,C3,NFKBIZ,IFI35,TNF |
| GO Process | GO:0043370 | Regulation of CD4-positive, alpha-beta T cell differentiation | 5 | 2.32 | 1.05E-06 | IL2,IFNG,IL12B,NFKBIZ,SOCS1 |
| GO Process | GO:1903039 | Positive regulation of leukocyte cell-cell adhesion | 9 | 2.31 | 5.02E-09 | IL2,IFNG,IL12B,HAVCR2,LILRB1,NFKBIZ,TNFSF13B,TNF,SOCS1 |
| GO Process | GO:0002250 | Adaptive immune response | 10 | 2.23 | 1.61E-09 | IL2,IFNG,IL12B,C3,HAVCR2,LILRB1,SLAMF7,NFKB2,TNFSF13B,IFNB1 |
| GO Process | GO:1903037 | Regulation of leukocyte cell-cell adhesion | 10 | 2.2 | 1.89E-09 | IL2,IFNG,IL12B,HAVCR2,LILRB1,NFKBIZ,TNFSF13B,IFNB1,TNF,SOCS1 |
| GO Process | GO:0031349 | Positive regulation of defense response | 9 | 2.19 | 9.20E-09 | IL2,IFNG,IL12B,C3,HAVCR2,NFKBIZ,IFNB1,IFI35,TNF |
| GO Process | GO:0002824 | Positive regulation of adaptive immune response based on somatic reco | 6 | 2.19 | 5.71E-07 | IL2,IL12B,C3,NFKBIZ,TNFSF13B,TNF |
| GO Process | GO:0032103 | Positive regulation of response to external stimulus | 11 | 1.34 | 4.54E-10 | IL2,IFNG,IL12B,C3,CXCL10,HAVCR2,NFKBIZ,IFNB1,IFI35,TNF,CCL4 |
| GO Process | GO:0046634 | Regulation of alpha-beta T cell activation | 6 | 2.15 | 7.16E-07 | IL2,IFNG,IL12B,LILRB1,NFKBIZ,SOCS1 |
| GO Process | GO:0002706 | Regulation of lymphocyte mediated immunity | 7 | 2.12 | 2.24E-07 | IL2,IL12B,C3,HAVCR2,LILRB1,IFNB1,TNF |
| GO Process | GO:0031347 | Regulation of defense response | 13 | 2.1 | 1.94E-11 | IL2,IFNG,IL12B,C3,HAVCR2,STAT2,LILRB1,NFKBIZ,IFNB1,OAS1,IFI35,TNF,SOCS1 |
| GO Process | GO:0050870 | Positive regulation of T cell activation | 8 | 2.07 | 7.23E-08 | IL2,IFNG,IL12B,HAVCR2,LILRB1,NFKBIZ,TNFSF13B,SOCS1 |
| GO Process | GO:0050777 | Negative regulation of immune response | 7 | 2.06 | 2.98E-07 | IL2,IL12B,HAVCR2,STAT2,LILRB1,IFNB1,OAS1 |
| GO Process | GO:0006955 | Immune response | 20 | 2.04 | 8.86E-18 | IL2,IFNG,IL12B,TNFSF10,C3,CXCL10,HAVCR2,STAT2,LILRB1,CXCL9,SLAMF7,NFKB2,TNFSF13B,IFNB1,CCL8,OAS1,IFI35,TNF,CCL4,IFIH1 |
| GO Process | GO:0045087 | Innate immune response | 14 | 2.04 | 5.61E-12 | IFNG,IL12B,C3,CXCL10,HAVCR2,STAT2,SLAMF7,IFNB1,CCL8,OAS1,IFI35,TNF,CCL4,IFIH1 |
| GO Process | GO:0002366 | Leukocyte activation involved in immune response | 7 | 2.04 | 3.39E-07 | IL2,IFNG,IL12B,HAVCR2,LILRB1,IFNB1,IFI35 |
| GO Process | GO:0050776 | Regulation of immune response | 15 | 2.03 | 1.15E-12 | IL2,IFNG,IL12B,C3,HAVCR2,STAT2,LILRB1,NFKBIZ,TNFSF13B,IFNB1,OAS1,IFI35,TNF,SOCS1,IFIH1 |
| GO Process | GO:0043372 | Positive regulation of CD4-positive, alpha-beta T cell differentiation | 4 | 2.02 | 1.19E-05 | IFNG,IL12B,NFKBIZ,SOCS1 |
| GO Process | GO:0050778 | Positive regulation of immune response | 11 | 2 | 1.29E-09 | IL2,IFNG,IL12B,C3,HAVCR2,LILRB1,NFKBIZ,TNFSF13B,IFNB1,IFI35,TNF |
| GO Process | GO:0042509 | Regulation of tyrosine phosphorylation of STAT protein | 5 | 1.99 | 4.93E-06 | IL2,IFNG,IL12B,TNF,SOCS1 |
| GO Process | GO:0002684 | Positive regulation of immune system process | 15 | 1.98 | 1.43E-12 | IL2,IFNG,IL12B,C3,CXCL10,HAVCR2,LILRB1,NFKBIZ,TNFSF13B,IFNB1,CCL8,IFI35,TNF,CCL4,SOCS1 |
| GO Process | GO:0032729 | Positive regulation of interferon-gamma production | 5 | 1.98 | 5.13E-06 | IL2,IL12B,HAVCR2,LILRB1,TNF |
| GO Process | GO:0098542 | Defense response to other organism | 16 | 1.94 | 3.23E-13 | IFNG,IL12B,C3,CXCL10,HAVCR2,STAT2,LILRB1,CXCL9,SLAMF7,IFNB1,CCL8,OAS1,IFI35,TNF,CCL4,IFIH1 |
| GO Process | GO:0034138 | Toll-like receptor 3 signaling pathway | 3 | 1.94 | 3.59E-05 | HAVCR2,OAS1,TNF |
| GO Process | GO:0098586 | Cellular response to virus | 5 | 1.89 | 8.00E-06 | IFNG,CXCL10,IFNB1,OAS1,IFIH1 |
| GO Process | GO:0002252 | Immune effector process | 9 | 1.88 | 5.44E-08 | IL2,IFNG,IL12B,C3,HAVCR2,LILRB1,SLAMF7,IFNB1,IFI35 |
| GO Process | GO:0050863 | Regulation of T cell activation | 9 | 1.87 | 5.44E-08 | IL2,IFNG,IL12B,HAVCR2,LILRB1,NFKBIZ,TNFSF13B,IFNB1,SOCS1 |
| GO Process | GO:0002696 | Positive regulation of leukocyte activation | 9 | 1.85 | 6.24E-08 | IL2,IFNG,IL12B,HAVCR2,LILRB1,NFKBIZ,TNFSF13B,TNF,SOCS1 |
| GO Process | GO:0002697 | Regulation of immune effector process | 9 | 1.85 | 6.24E-08 | IL2,IFNG,IL12B,C3,HAVCR2,LILRB1,NFKBIZ,IFNB1,TNF |
| GO Process | GO:0032101 | Regulation of response to external stimulus | 15 | 1.83 | 4.76E-12 | IL2,IFNG,IL12B,C3,CXCL10,HAVCR2,STAT2,LILRB1,NFKBIZ,IFNB1,OAS1,IFI35,TNF,CCL4,SOCS1 |
| GO Process | GO:1902105 | Regulation of leukocyte differentiation | 8 | 1.8 | 3.25E-07 | IL2,IFNG,IL12B,LILRB1,NFKBIZ,IFNB1,TNF,SOCS1 |
| GO Process | GO:0001775 | Cell activation | 12 | 1.77 | 1.29E-09 | IL2,IFNG,IL12B,CXCL10,HAVCR2,LILRB1,SLAMF7,NFKB2,TNFSF13B,IFNB1,IFI35,TNF |
| GO Process | GO:0032680 | Regulation of tumor necrosis factor production | 6 | 1.77 | 4.56E-06 | IFNG,IL12B,HAVCR2,LILRB1,OAS1,IFIH1 |
| GO Process | GO:0032760 | Positive regulation of tumor necrosis factor production | 5 | 1.75 | 1.54E-05 | IFNG,IL12B,HAVCR2,OAS1,IFIH1 |
| GO Process | GO:0031341 | Regulation of cell killing | 5 | 1.74 | 1.59E-05 | IFNG,IL12B,C3,HAVCR2,LILRB1 |
| GO Process | GO:0071346 | Cellular response to interferon-gamma | 5 | 1.72 | 1.80E-05 | IFNG,IL12B,CCL8,TNF,CCL4 |
| GO Process | GO:0007259 | Receptor signaling pathway via JAK-STAT | 4 | 1.71 | 4.79E-05 | IL2,IFNG,STAT2,SOCS1 |
| GO Process | GO:0002683 | Negative regulation of immune system process | 9 | 1.7 | 1.44E-07 | IL2,IL12B,HAVCR2,STAT2,LILRB1,IFNB1,OAS1,TNF,SOCS1 |
| GO Process | GO:0045580 | Regulation of T cell differentiation | 6 | 1.69 | 6.83E-06 | IL2,IFNG,IL12B,NFKBIZ,IFNB1,SOCS1 |
| GO Process | GO:1902107 | Positive regulation of leukocyte differentiation | 6 | 1.67 | 7.36E-06 | IL2,IFNG,IL12B,NFKBIZ,TNF,SOCS1 |
| GO Process | GO:0002699 | Positive regulation of immune effector process | 7 | 1.66 | 2.40E-06 | IL2,IFNG,IL12B,C3,LILRB1,NFKBIZ,TNF |
| GO Process | GO:0002682 | Regulation of immune system process | 18 | 1.65 | 9.80E-14 | IL2,IFNG,IL12B,C3,CXCL10,HAVCR2,STAT2,LILRB1,NFKBIZ,TNFSF13B,IFNB1,CCL8,OAS1,IFI35,TNF,CCL4,SOCS1,IFIH1 |
| GO Process | GO:0006959 | Humoral immune response | 7 | 1.65 | 2.62E-06 | IFNG,C3,CXCL10,CXCL9,IFNB1,CCL8,TNF |
| GO Process | GO:0045071 | Negative regulation of viral genome replication | 4 | 1.65 | 6.56E-05 | IFNB1,OAS1,TNF,IFIH1 |
| GO Process | GO:0045582 | Positive regulation of T cell differentiation | 5 | 1.63 | 2.71E-05 | IL2,IFNG,IL12B,NFKBIZ,SOCS1 |
| GO Process | GO:0002823 | Negative regulation of adaptive immune response based on somatic reco | 4 | 1.63 | 6.96E-05 | IL2,HAVCR2,LILRB1,IFNB1 |
| GO Process | GO:0045321 | Leukocyte activation | 10 | 1.6 | 7.23E-08 | IL2,IFNG,IL12B,HAVCR2,LILRB1,SLAMF7,TNFSF13B,IFNB1,IFI35,TNF |
| GO Process | GO:0071222 | Cellular response to lipopolysaccharide | 6 | 1.6 | 1.08E-05 | IL12B,CXCL10,HAVCR2,LILRB1,CXCL9,TNF |
| GO Process | GO:0001818 | Negative regulation of cytokine production | 7 | 1.59 | 3.69E-06 | IFNG,IL12B,HAVCR2,LILRB1,IFNB1,OAS1,TNF |
| GO Process | GO:0006952 | Defense response | 17 | 1.58 | 1.29E-12 | IFNG,IL12B,C3,CXCL10,HAVCR2,STAT2,LILRB1,NFKBIZ,CXCL9,SLAMF7,IFNB1,CCL8,OAS1,IFI35,TNF,CCL4,IFIH1 |
| GO Process | GO:0001819 | Positive regulation of cytokine production | 9 | 1.58 | 3.28E-07 | IL2,IFNG,IL12B,C3,HAVCR2,LILRB1,OAS1,TNF,IFIH1 |
| GO Process | GO:0002698 | Negative regulation of immune effector process | 5 | 1.58 | 3.53E-05 | IL2,HAVCR2,LILRB1,IFNB1,TNF |
| GO Process | GO:0002281 | Macrophage activation involved in immune response | 3 | 1.58 | 0.00019 | IFNG,HAVCR2,IFI35 |
| GO Process | GO:0046651 | Lymphocyte proliferation | 5 | 1.57 | 3.61E-05 | IL2,IL12B,LILRB1,TNFSF13B,IFNB1 |
| GO Process | GO:0042116 | Macrophage activation | 4 | 1.57 | 9.29E-05 | IFNG,HAVCR2,IFI35,TNF |
| GO Process | GO:2000319 | Regulation of T-helper 17 cell differentiation | 3 | 1.56 | 0.00021 | IL2,IL12B,NFKBIZ |
| GO Process | GO:0002694 | Regulation of leukocyte activation | 10 | 1.55 | 1.04E-07 | IL2,IFNG,IL12B,HAVCR2,LILRB1,NFKBIZ,TNFSF13B,IFNB1,TNF,SOCS1 |
| GO Process | GO:0002521 | Leukocyte differentiation | 8 | 1.54 | 1.45E-06 | IL2,IFNG,IL12B,LILRB1,TNFSF13B,IFNB1,TNF,SOCS1 |

| | | | | | | |
|---|---|---|---|---|---|---|
| GO Process | GO:0001906 | Cell killing | 5 | 1.54 | 4.25E-05 | C3,CXCL10,CXCL9,SLAMF7,CCL8 |
| GO Process | GO:0050868 | Negative regulation of T cell activation | 5 | 1.54 | 4.25E-05 | IL2,HAVCR2,LILRB1,IFNB1,SOCS1 |
| GO Process | GO:0030101 | Natural killer cell activation | 4 | 1.54 | 0.00011 | IL2,IL12B,SLAMF7,IFNB1 |
| GO Process | GO:0042531 | Positive regulation of tyrosine phosphorylation of STAT protein | 4 | 1.54 | 0.00011 | IL2,IFNG,IL12B,TNF |
| GO Process | GO:0002376 | Immune system process | 22 | 1.52 | 8.86E-18 | IL2,IFNG,IL12B,TNFSF10,C3,CXCL10,HAVCR2,STAT2,LILRB1,NFKBIZ,CXCL9,SLAMF7,NFKB2,TNFSF13B,IFNB1,CCL8,OAS1,IFI35,TNF,CCL4,SOCS1,IFIH1 |
| GO Process | GO:1901224 | Positive regulation of NIK/NF-kappaB signaling | 4 | 1.52 | 0.00012 | IL12B,HAVCR2,IFI35,TNF |
| GO Process | GO:0001817 | Regulation of cytokine production | 11 | 1.5 | 4.22E-08 | IL2,IFNG,IL12B,C3,HAVCR2,LILRB1,IFNB1,OAS1,TNF,SOCS1,IFIH1 |
| GO Process | GO:0032722 | Positive regulation of chemokine production | 4 | 1.49 | 0.00014 | IFNG,HAVCR2,OAS1,TNF |
| GO Process | GO:0045670 | Regulation of osteoclast differentiation | 4 | 1.49 | 0.00014 | IFNG,IL12B,LILRB1,TNF |
| GO Process | GO:0045591 | Positive regulation of regulatory T cell differentiation | 3 | 1.49 | 0.00029 | IL2,IFNG,SOCS1 |
| GO Process | GO:0042098 | T cell proliferation | 4 | 1.47 | 0.00015 | IL2,IL12B,LILRB1,TNFSF13B |
| GO Process | GO:0045824 | Negative regulation of innate immune response | 4 | 1.46 | 0.00016 | HAVCR2,STAT2,LILRB1,OAS1 |
| GO Process | GO:0006954 | Inflammatory response | 9 | 1.45 | 7.16E-07 | IFNG,C3,CXCL10,HAVCR2,NFKBIZ,CXCL9,CCL8,TNF,CCL4 |
| GO Process | GO:0031348 | Negative regulation of defense response | 6 | 1.44 | 2.63E-05 | IL2,IL12B,HAVCR2,STAT2,LILRB1,OAS1 |
| GO Process | GO:0060557 | Positive regulation of vitamin D biosynthetic process | 2 | 1.44 | 0.00059 | IFNG,TNF |
| GO Process | GO:0045672 | Positive regulation of osteoclast differentiation | 3 | 1.43 | 0.00039 | IFNG,IL12B,TNF |
| GO Process | GO:0030593 | Neutrophil chemotaxis | 4 | 1.4 | 0.00021 | CXCL10,CXCL9,CCL8,CCL4 |
| GO Process | GO:0070098 | Chemokine-mediated signaling pathway | 4 | 1.39 | 0.00023 | CXCL10,CXCL9,CCL8,CCL4 |
| GO Process | GO:0060559 | Positive regulation of calcidiol 1-monooxygenase activity | 2 | 1.35 | 0.00092 | IFNG,TNF |
| GO Process | GO:2001189 | Negative regulation of T cell activation via T cell receptor contact with a | 2 | 1.35 | 0.00092 | HAVCR2,LILRB1 |
| GO Process | GO:0002520 | Immune system development | 10 | 1.3 | 6.40E-07 | IL2,IFNG,IL12B,HAVCR2,LILRB1,NFKB2,TNFSF13B,IFNB1,TNF,SOCS1 |
| GO Process | GO:0070663 | Regulation of leukocyte proliferation | 6 | 1.3 | 5.51E-05 | IL2,IL12B,HAVCR2,LILRB1,TNFSF13B,CCL8 |
| GO Process | GO:0002719 | Negative regulation of cytokine production involved in immune respons | 3 | 1.29 | 0.00078 | LILRB1,IFNB1,TNF |
| GO Process | GO:0032102 | Negative regulation of response to external stimulus | 7 | 1.28 | 2.32E-05 | IL2,IL12B,HAVCR2,STAT2,LILRB1,OAS1,TNF |
| GO Process | GO:0042129 | Regulation of T cell proliferation | 5 | 1.25 | 0.00019 | IL2,IL12B,HAVCR2,LILRB1,TNFSF13B |
| GO Process | GO:0070664 | Negative regulation of leukocyte proliferation | 4 | 1.25 | 0.00044 | IL2,HAVCR2,LILRB1,CCL8 |
| GO Process | GO:0002221 | Pattern recognition receptor signaling pathway | 4 | 1.25 | 0.00045 | HAVCR2,OAS1,TNF,IFIH1 |
| GO Process | GO:0032479 | Regulation of type I interferon production | 4 | 1.24 | 0.00048 | HAVCR2,LILRB1,OAS1,IFIH1 |
| GO Process | GO:0042110 | T cell activation | 6 | 1.22 | 9.06E-05 | IL2,IL12B,LILRB1,SLAMF7,TNFSF13B,IFNB1 |
| GO Process | GO:1902106 | Negative regulation of leukocyte differentiation | 4 | 1.21 | 0.00055 | IL2,LILRB1,IFNB1,SOCS1 |
| GO Process | GO:0042102 | Positive regulation of T cell proliferation | 4 | 1.2 | 0.00056 | IL2,IL12B,HAVCR2,TNFSF13B |
| GO Process | GO:0002861 | Regulation of inflammatory response to antigenic stimulus | 3 | 1.2 | 0.0012 | IL12B,C3,TNF |
| GO Process | GO:0002891 | Positive regulation of immunoglobulin mediated immune response | 3 | 1.2 | 0.0012 | IL2,C3,TNF |
| GO Process | GO:0032660 | Regulation of interleukin-17 production | 3 | 1.2 | 0.0012 | IL2,IFNG,IL12B |
| GO Process | GO:0060337 | Type I interferon signaling pathway | 3 | 1.2 | 0.0012 | STAT2,IFNB1,OAS1 |
| GO Process | GO:0048534 | Hematopoietic or lymphoid organ development | 9 | 1.18 | 5.71E-06 | IL2,IFNG,IL12B,LILRB1,NFKB2,TNFSF13B,IFNB1,TNF,SOCS1 |
| GO Process | GO:0002764 | Immune response-regulating signaling pathway | 6 | 1.18 | 0.00011 | HAVCR2,LILRB1,NFKBIZ,OAS1,TNF,IFIH1 |
| GO Process | GO:2001186 | Negative regulation of CD8-positive, alpha-beta T cell activation | 2 | 1.18 | 0.0021 | LILRB1,SOCS1 |
| GO Process | GO:0002708 | Positive regulation of lymphocyte mediated immunity | 4 | 1.16 | 0.00071 | IL2,IL12B,C3,TNF |
| GO Process | GO:0046636 | Negative regulation of alpha-beta T cell activation | 3 | 1.15 | 0.0015 | IL2,LILRB1,SOCS1 |
| GO Process | GO:0042269 | Regulation of natural killer cell mediated cytotoxicity | 3 | 1.14 | 0.0016 | IL12B,HAVCR2,LILRB1 |
| GO Process | GO:0045581 | Negative regulation of T cell differentiation | 3 | 1.14 | 0.0016 | IL2,IFNB1,SOCS1 |
| GO Process | GO:2000321 | Positive regulation of T-helper 17 cell differentiation | 2 | 1.14 | 0.0026 | IL12B,NFKBIZ |
| GO Process | GO:0043457 | Regulation of cellular respiration | 3 | 1.13 | 0.0016 | IFNG,OAS1,TNF |
| GO Process | GO:0048247 | Lymphocyte chemotaxis | 3 | 1.1 | 0.0019 | CXCL10,CCL8,CCL4 |
| GO Process | GO:0002858 | Regulation of natural killer cell mediated cytotoxicity directed against tu | 2 | 1.1 | 0.0031 | IL12B,HAVCR2 |
| GO Process | GO:0010533 | Regulation of activation of Janus kinase activity | 2 | 1.07 | 0.0037 | IL12B,SOCS1 |
| GO Process | GO:0051709 | Regulation of killing of cells of another organism | 2 | 1.07 | 0.0037 | IFNG,C3 |
| GO Process | GO:1900222 | Negative regulation of amyloid-beta clearance | 2 | 1.07 | 0.0037 | IFNG,TNF |
| GO Process | GO:0043065 | Positive regulation of apoptotic process | 7 | 1.05 | 0.00011 | IFNG,IL12B,TNFSF10,C3,LILRB1,IFNB1,TNF |
| GO Process | GO:0002707 | Negative regulation of lymphocyte mediated immunity | 3 | 1.05 | 0.0024 | HAVCR2,LILRB1,IFNB1 |
| GO Process | GO:0007166 | Cell surface receptor signaling pathway | 16 | 1.04 | 4.03E-09 | IL2,IFNG,IL12B,C3,CXCL10,STAT2,LILRB1,NFKBIZ,CXCL9,TNFSF13B,IFNB1,CCL8,OAS1,TNF,CCL4,SOCS1 |
| GO Process | GO:0009617 | Response to bacterium | 8 | 1.04 | 4.57E-05 | IL12B,C3,CXCL10,HAVCR2,LILRB1,CXCL9,OAS1,TNF |
| GO Process | GO:0050900 | Leukocyte migration | 5 | 1.04 | 0.00062 | CXCL10,CXCL9,CCL8,TNF,CCL4 |
| GO Process | GO:0032648 | Regulation of interferon-beta production | 3 | 1.04 | 0.0026 | LILRB1,OAS1,IFIH1 |
| GO Process | GO:0002467 | Germinal center formation | 2 | 1.04 | 0.0043 | NFKB2,TNFSF13B |
| GO Process | GO:0045089 | Positive regulation of innate immune response | 4 | 1.03 | 0.0014 | IL12B,HAVCR2,IFNB1,IFI35 |
| GO Process | GO:0034105 | Positive regulation of tissue remodeling | 2 | 1.01 | 0.0049 | IL2,IL12B |
| GO Process | GO:0048584 | Positive regulation of response to stimulus | 16 | 1 | 7.35E-09 | IL2,IFNG,IL12B,TNFSF10,C3,CXCL10,HAVCR2,LILRB1,NFKBIZ,TNFSF13B,IFNB1,CCL8,IFI35,TNF,CCL4,IFIH1 |
| GO Process | GO:0002687 | Positive regulation of leukocyte migration | 4 | 1 | 0.0016 | CXCL10,CCL8,TNF,CCL4 |
| GO Process | GO:0032675 | Regulation of interleukin-6 production | 4 | 1 | 0.0016 | IFNG,HAVCR2,TNF,IFIH1 |
| GO Process | GO:0051241 | Negative regulation of multicellular organismal process | 10 | 0.99 | 9.75E-06 | IL2,IFNG,IL12B,CXCL10,HAVCR2,LILRB1,IFNB1,OAS1,TNF,SOCS1 |
| GO Process | GO:0032655 | Regulation of interleukin-12 production | 3 | 0.99 | 0.0032 | IFNG,IL12B,LILRB1 |
| GO Process | GO:0008285 | Negative regulation of cell population proliferation | 8 | 0.98 | 7.34E-05 | IL2,IFNG,IL12B,HAVCR2,LILRB1,CCL8,IFI35,TNF |
| GO Process | GO:0034393 | Positive regulation of smooth muscle cell apoptotic process | 2 | 0.98 | 0.0056 | IFNG,IL12B |
| GO Process | GO:0071310 | Cellular response to organic substance | 15 | 0.96 | 4.89E-08 | IFNG,IL12B,CXCL10,HAVCR2,STAT2,LILRB1,CXCL9,TNFSF13B,IFNB1,CCL8,OAS1,TNF,CCL4,SOCS1,IFIH1 |
| GO Process | GO:0002863 | Positive regulation of inflammatory response to antigenic stimulus | 2 | 0.96 | 0.0063 | C3,TNF |

| | | | | | | |
|---|---|---|---|---|---|---|
| GO Process | GO:0060759 | Regulation of response to cytokine stimulus | 4 | 0.95 | 0.0021 | STAT2,OAS1,SOCS1,IFIH1 |
| GO Process | GO:0031640 | Killing of cells of another organism | 3 | 0.94 | 0.0042 | CXCL10,CXCL9,CCL8 |
| GO Process | GO:0002460 | Adaptive immune response based on somatic recombination of immune | 4 | 0.93 | 0.0023 | IL12B,C3,NFKB2,TNFSF13B |
| GO Process | GO:0032732 | Positive regulation of interleukin-1 production | 3 | 0.93 | 0.0044 | IFNG,HAVCR2,TNF |
| GO Process | GO:1901857 | Positive regulation of cellular respiration | 2 | 0.93 | 0.0071 | IFNG,OAS1 |
| GO Process | GO:0050766 | Positive regulation of phagocytosis | 3 | 0.92 | 0.0047 | IFNG,C3,TNF |
| GO Process | GO:0071677 | Positive regulation of mononuclear cell migration | 3 | 0.92 | 0.0047 | CXCL10,TNF,CCL4 |
| GO Process | GO:0071356 | Cellular response to tumor necrosis factor | 4 | 0.91 | 0.0026 | TNFSF13B,CCL8,TNF,CCL4 |
| GO Process | GO:0002286 | T cell activation involved in immune response | 3 | 0.91 | 0.0048 | IL12B,LILRB1,IFNB1 |
| GO Process | GO:0045661 | Regulation of myoblast differentiation | 3 | 0.91 | 0.005 | CXCL10,CXCL9,TNF |
| GO Process | GO:0032700 | Negative regulation of interleukin-17 production | 2 | 0.91 | 0.0079 | IFNG,IL12B |
| GO Process | GO:0046425 | Regulation of receptor signaling pathway via JAK-STAT | 3 | 0.9 | 0.0051 | IL12B,TNF,SOCS1 |
| GO Process | GO:0010033 | Response to organic substance | 17 | 0.87 | 1.33E-08 | IL2,IFNG,IL12B,TNFSF10,CXCL10,HAVCR2,STAT2,LILRB1,CXCL9,TNFSF13B,IFNB1,CCL8,OAS1,TNF,CCL4,SOCS1,IFIH1 |
| GO Process | GO:1903131 | Mononuclear cell differentiation | 5 | 0.87 | 0.0017 | IL2,IL12B,LILRB1,TNFSF13B,IFNB1 |
| GO Process | GO:0002700 | Regulation of production of molecular mediator of immune response | 4 | 0.87 | 0.0033 | IL2,LILRB1,IFNB1,TNF |
| GO Process | GO:0048143 | Astrocyte activation | 2 | 0.87 | 0.0097 | IFNG,TNF |
| GO Process | GO:0048245 | Eosinophil chemotaxis | 2 | 0.87 | 0.0097 | CCL8,CCL4 |
| GO Process | GO:0051044 | Positive regulation of membrane protein ectodomain proteolysis | 2 | 0.87 | 0.0097 | IFNG,TNF |
| GO Process | GO:0002709 | Regulation of T cell mediated immunity | 3 | 0.85 | 0.0065 | IL12B,LILRB1,IFNB1 |
| GO Process | GO:2000341 | Regulation of chemokine (C-X-C motif) ligand 2 production | 2 | 0.85 | 0.0106 | OAS1,TNF |
| GO Process | GO:0042127 | Regulation of cell population proliferation | 12 | 0.84 | 6.87E-06 | IL2,IFNG,IL12B,CXCL10,HAVCR2,STAT2,LILRB1,CXCL9,TNFSF13B,CCL8,IFI35,TNF |
| GO Process | GO:0045953 | Negative regulation of natural killer cell mediated cytotoxicity | 2 | 0.84 | 0.0115 | HAVCR2,LILRB1 |
| GO Process | GO:0060339 | Negative regulation of type I interferon-mediated signaling pathway | 2 | 0.84 | 0.0115 | STAT2,OAS1 |
| GO Process | GO:0071360 | Cellular response to exogenous dsRNA | 2 | 0.84 | 0.0115 | IFNB1,IFIH1 |
| GO Process | GO:0050672 | Negative regulation of lymphocyte proliferation | 3 | 0.83 | 0.0075 | IL2,HAVCR2,LILRB1 |
| GO Process | GO:0045596 | Negative regulation of cell differentiation | 7 | 0.82 | 0.00057 | IL2,IFNG,CXCL10,LILRB1,IFNB1,TNF,SOCS1 |
| GO Process | GO:0051240 | Positive regulation of multicellular organismal process | 11 | 0.81 | 2.45E-05 | IL2,IFNG,IL12B,C3,HAVCR2,LILRB1,NFKBIZ,OAS1,TNF,SOCS1,IFIH1 |
| GO Process | GO:0032693 | Negative regulation of interleukin-10 production | 2 | 0.81 | 0.0132 | IL12B,LILRB1 |
| GO Process | GO:0033141 | Positive regulation of peptidyl-serine phosphorylation of STAT protein | 2 | 0.81 | 0.0132 | IFNG,IFNB1 |
| GO Process | GO:1901739 | Regulation of myoblast fusion | 2 | 0.81 | 0.0132 | CXCL10,CXCL9 |
| GO Process | GO:1902004 | Positive regulation of amyloid-beta formation | 2 | 0.81 | 0.0132 | IFNG,TNF |
| GO Process | GO:0070374 | Positive regulation of ERK1 and ERK2 cascade | 4 | 0.8 | 0.0048 | HAVCR2,CCL8,TNF,CCL4 |
| GO Process | GO:0002922 | Positive regulation of humoral immune response | 2 | 0.8 | 0.014 | C3,TNF |
| GO Process | GO:0140374 | Antiviral innate immune response | 2 | 0.8 | 0.014 | CXCL10,OAS1 |
| GO Process | GO:0032755 | Positive regulation of interleukin-6 production | 3 | 0.79 | 0.0091 | IFNG,TNF,IFIH1 |
| GO Process | GO:0035458 | Cellular response to interferon-beta | 2 | 0.78 | 0.015 | IFNB1,OAS1 |
| GO Process | GO:0045822 | Negative regulation of heart contraction | 2 | 0.78 | 0.015 | IL2,TNF |
| GO Process | GO:2000026 | Regulation of multicellular organismal development | 10 | 0.76 | 0.00011 | IL2,IFNG,IL12B,C3,CXCL10,LILRB1,NFKBIZ,IFNB1,TNF,SOCS1 |
| GO Process | GO:0034764 | Positive regulation of transmembrane transport | 4 | 0.75 | 0.0066 | IFNG,C3,CXCL10,CXCL9 |
| GO Process | GO:0002323 | Natural killer cell activation involved in immune response | 2 | 0.75 | 0.0178 | IL12B,IFNB1 |
| GO Process | GO:0046639 | Negative regulation of alpha-beta T cell differentiation | 2 | 0.75 | 0.0178 | IL2,SOCS1 |
| GO Process | GO:0033138 | Positive regulation of peptidyl-serine phosphorylation | 3 | 0.74 | 0.0119 | IFNG,IFNB1,TNF |
| GO Process | GO:0010959 | Regulation of metal ion transport | 5 | 0.73 | 0.0042 | IFNG,CXCL10,LILRB1,CXCL9,CCL4 |
| GO Process | GO:0061844 | Antimicrobial humoral immune response mediated by antimicrobial pep | 3 | 0.73 | 0.0128 | CXCL10,CXCL9,CCL8 |
| GO Process | GO:0097191 | Extrinsic apoptotic signaling pathway | 3 | 0.73 | 0.0128 | IL2,IFNG,TNF |
| GO Process | GO:0002675 | Positive regulation of acute inflammatory response | 2 | 0.73 | 0.0189 | C3,TNF |
| GO Process | GO:0002710 | Negative regulation of T cell mediated immunity | 2 | 0.73 | 0.0189 | LILRB1,IFNB1 |
| GO Process | GO:1901623 | Regulation of lymphocyte chemotaxis | 2 | 0.73 | 0.0189 | CXCL10,CCL4 |
| GO Process | GO:0032740 | Positive regulation of interleukin-17 production | 2 | 0.72 | 0.0201 | IL2,IL12B |
| GO Process | GO:0042104 | Positive regulation of activated T cell proliferation | 2 | 0.72 | 0.0201 | IL2,IL12B |
| GO Process | GO:0045662 | Negative regulation of myoblast differentiation | 2 | 0.72 | 0.0201 | CXCL10,TNF |
| GO Process | GO:0048583 | Regulation of response to stimulus | 19 | 0.71 | 1.86E-08 | IL2,IFNG,IL12B,TNFSF10,C3,CXCL10,HAVCR2,STAT2,LILRB1,NFKBIZ,TNFSF13B,IFNB1,CCL8,OAS1,IFI35,TNF,CCL4,SOCS1,IFIH1 |
| GO Process | GO:0030225 | Macrophage differentiation | 2 | 0.71 | 0.0212 | IFNG,SOCS1 |
| GO Process | GO:1901700 | Response to oxygen-containing compound | 10 | 0.69 | 0.00024 | IL2,IL12B,TNFSF10,CXCL10,HAVCR2,STAT2,LILRB1,CXCL9,TNF,SOCS1 |
| GO Process | GO:0001774 | Microglial cell activation | 2 | 0.69 | 0.0237 | IFNG,TNF |
| GO Process | GO:0002825 | Regulation of T-helper 1 type immune response | 2 | 0.69 | 0.0237 | IL12B,HAVCR2 |
| GO Process | GO:0032647 | Regulation of interferon-alpha production | 2 | 0.69 | 0.0237 | HAVCR2,IFIH1 |
| GO Process | GO:0051924 | Regulation of calcium ion transport | 4 | 0.68 | 0.0099 | CXCL10,LILRB1,CXCL9,CCL4 |
| GO Process | GO:0032480 | Negative regulation of type I interferon production | 2 | 0.68 | 0.0248 | HAVCR2,LILRB1 |
| GO Process | GO:0045595 | Regulation of cell differentiation | 10 | 0.67 | 0.00028 | IL2,IFNG,IL12B,CXCL10,LILRB1,NFKBIZ,CXCL9,IFNB1,TNF,SOCS1 |
| GO Process | GO:0051094 | Positive regulation of developmental process | 9 | 0.67 | 0.00056 | IL2,IFNG,IL12B,C3,NFKBIZ,CXCL9,TNFSF13B,TNF,SOCS1 |
| GO Process | GO:2001235 | Positive regulation of apoptotic signaling pathway | 3 | 0.67 | 0.0172 | TNFSF10,IFNB1,TNF |
| GO Process | GO:0048585 | Negative regulation of response to stimulus | 10 | 0.66 | 0.00033 | IL2,IL12B,HAVCR2,STAT2,LILRB1,IFNB1,OAS1,IFI35,TNF,SOCS1 |
| GO Process | GO:0001932 | Regulation of protein phosphorylation | 8 | 0.66 | 0.0012 | IL2,IFNG,IL12B,C3,STAT2,IFNB1,TNF,SOCS1 |
| GO Process | GO:0045597 | Positive regulation of cell differentiation | 7 | 0.66 | 0.0021 | IL2,IFNG,IL12B,NFKBIZ,CXCL9,TNF,SOCS1 |
| GO Process | GO:0030098 | Lymphocyte differentiation | 4 | 0.66 | 0.0115 | IL2,IL12B,TNFSF13B,IFNB1 |
| GO Process | GO:0051928 | Positive regulation of calcium ion transport | 3 | 0.66 | 0.018 | CXCL10,CXCL9,CCL4 |

| | | | | | | |
|---|---|---|---|---|---|---|
| GO Process | GO:0002573 | Myeloid leukocyte differentiation | 3 | 0.66 | 0.0184 | IFNG,TNF,SOCS1 |
| GO Process | GO:0002230 | Positive regulation of defense response to virus by host | 2 | 0.66 | 0.0275 | IL12B,LILRB1 |
| GO Process | GO:0050793 | Regulation of developmental process | 13 | 0.64 | 4.51E-05 | IL2,IFNG,IL12B,C3,CXCL10,STAT2,LILRB1,NFKBIZ,CXCL9,TNFSF13B,IFNB1,TNF,SOCS1 |
| GO Process | GO:0043270 | Positive regulation of ion transport | 4 | 0.64 | 0.0124 | IFNG,CXCL10,CXCL9,CCL4 |
| GO Process | GO:0007165 | Signal transduction | 20 | 0.63 | 2.29E-08 | IL2,IFNG,IL12B,TNFSF10,C3,CXCL10,HAVCR2,STAT2,LILRB1,NFKBIZ,CXCL9,NFKB2,TNFSF13B,IFNB1,CCL8,OAS1,TNF,CCL4,SOCS1,IFIH1 |
| GO Process | GO:0048660 | Regulation of smooth muscle cell proliferation | 3 | 0.63 | 0.0212 | IFNG,IL12B,TNF |
| GO Process | GO:0043269 | Regulation of ion transport | 6 | 0.62 | 0.005 | IFNG,CXCL10,LILRB1,CXCL9,TNF,CCL4 |
| GO Process | GO:0001914 | Regulation of T cell mediated cytotoxicity | 2 | 0.62 | 0.0346 | IL12B,LILRB1 |
| GO Process | GO:0042981 | Regulation of apoptotic process | 9 | 0.61 | 0.0011 | IL2,IFNG,IL12B,TNFSF10,C3,CXCL10,LILRB1,IFNB1,TNF |
| GO Process | GO:0008284 | Positive regulation of cell population proliferation | 7 | 0.61 | 0.0032 | IL2,IFNG,IL12B,CXCL10,HAVCR2,TNFSF13B,TNF |
| GO Process | GO:0008283 | Cell population proliferation | 6 | 0.61 | 0.0054 | IL2,IL12B,LILRB1,TNFSF13B,IFNB1,TNF |
| GO Process | GO:0042742 | Defense response to bacterium | 4 | 0.61 | 0.015 | IL12B,HAVCR2,OAS1,TNF |
| GO Process | GO:0001959 | Regulation of cytokine-mediated signaling pathway | 3 | 0.6 | 0.025 | STAT2,OAS1,SOCS1 |
| GO Process | GO:0032728 | Positive regulation of interferon-beta production | 2 | 0.6 | 0.0378 | OAS1,IFIH1 |
| GO Process | GO:0009967 | Positive regulation of signal transduction | 9 | 0.59 | 0.0014 | IL12B,TNFSF10,C3,HAVCR2,IFNB1,CCL8,IFI35,TNF,CCL4 |
| GO Process | GO:0001934 | Positive regulation of protein phosphorylation | 6 | 0.59 | 0.0067 | IL2,IFNG,IL12B,C3,IFNB1,TNF |
| GO Process | GO:0050850 | Positive regulation of calcium-mediated signaling | 2 | 0.59 | 0.0393 | TNF,CCL4 |
| GO Process | GO:0051239 | Regulation of multicellular organismal process | 13 | 0.58 | 0.00012 | IL2,IFNG,IL12B,C3,CXCL10,HAVCR2,LILRB1,NFKBIZ,IFNB1,OAS1,TNF,SOCS1,IFIH1 |
| GO Process | GO:1902533 | Positive regulation of intracellular signal transduction | 7 | 0.58 | 0.0042 | IL12B,TNFSF10,HAVCR2,CCL8,IFI35,TNF,CCL4 |
| GO Process | GO:0051336 | Regulation of hydrolase activity | 7 | 0.58 | 0.0045 | IFNG,TNFSF10,C3,CCL8,OAS1,TNF,CCL4 |
| GO Process | GO:0030335 | Positive regulation of cell migration | 5 | 0.58 | 0.0118 | IFNG,CXCL10,CCL8,TNF,CCL4 |
| GO Process | GO:0001937 | Negative regulation of endothelial cell proliferation | 2 | 0.58 | 0.0408 | IL12B,TNF |
| GO Process | GO:0002548 | Monocyte chemotaxis | 2 | 0.58 | 0.0408 | CCL8,CCL4 |
| GO Process | GO:0032735 | Positive regulation of interleukin-12 production | 2 | 0.58 | 0.0408 | IFNG,IL12B |
| GO Process | GO:0045429 | Positive regulation of nitric oxide biosynthetic process | 2 | 0.58 | 0.0408 | IFNG,TNF |
| GO Process | GO:0051281 | Positive regulation of release of sequestered calcium ion into cytosol | 2 | 0.58 | 0.0408 | CXCL10,CXCL9 |
| GO Process | GO:2000403 | Positive regulation of lymphocyte migration | 2 | 0.58 | 0.0408 | CXCL10,CCL4 |
| GO Process | GO:0032689 | Negative regulation of interferon-gamma production | 2 | 0.58 | 0.0417 | HAVCR2,LILRB1 |
| GO Process | GO:0051048 | Negative regulation of secretion | 3 | 0.57 | 0.0304 | IL12B,LILRB1,TNF |
| GO Process | GO:0030890 | Positive regulation of B cell proliferation | 2 | 0.57 | 0.0433 | IL2,TNFSF13B |
| GO Process | GO:0046427 | Positive regulation of receptor signaling pathway via JAK-STAT | 2 | 0.57 | 0.0433 | IL12B,TNF |
| GO Process | GO:0032814 | Regulation of natural killer cell activation | 2 | 0.57 | 0.0444 | IL12B,HAVCR2 |
| GO Process | GO:1901701 | Cellular response to oxygen-containing compound | 7 | 0.55 | 0.0056 | IL12B,CXCL10,HAVCR2,LILRB1,CXCL9,TNF,SOCS1 |
| GO Process | GO:1904064 | Positive regulation of cation transmembrane transport | 3 | 0.55 | 0.0337 | IFNG,CXCL10,CXCL9 |
| GO Process | GO:0046890 | Regulation of lipid biosynthetic process | 3 | 0.55 | 0.0342 | IFNG,C3,TNF |
| GO Process | GO:0032715 | Negative regulation of interleukin-6 production | 2 | 0.55 | 0.0478 | HAVCR2,TNF |
| GO Process | GO:0046640 | Regulation of alpha-beta T cell proliferation | 2 | 0.55 | 0.0495 | IL2,IL12B |
| GO Process | GO:0048522 | Positive regulation of cellular process | 20 | 0.54 | 3.47E-07 | IL2,IFNG,IL12B,TNFSF10,C3,CXCL10,HAVCR2,STAT2,LILRB1,NFKBIZ,CXCL9,NFKB2,TNFSF13B,IFNB1,CCL8,OAS1,IFI35,TNF,CCL4,SOCS1 |
| GO Process | GO:0007267 | Cell-cell signaling | 7 | 0.54 | 0.0063 | IL2,TNFSF10,C3,CXCL10,CXCL9,CCL8,CCL4 |
| GO Process | GO:0009966 | Regulation of signal transduction | 13 | 0.53 | 0.00027 | IFNG,IL12B,TNFSF10,C3,HAVCR2,STAT2,IFNB1,CCL8,OAS1,IFI35,TNF,CCL4,SOCS1 |
| GO Process | GO:0034762 | Regulation of transmembrane transport | 5 | 0.53 | 0.017 | IFNG,C3,CXCL10,CXCL9,TNF |
| GO Process | GO:0051345 | Positive regulation of hydrolase activity | 5 | 0.53 | 0.0172 | IFNG,TNFSF10,CCL8,TNF,CCL4 |
| GO Process | GO:0048518 | Positive regulation of biological process | 21 | 0.52 | 1.34E-07 | IL2,IFNG,IL12B,TNFSF10,C3,CXCL10,HAVCR2,STAT2,LILRB1,NFKBIZ,CXCL9,NFKB2,TNFSF13B,IFNB1,CCL8,OAS1,IFI35,TNF,CCL4,SOCS1,IFIH1 |
| GO Process | GO:0050790 | Regulation of catalytic activity | 11 | 0.52 | 0.0012 | IL2,IFNG,IL12B,TNFSF10,C3,CXCL10,CCL8,OAS1,TNF,CCL4,SOCS1 |
| GO Process | GO:0010604 | Positive regulation of macromolecule metabolic process | 14 | 0.5 | 0.00026 | IL2,IFNG,IL12B,TNFSF10,C3,CXCL10,HAVCR2,STAT2,LILRB1,NFKB2,IFNB1,OAS1,TNF,IFIH1 |
| GO Process | GO:0043085 | Positive regulation of catalytic activity | 7 | 0.49 | 0.0108 | IL2,IFNG,IL12B,TNFSF10,CCL8,TNF,CCL4 |
| GO Process | GO:0016477 | Cell migration | 6 | 0.49 | 0.0158 | IL12B,CXCL10,CXCL9,CCL8,TNF,CCL4 |
| GO Process | GO:0051050 | Positive regulation of transport | 6 | 0.48 | 0.0166 | IFNG,C3,CXCL10,CXCL9,TNF,CCL4 |
| GO Process | GO:0045944 | Positive regulation of transcription by RNA polymerase II | 7 | 0.46 | 0.0134 | IL2,CXCL10,STAT2,LILRB1,NFKB2,IFNB1,TNF |
| GO Process | GO:0045936 | Negative regulation of phosphate metabolic process | 4 | 0.46 | 0.0413 | IL2,IFNG,TNF,SOCS1 |
| GO Process | GO:0065009 | Regulation of molecular function | 12 | 0.45 | 0.002 | IL2,IFNG,IL12B,TNFSF10,C3,CXCL10,HAVCR2,CCL8,OAS1,TNF,CCL4,SOCS1 |
| GO Process | GO:0050896 | Response to stimulus | 22 | 0.44 | 3.76E-07 | IL2,IFNG,IL12B,TNFSF10,C3,CXCL10,HAVCR2,STAT2,LILRB1,NFKBIZ,CXCL9,SLAMF7,NFKB2,TNFSF13B,IFNB1,CCL8,OAS1,IFI35,TNF,CCL4,SOCS1,IFIH1 |
| GO Process | GO:0010605 | Negative regulation of macromolecule metabolic process | 11 | 0.44 | 0.0037 | IL2,IFNG,IL12B,C3,HAVCR2,LILRB1,NFKB2,IFNB1,OAS1,TNF,SOCS1 |
| GO Process | GO:0051049 | Regulation of transport | 8 | 0.41 | 0.0172 | IFNG,IL12B,C3,CXCL10,LILRB1,CXCL9,TNF,CCL4 |
| GO Process | GO:1901698 | Response to nitrogen compound | 6 | 0.41 | 0.0318 | TNFSF10,STAT2,IFNB1,TNF,SOCS1,IFIH1 |
| GO Process | GO:0048519 | Negative regulation of biological process | 16 | 0.4 | 0.00081 | IL2,IFNG,IL12B,C3,CXCL10,HAVCR2,STAT2,LILRB1,NFKB2,IFNB1,CCL8,OAS1,IFI35,TNF,SOCS1,IFIH1 |
| GO Process | GO:0010468 | Regulation of gene expression | 15 | 0.39 | 0.0016 | IL2,IFNG,IL12B,C3,CXCL10,HAVCR2,STAT2,LILRB1,NFKBIZ,NFKB2,IFNB1,OAS1,TNF,SOCS1,IFIH1 |
| GO Process | GO:0043549 | Regulation of kinase activity | 5 | 0.39 | 0.0495 | IL2,IFNG,IL12B,TNF,SOCS1 |
| GO Process | GO:0031325 | Positive regulation of cellular metabolic process | 11 | 0.38 | 0.0097 | IL2,IFNG,IL12B,C3,CXCL10,STAT2,LILRB1,NFKB2,IFNB1,OAS1,TNF |
| GO Process | GO:0051173 | Positive regulation of nitrogen compound metabolic process | 11 | 0.37 | 0.0109 | IL2,IFNG,IL12B,TNFSF10,C3,CXCL10,STAT2,LILRB1,NFKB2,IFNB1,TNF |
| GO Process | GO:0051247 | Positive regulation of protein metabolic process | 7 | 0.37 | 0.0351 | IL2,IFNG,IL12B,TNFSF10,C3,IFNB1,TNF |
| GO Process | GO:0048523 | Negative regulation of cellular process | 14 | 0.36 | 0.0049 | IL2,IFNG,IL12B,CXCL10,HAVCR2,STAT2,LILRB1,NFKB2,IFNB1,CCL8,OAS1,IFI35,TNF,SOCS1 |
| GO Process | GO:0048513 | Animal organ development | 11 | 0.36 | 0.013 | IL2,IFNG,IL12B,TNFSF10,CXCL10,LILRB1,NFKB2,TNFSF13B,IFNB1,TNF,SOCS1 |
| GO Process | GO:0051128 | Regulation of cellular component organization | 9 | 0.36 | 0.0231 | IL2,IFNG,TNFSF10,C3,CXCL10,STAT2,LILRB1,CXCL9,TNF |
| GO Process | GO:0010557 | Positive regulation of macromolecule biosynthetic process | 8 | 0.36 | 0.0295 | IL2,IFNG,CXCL10,STAT2,LILRB1,NFKB2,IFNB1,TNF |
| GO Process | GO:0007186 | G protein-coupled receptor signaling pathway | 6 | 0.36 | 0.0495 | IL2,C3,CXCL10,CXCL9,CCL8,CCL4 |
| GO Process | GO:0048731 | System development | 12 | 0.34 | 0.0131 | IL2,IFNG,IL12B,TNFSF10,C3,HAVCR2,LILRB1,NFKB2,TNFSF13B,IFNB1,TNF,SOCS1 |

| | | | | | | |
|---|---|---|---|---|---|---|
| GO Process | GO:0031328 | Positive regulation of cellular biosynthetic process | 8 | 0.34 | 0.0402 | IL2,IFNG,CXCL10,STAT2,LILRB1,NFKB2,IFNB1,TNF |
| GO Process | GO:0045935 | Positive regulation of nucleobase-containing compound metabolic proce | 8 | 0.33 | 0.0414 | IL2,IFNG,CXCL10,STAT2,LILRB1,NFKB2,IFNB1,TNF |
| GO Process | GO:0060255 | Regulation of macromolecule metabolic process | 16 | 0.32 | 0.005 | IL2,IFNG,IL12B,TNFSF10,C3,CXCL10,HAVCR2,STAT2,LILRB1,NFKBIZ,NFKB2,IFNB1,OAS1,TNF,SOCS1,IFIH1 |
| GO Process | GO:0051246 | Regulation of protein metabolic process | 9 | 0.31 | 0.0438 | IL2,IFNG,IL12B,TNFSF10,C3,STAT2,IFNB1,TNF,SOCS1 |
| GO Process | GO:0050794 | Regulation of cellular process | 21 | 0.27 | 0.0031 | IL2,IFNG,IL12B,TNFSF10,C3,CXCL10,HAVCR2,STAT2,LILRB1,NFKBIZ,CXCL9,NFKB2,TNFSF13B,IFNB1,CCL8,OAS1,IFI35,TNF,CCL4,SOCS1,IFIH1 |
| GO Process | GO:0051171 | Regulation of nitrogen compound metabolic process | 14 | 0.27 | 0.0304 | IL2,IFNG,IL12B,TNFSF10,C3,CXCL10,HAVCR2,STAT2,LILRB1,NFKB2,IFNB1,OAS1,TNF,SOCS1 |
| GO Process | GO:0048856 | Anatomical structure development | 13 | 0.27 | 0.0378 | IL2,IFNG,IL12B,TNFSF10,C3,CXCL10,HAVCR2,LILRB1,NFKB2,TNFSF13B,IFNB1,TNF,SOCS1 |
| GO Process | GO:0080090 | Regulation of primary metabolic process | 14 | 0.26 | 0.04 | IL2,IFNG,IL12B,TNFSF10,C3,CXCL10,HAVCR2,STAT2,LILRB1,NFKB2,IFNB1,OAS1,TNF,SOCS1 |
| GO Function | GO:0005125 | Cytokine activity | 11 | 3.25 | 2.68E-12 | IL2,IFNG,IL12B,TNFSF10,CXCL10,CXCL9,TNFSF13B,IFNB1,CCL8,TNF,CCL4 |
| GO Function | GO:0005126 | Cytokine receptor binding | 11 | 3.01 | 6.12E-12 | IL2,IFNG,IL12B,TNFSF10,CXCL10,CXCL9,TNFSF13B,IFNB1,CCL8,TNF,CCL4 |
| GO Function | GO:0008009 | Chemokine activity | 4 | 1.51 | 0.00019 | CXCL10,CXCL9,CCL8,CCL4 |
| GO Function | GO:0005102 | Signaling receptor binding | 14 | 1.13 | 3.38E-08 | IL2,IFNG,IL12B,TNFSF10,C3,CXCL10,LILRB1,CXCL9,TNFSF13B,IFNB1,CCL8,TNF,CCL4,SOCS1 |
| GO Function | GO:0001664 | G protein-coupled receptor binding | 6 | 1.08 | 0.0004 | IL2,C3,CXCL10,CXCL9,CCL8,CCL4 |
| GO Function | GO:0005164 | Tumor necrosis factor receptor binding | 3 | 1.04 | 0.0035 | TNFSF10,TNFSF13B,TNF |
| GO Function | GO:0048248 | CXCR3 chemokine receptor binding | 2 | 0.9 | 0.0094 | CXCL10,CXCL9 |
| GO Function | GO:0098772 | Molecular function regulator activity | 13 | 0.78 | 1.42E-05 | IL2,IFNG,IL12B,TNFSF10,C3,CXCL10,CXCL9,TNFSF13B,IFNB1,CCL8,TNF,CCL4,SOCS1 |
| GO Function | GO:0005515 | Protein binding | 18 | 0.31 | 0.0083 | IL2,IFNG,IL12B,TNFSF10,C3,CXCL10,STAT2,LILRB1,CXCL9,SLAMF7,TNFSF13B,IFNB1,CCL8,IFI35,TNF,CCL4,SOCS1,IFIH1 |
| GO Component | GO:0005576 | Extracellular region | 15 | 0.4 | 0.0059 | IL2,IFNG,IL12B,TNFSF10,C3,CXCL10,LILRB1,CXCL9,TNFSF13B,IFNB1,CCL8,OAS1,IFI35,TNF,CCL4 |
| GO Component | GO:0005615 | Extracellular space | 13 | 0.42 | 0.0059 | IL2,IFNG,IL12B,TNFSF10,C3,CXCL10,CXCL9,TNFSF13B,IFNB1,CCL8,IFI35,TNF,CCL4 |
| GO Component | GO:0009897 | External side of plasma membrane | 6 | 0.74 | 0.0059 | IL12B,CXCL10,LILRB1,CXCL9,SLAMF7,TNF |
| GO Component | GO:0009986 | Cell surface | 8 | 0.62 | 0.0059 | IL12B,C3,CXCL10,HAVCR2,LILRB1,CXCL9,SLAMF7,TNF |
| STRING cluster | CL:15536 | Mixed, incl, Chemokine-mediated signaling pathway, and Adaptive imr | 6 | 1.37 | 0.00014 | CXCL10,HAVCR2,CXCL9,SLAMF7,CCL8,CCL4 |
| STRING cluster | CL:15743 | Chemokine receptors bind chemokines | 4 | 1.58 | 0.00016 | CXCL10,CXCL9,CCL8,CCL4 |
| STRING cluster | CL:15942 | JAK-STAT signaling pathway | 5 | 1.43 | 0.00016 | IL2,IFNG,IL12B,STAT2,IFNB1 |
| STRING cluster | CL:15943 | JAK-STAT signaling pathway | 4 | 1.08 | 0.0016 | IL2,IFNG,IL12B,IFNB1 |
| STRING cluster | CL:15945 | JAK-STAT signaling pathway | 3 | 0.73 | 0.0172 | IL2,IFNG,IFNB1 |
| STRING cluster | CL:16585 | Mixed, incl, Novel intracellular components of RIG-I-like receptor path | 3 | 0.66 | 0.0245 | NFKB2,TNF,IFIH1 |
| STRING cluster | CL:16017 | Mixed, incl, Type III interferon signaling pathway, and Interferon recep | 2 | 0.68 | 0.0289 | IFNG,IFNB1 |
| STRING cluster | CL:15784 | Mixed, incl, CXC chemokine, and CXC chemokine receptor 1/2 | 2 | 0.64 | 0.0359 | CCL8,CCL4 |
| STRING cluster | CL:15746 | CXC Chemokine domain, and Regulation of dendritic cell dendrite asse | 2 | 0.62 | 0.0385 | CXCL10,CXCL9 |
| KEGG | hsa04060 | Cytokine-cytokine receptor interaction | 11 | 3.06 | 1.32E-12 | IL2,IFNG,IL12B,TNFSF10,CXCL10,CXCL9,TNFSF13B,IFNB1,CCL8,TNF,CCL4 |
| KEGG | hsa05164 | Influenza A | 9 | 3.27 | 1.79E-11 | IFNG,IL12B,TNFSF10,CXCL10,STAT2,IFNB1,OAS1,TNF,IFIH1 |
| KEGG | hsa04061 | Viral protein interaction with cytokine and cytokine receptor | 7 | 3.06 | 1.54E-09 | IL2,TNFSF10,CXCL10,CXCL9,CCL8,TNF,CCL4 |
| KEGG | hsa04380 | Osteoclast differentiation | 7 | 2.77 | 5.14E-09 | IFNG,STAT2,LILRB1,NFKB2,IFNB1,TNF,SOCS1 |
| KEGG | hsa04620 | Toll-like receptor signaling pathway | 6 | 2.47 | 8.21E-08 | IL12B,CXCL10,CXCL9,IFNB1,TNF,CCL4 |
| KEGG | hsa05142 | Chagas disease | 6 | 2.49 | 8.21E-08 | IL2,IFNG,IL12B,C3,IFNB1,TNF |
| KEGG | hsa05162 | Measles | 6 | 2.11 | 4.26E-07 | IL2,IL12B,STAT2,IFNB1,OAS1,IFIH1 |
| KEGG | hsa04622 | RIG-I-like receptor signaling pathway | 5 | 2.3 | 6.82E-07 | IL12B,CXCL10,IFNB1,TNF,IFIH1 |
| KEGG | hsa04630 | JAK-STAT signaling pathway | 6 | 1.98 | 7.26E-07 | IL2,IFNG,IL12B,STAT2,IFNB1,SOCS1 |
| KEGG | hsa05160 | Hepatitis C | 6 | 1.98 | 7.26E-07 | IFNG,CXCL10,STAT2,IFNB1,OAS1,TNF |
| KEGG | hsa05168 | Herpes simplex virus 1 infection | 8 | 1.44 | 9.26E-07 | IFNG,IL12B,C3,STAT2,IFNB1,OAS1,TNF,IFIH1 |
| KEGG | hsa05169 | Epstein-Barr virus infection | 6 | 1.79 | 1.74E-06 | CXCL10,STAT2,NFKB2,IFNB1,OAS1,TNF |
| KEGG | hsa05330 | Allograft rejection | 4 | 2.28 | 2.16E-06 | IL2,IFNG,IL12B,TNF |
| KEGG | hsa04625 | C-type lectin receptor signaling pathway | 5 | 1.98 | 2.40E-06 | IL2,IL12B,STAT2,NFKB2,TNF |
| KEGG | hsa04940 | Type I diabetes mellitus | 4 | 2.21 | 2.83E-06 | IL2,IFNG,IL12B,TNF |
| KEGG | hsa05134 | Legionellosis | 4 | 1.92 | 1.07E-05 | IL12B,C3,NFKB2,TNF |
| KEGG | hsa04217 | Necroptosis | 5 | 1.64 | 1.19E-05 | IFNG,TNFSF10,STAT2,IFNB1,TNF |
| KEGG | hsa05321 | Inflammatory bowel disease | 4 | 1.88 | 1.24E-05 | IL2,IFNG,IL12B,TNF |
| KEGG | hsa05152 | Tuberculosis | 5 | 1.54 | 1.86E-05 | IFNG,IL12B,C3,IFNB1,TNF |
| KEGG | hsa05140 | Leishmaniasis | 4 | 1.76 | 2.02E-05 | IFNG,IL12B,C3,TNF |
| KEGG | hsa04062 | Chemokine signaling pathway | 5 | 1.44 | 2.98E-05 | CXCL10,STAT2,CXCL9,CCL8,CCL4 |
| KEGG | hsa04064 | NF-kappa B signaling pathway | 4 | 1.47 | 7.87E-05 | NFKB2,TNFSF13B,TNF,CCL4 |
| KEGG | hsa05145 | Toxoplasmosis | 4 | 1.46 | 8.12E-05 | IFNG,IL12B,TNF,SOCS1 |
| KEGG | hsa04650 | Natural killer cell mediated cytotoxicity | 4 | 1.35 | 0.00014 | IFNG,TNFSF10,IFNB1,TNF |
| KEGG | hsa05143 | African trypanosomiasis | 3 | 1.56 | 0.00014 | IFNG,IL12B,TNF |
| KEGG | hsa05332 | Graft-versus-host disease | 3 | 1.56 | 0.00014 | IL2,IFNG,TNF |
| KEGG | hsa05161 | Hepatitis B | 4 | 1.16 | 0.00036 | STAT2,IFNB1,TNF,IFIH1 |
| KEGG | hsa04621 | NOD-like receptor signaling pathway | 4 | 1.1 | 0.00049 | STAT2,IFNB1,OAS1,TNF |
| KEGG | hsa04623 | Cytosolic DNA-sensing pathway | 3 | 1.25 | 0.00058 | CXCL10,IFNB1,CCL4 |
| KEGG | hsa05133 | Pertussis | 3 | 1.16 | 0.0009 | IL12B,C3,TNF |
| KEGG | hsa04658 | Th1 and Th2 cell differentiation | 3 | 1.08 | 0.0013 | IL2,IFNG,IL12B |
| KEGG | hsa05323 | Rheumatoid arthritis | 3 | 1.09 | 0.0013 | IFNG,TNFSF13B,TNF |
| KEGG | hsa04657 | IL-17 signaling pathway | 3 | 1.05 | 0.0015 | IFNG,CXCL10,TNF |
| KEGG | hsa05322 | Systemic lupus erythematosus | 3 | 1.04 | 0.0016 | IFNG,C3,TNF |
| KEGG | hsa04660 | T cell receptor signaling pathway | 3 | 1 | 0.0019 | IL2,IFNG,TNF |
| KEGG | hsa05146 | Amoebiasis | 3 | 1 | 0.0019 | IFNG,IL12B,TNF |
| KEGG | hsa05200 | Pathways in cancer | 5 | 0.72 | 0.0021 | IL2,IFNG,IL12B,STAT2,NFKB2 |

| | | | | | | |
|---|---|---|---|---|---|---|
| KEGG | hsa04668 | TNF signaling pathway | 3 | 0.95 | 0.0024 | CXCL10,IFNB1,TNF |
| KEGG | hsa05135 | Yersinia infection | 3 | 0.9 | 0.0032 | IL2,IFNB1,TNF |
| KEGG | hsa04672 | Intestinal immune network for IgA production | 2 | 0.82 | 0.0096 | IL2,TNFSF13B |
| KEGG | hsa05167 | Kaposi sarcoma-associated herpesvirus infection | 3 | 0.69 | 0.0097 | C3,STAT2,IFNB1 |
| KEGG | hsa04930 | Type II diabetes mellitus | 2 | 0.81 | 0.01 | TNF,SOCS1 |
| KEGG | hsa05144 | Malaria | 2 | 0.81 | 0.0102 | IFNG,TNF |
| KEGG | hsa05166 | Human T-cell leukemia virus 1 infection | 3 | 0.64 | 0.0126 | IL2,NFKB2,TNF |
| KEGG | hsa05131 | Shigellosis | 3 | 0.63 | 0.0135 | C3,IFNB1,TNF |
| KEGG | hsa05163 | Human cytomegalovirus infection | 3 | 0.63 | 0.0135 | IFNB1,TNF,CCL4 |
| KEGG | hsa04612 | Antigen processing and presentation | 2 | 0.7 | 0.0174 | IFNG,TNF |
| KEGG | hsa04350 | TGF-beta signaling pathway | 2 | 0.57 | 0.0335 | IFNG,TNF |
| KEGG | hsa04659 | Th17 cell differentiation | 2 | 0.55 | 0.0378 | IL2,IFNG |
| KEGG | hsa05165 | Human papillomavirus infection | 3 | 0.47 | 0.0378 | STAT2,IFNB1,TNF |
| Reactome | HSA-1280215 | Cytokine Signaling in Immune system | 14 | 2.13 | 3.68E-12 | IL2,IFNG,IL12B,CXCL10,HAVCR2,STAT2,NFKB2,TNFSF13B,IFNB1,OAS1,IFI35,TNF,CCL4,SOCS1 |
| Reactome | HSA-909733 | Interferon alpha/beta signaling | 5 | 1.91 | 1.06E-05 | IL2,IFNG,IL12B,C3,CXCL10,HAVCR2,STAT2,LILRB1,SLAMF7,NFKB2,TNFSF13B,IFNB1,OAS1,IFI35,TNF,CCL4,SOCS1,IFIH1 |
| Reactome | HSA-449147 | Signaling by Interleukins | 10 | 1.87 | 1.77E-08 | IL2,IFNG,IL12B,CXCL10,HAVCR2,STAT2,NFKB2,TNF,CCL4,SOCS1 |
| Reactome | HSA-6783783 | Interleukin-10 signaling | 4 | 1.64 | 9.06E-05 | STAT2,IFNB1,OAS1,IFI35,SOCS1 |
| Reactome | HSA-913531 | Interferon Signaling | 6 | 1.47 | 3.48E-05 | IFNG,STAT2,IFNB1,OAS1,IFI35,SOCS1 |
| Reactome | HSA-168256 | Immune System | 18 | 1.25 | 6.55E-12 | IL12B,CXCL10,TNF,CCL4 |
| Reactome | HSA-912694 | Regulation of IFNA/IFNB signaling | 3 | 1.23 | 0.0013 | STAT2,IFNB1,SOCS1 |
| Reactome | HSA-380108 | Chemokine receptors bind chemokines | 3 | 0.8 | 0.0113 | CXCL10,CXCL9,CCL4 |
| Reactome | HSA-8877330 | RUNX1 and FOXP3 control the development of regulatory T lymphocy | 2 | 0.77 | 0.0178 | C3,CXCL10,CXCL9,CCL4 |
| Reactome | HSA-877312 | Regulation of IFNG signaling | 2 | 0.69 | 0.0269 | IL2,IFNG |
| Reactome | HSA-918233 | TRAF3-dependent IRF activation pathway | 2 | 0.69 | 0.0269 | NFKB2,IFNB1,IFIH1 |
| Reactome | HSA-168928 | DDX58/IFIH1-mediated induction of interferon-alpha/beta | 3 | 0.68 | 0.0217 | IFNG,OAS1,SOCS1 |
| Reactome | HSA-375276 | Peptide ligand-binding receptors | 4 | 0.67 | 0.0159 | IFNG,SOCS1 |
| Reactome | HSA-877300 | Interferon gamma signaling | 3 | 0.63 | 0.0269 | IFNB1,IFIH1 |
| Reactome | HSA-5668541 | TNFR2 non-canonical NF-kB pathway | 3 | 0.61 | 0.0293 | NFKB2,TNFSF13B,TNF |
| Reactome | HSA-6785807 | Interleukin-4 and Interleukin-13 signaling | 3 | 0.58 | 0.0343 | IL12B,TNF,SOCS1 |
| Reactome | HSA-933542 | TRAF6 mediated NF-kB activation | 2 | 0.57 | 0.0493 | C3,CXCL10,CXCL9,CCL4 |
| Reactome | HSA-9660826 | Purinergic signaling in leishmaniasis infection | 2 | 0.57 | 0.0493 | STAT2,IFNB1,IFIH1 |
| Reactome | HSA-9705671 | SARS-CoV-2 activates/modulates innate and adaptive immune response | 3 | 0.52 | 0.0482 | C3,LILRB1,SLAMF7 |
| Reactome | HSA-198933 | Immunoregulatory interactions between a Lymphoid and a non-Lympho | 3 | 0.52 | 0.0493 | NFKB2,IFIH1 |
| Reactome | HSA-418594 | G alpha (i) signalling events | 4 | 0.47 | 0.0482 | C3,NFKB2 |
| WikiPathways | WP5095 | Overview of proinflammatory and profibrotic mediators | 10 | 4.19 | 8.18E-14 | IL2,IFNG,IL12B,CXCL10,CXCL9,TNFSF13B,IFNB1,CCL8,TNF,CCL4 |
| WikiPathways | WP5115 | Network map of SARS-CoV-2 signaling pathway | 10 | 3.14 | 7.83E-12 | IFNG,TNFSF10,CXCL10,CXCL9,NFKB2,IFNB1,CCL8,TNF,CCL4,IFIH1 |
| WikiPathways | WP619 | Type II interferon signaling | 7 | 4.37 | 7.83E-12 | IFNG,CXCL10,STAT2,CXCL9,IFNB1,OAS1,SOCS1 |
| WikiPathways | WP5039 | SARS-CoV-2 innate immunity evasion and cell-specific immune respon | 7 | 3.55 | 2.26E-10 | CXCL10,HAVCR2,STAT2,CXCL9,IFNB1,TNF,CCL4 |
| WikiPathways | WP4630 | Measles virus infection | 7 | 2.54 | 2.10E-08 | IL2,IL12B,STAT2,NFKB2,IFNB1,OAS1,IFIH1 |
| WikiPathways | WP530 | Cytokines and inflammatory response | 5 | 3.14 | 2.30E-08 | IL2,IFNG,IL12B,IFNB1,TNF |
| WikiPathways | WP5218 | Extrafollicular and follicular B cell activation by SARS-CoV-2 | 6 | 2.8 | 2.49E-08 | IL2,IFNG,C3,SLAMF7,TNFSF13B,TNF |
| WikiPathways | WP4298 | Acute viral myocarditis | 6 | 2.61 | 5.61E-08 | IL2,IFNG,IL12B,NFKB2,TNF,SOCS1 |
| WikiPathways | WP2328 | Allograft rejection | 6 | 2.58 | 6.07E-08 | IL2,IFNG,IL12B,C3,CXCL9,TNF |
| WikiPathways | WP75 | Toll-like receptor signaling pathway | 6 | 2.41 | 1.27E-07 | IL12B,CXCL10,CXCL9,IFNB1,TNF,CCL4 |
| WikiPathways | WP3865 | Novel intracellular components of RIG-I-like receptor pathway | 5 | 2.39 | 5.50E-07 | IFNG,CXCL10,IFNB1,TNF,IFIH1 |
| WikiPathways | WP4197 | Immune response to tuberculosis | 4 | 2.45 | 1.30E-06 | STAT2,OAS1,IFI35,SOCS1 |
| WikiPathways | WP4494 | Selective expression of chemokine receptors during T-cell polarization | 4 | 2.28 | 2.79E-06 | IL2,IFNG,IL12B,CCL4 |
| WikiPathways | WP5098 | T-cell activation SARS-CoV-2 | 5 | 2.01 | 2.88E-06 | IL2,IFNG,IL12B,IFNB1,TNF |
| WikiPathways | WP5088 | Prostaglandin signaling | 4 | 2.22 | 3.48E-06 | IFNG,CXCL10,CXCL9,TNF |
| WikiPathways | WP5285 | Immune infiltration in pancreatic cancer | 4 | 2.09 | 6.17E-06 | IL2,IFNG,IL12B,TNF |
| WikiPathways | WP2431 | Spinal cord injury | 5 | 1.77 | 8.54E-06 | IL2,IFNG,CXCL10,TNFSF13B,TNF |
| WikiPathways | WP4136 | Fibrin complement receptor 3 signaling pathway | 4 | 2.01 | 8.71E-06 | IL12B,CXCL10,IFNB1,TNF |
| WikiPathways | WP2873 | Aryl hydrocarbon receptor pathway | 4 | 1.98 | 9.78E-06 | IL2,IFNG,IL12B,TNF |
| WikiPathways | WP3611 | Photodynamic therapy-induced AP-1 survival signaling | 4 | 1.9 | 1.38E-05 | IL2,IFNG,TNFSF10,TNF |
| WikiPathways | WP4754 | IL-18 signaling pathway | 6 | 1.41 | 1.67E-05 | IFNG,IL12B,NFKBIZ,NFKB2,TNF,CCL4 |
| WikiPathways | WP4891 | COVID-19 adverse outcome pathway | 3 | 1.91 | 3.45E-05 | IL2,CXCL10,TNF |
| WikiPathways | WP176 | Folate metabolism | 4 | 1.69 | 3.65E-05 | IL2,IFNG,NFKB2,TNF |
| WikiPathways | WP4705 | Pathways of nucleic acid metabolism and innate immune sensing | 3 | 1.88 | 3.76E-05 | IFNB1,OAS1,IFIH1 |
| WikiPathways | WP4341 | Non-genomic actions of 1,25 dihydroxyvitamin D3 | 4 | 1.62 | 4.90E-05 | IFNG,STAT2,NFKB2,TNF |
| WikiPathways | WP262 | Ebstein-Barr virus LMP1 signaling | 3 | 1.69 | 9.26E-05 | NFKB2,IFNB1,TNF |
| WikiPathways | WP5044 | Kynurenine pathway and links to cell senescence | 3 | 1.69 | 9.26E-05 | IFNG,IFNB1,TNF |
| WikiPathways | WP5092 | Interactions of natural killer cells in pancreatic cancer | 3 | 1.61 | 0.00013 | IFNG,TNF,CCL4 |
| WikiPathways | WP4559 | Interactions between immune cells and microRNAs in tumor microenvi | 3 | 1.6 | 0.00014 | CXCL10,NFKB2,SOCS1 |
| WikiPathways | WP4868 | Type I interferon induction and signaling during SARS-CoV-2 infection | 3 | 1.54 | 0.00018 | STAT2,OAS1,IFIH1 |
| WikiPathways | WP4912 | SARS coronavirus and innate immunity | 3 | 1.54 | 0.00018 | STAT2,IFNB1,IFIH1 |
| WikiPathways | WP3893 | Development and heterogeneity of the ILC family | 3 | 1.53 | 0.00019 | IFNG,IL12B,TNF |
| WikiPathways | WP4880 | Host-pathogen interaction of human coronaviruses - interferon inductio | 3 | 1.51 | 0.0002 | STAT2,OAS1,IFIH1 |

| | | | | | | |
|---|---|---|---|---|---|---|
| WikiPathways | WP3617 | Photodynamic therapy-induced NF-kB survival signaling | 3 | 1.5 | 0.00021 | IL2,NFKB2,TNF |
| WikiPathways | WP4558 | Overview of interferons-mediated signaling pathway | 3 | 1.47 | 0.00024 | IFNG,STAT2,IFNB1 |
| WikiPathways | WP4329 | miRNA role in immune response in sepsis | 3 | 1.46 | 0.00025 | NFKB2,TNF,CCL4 |
| WikiPathways | WP5083 | Neuroinflammation and glutamatergic signaling | 4 | 1.19 | 0.00037 | IFNG,IL12B,NFKB2,TNF |
| WikiPathways | WP5198 | Inflammatory bowel disease signaling | 3 | 1.38 | 0.00037 | IL2,IFNG,TNF |
| WikiPathways | WP2882 | Nuclear receptors meta-pathway | 5 | 1 | 0.00044 | IL2,IFNG,IL12B,NFKB2,TNF |
| WikiPathways | WP4666 | Hepatitis B infection | 4 | 1.15 | 0.00046 | STAT2,IFNB1,TNF,IFIH1 |
| WikiPathways | WP1533 | Vitamin B12 metabolism | 3 | 1.3 | 0.00052 | IFNG,NFKB2,TNF |
| WikiPathways | WP3929 | Chemokine signaling pathway | 4 | 1.09 | 0.00063 | CXCL10,STAT2,CXCL9,CCL4 |
| WikiPathways | WP3863 | T-cell antigen receptor (TCR) pathway during Staphylococcus aureus in | 3 | 1.19 | 0.00091 | IL2,IFNG,TNF |
| WikiPathways | WP3624 | Lung fibrosis | 3 | 1.18 | 0.00094 | IL12B,TNF,CCL4 |
| WikiPathways | WP4655 | Cytosolic DNA-sensing pathway | 3 | 1.1 | 0.0014 | CXCL10,IFNB1,CCL4 |
| WikiPathways | WP15 | Selenium micronutrient network | 3 | 1.03 | 0.002 | IFNG,NFKB2,TNF |
| WikiPathways | WP4484 | Control of immune tolerance by vasoactive intestinal peptide | 2 | 1.16 | 0.0021 | IL2,IFNG |
| WikiPathways | WP4493 | Cells and molecules involved in local acute inflammatory response | 2 | 1.07 | 0.0032 | C3,TNF |
| WikiPathways | WP5055 | Burn wound healing | 3 | 0.93 | 0.0032 | NFKBIZ,IFNB1,TNF |
| WikiPathways | WP5174 | Ulcerative colitis signaling | 2 | 1.05 | 0.0035 | IFNG,TNF |
| WikiPathways | WP4478 | LTF danger signal response pathway | 2 | 1.03 | 0.0038 | IFNB1,TNF |
| WikiPathways | WP236 | Adipogenesis | 3 | 0.8 | 0.0063 | STAT2,TNF,SOCS1 |
| WikiPathways | WP3851 | TLR4 signaling and tolerance | 2 | 0.89 | 0.0075 | IFNB1,TNF |
| WikiPathways | WP453 | Inflammatory response pathway | 2 | 0.87 | 0.0084 | IL2,IFNG |
| WikiPathways | WP4481 | Resistin as a regulator of inflammation | 2 | 0.84 | 0.0098 | IL12B,TNF |
| WikiPathways | WP4496 | Signal transduction through IL1R | 2 | 0.84 | 0.0098 | IFNB1,TNF |
| WikiPathways | WP5038 | Mitochondrial immune response to SARS-CoV-2 | 2 | 0.84 | 0.0098 | NFKB2,IFIH1 |
| WikiPathways | WP2036 | TNF-related weak inducer of apoptosis (TWEAK) signaling pathway | 2 | 0.75 | 0.0147 | NFKB2,TNF |
| WikiPathways | WP4008 | NO/cGMP/PKG mediated neuroprotection | 2 | 0.72 | 0.0172 | IFNG,TNF |
| WikiPathways | WP560 | TGF-beta receptor signaling | 2 | 0.67 | 0.023 | IFNG,TNF |
| WikiPathways | WP585 | Interferon type I signaling pathways | 2 | 0.67 | 0.023 | STAT2,SOCS1 |
| WikiPathways | WP4816 | TGF-beta receptor signaling in skeletal dysplasias | 2 | 0.64 | 0.0255 | IFNG,TNF |
| WikiPathways | WP5222 | 2q13 copy number variation syndrome | 2 | 0.64 | 0.0267 | IL2,SOCS1 |
| WikiPathways | WP5130 | Th17 cell differentiation pathway | 2 | 0.59 | 0.0344 | IL2,IFNG |
| WikiPathways | WP254 | Apoptosis | 2 | 0.52 | 0.0482 | TNFSF10,TNF |
| Monarch | EFO:0007937 | Blood protein measurement | 11 | 0.48 | 0.0133 | TNFSF10,C3,CXCL10,HAVCR2,LILRB1,CXCL9,SLAMF7,CCL8,OAS1,TNF,CCL4 |
| DISEASES | DOID:612 | Primary immunodeficiency disease | 10 | 1.72 | 1.52E-07 | IL2,IFNG,IL12B,C3,STAT2,NFKB2,IFNB1,OAS1,TNF,IFIH1 |
| DISEASES | DOID:417 | Autoimmune disease | 8 | 1.63 | 2.44E-06 | IL2,IFNG,IL12B,NFKB2,IFNB1,OAS1,TNF,IFIH1 |
| DISEASES | DOID:8469 | Influenza | 4 | 2.23 | 6.39E-06 | IL2,IFNG,IFNB1,TNF |
| DISEASES | DOID:0080599 | Coronavirus infectious disease | 4 | 2.07 | 1.33E-05 | IFNG,CXCL10,IFNB1,TNF |
| DISEASES | DOID:635 | Acquired immunodeficiency syndrome | 3 | 1.57 | 0.00026 | IL2,IFNG,TNF |
| DISEASES | DOID:934 | Viral infectious disease | 5 | 1.31 | 0.00026 | IL2,IFNG,CXCL10,IFNB1,TNF |
| DISEASES | DOID:0080600 | COVID-19 | 3 | 1.4 | 0.00059 | IFNG,CXCL10,TNF |
| DISEASES | DOID:11162 | Respiratory failure | 3 | 1.38 | 0.00062 | IFNG,CXCL10,TNF |
| DISEASES | DOID:633 | Myositis | 3 | 1.37 | 0.00066 | CCL8,TNF,IFIH1 |
| DISEASES | DOID:2377 | Multiple sclerosis | 3 | 1.35 | 0.0007 | IFNG,IFNB1,TNF |
| DISEASES | DOID:0050117 | Disease by infectious agent | 6 | 0.95 | 0.00085 | IL2,IFNG,C3,CXCL10,IFNB1,TNF |
| DISEASES | DOID:0060032 | Autoimmune disease of musculoskeletal system | 5 | 1.06 | 0.00085 | IL2,IFNG,IL12B,TNF,IFIH1 |
| DISEASES | DOID:0060039 | Autoimmune disease of skin and connective tissue | 4 | 1.11 | 0.0013 | IL2,IFNG,IL12B,TNF |
| DISEASES | DOID:0080162 | Lupus nephritis | 2 | 1.23 | 0.0018 | C3,TNFSF13B |
| DISEASES | DOID:0060056 | Hypersensitivity reaction disease | 3 | 1.08 | 0.0026 | IFNG,C3,TNF |
| DISEASES | DOID:2921 | Glomerulonephritis | 3 | 1.08 | 0.0026 | C3,TNFSF13B,TNF |
| DISEASES | DOID:1579 | Respiratory system disease | 5 | 0.86 | 0.0027 | IL2,IFNG,CXCL10,IFNB1,TNF |
| DISEASES | DOID:2841 | Asthma | 3 | 1.05 | 0.003 | IL2,IFNG,TNF |
| DISEASES | DOID:2957 | Pulmonary tuberculosis | 2 | 1.1 | 0.0034 | IFNG,TNF |
| DISEASES | DOID:37 | Skin disease | 6 | 0.73 | 0.0034 | IL2,IFNG,IL12B,C3,TNF,IFIH1 |
| DISEASES | DOID:986 | Alopecia areata | 3 | 0.99 | 0.0039 | IL2,IFNG,TNF |
| DISEASES | DOID:11338 | Tetanus | 2 | 1.01 | 0.0052 | IL2,IFNG |
| DISEASES | DOID:0050589 | Inflammatory bowel disease | 3 | 0.9 | 0.0062 | IFNG,IL12B,TNF |
| DISEASES | DOID:11335 | Sarcoidosis | 2 | 0.98 | 0.0062 | IFNG,TNF |
| DISEASES | DOID:974 | Upper respiratory tract disease | 3 | 0.83 | 0.0087 | IL2,IFNG,TNF |
| DISEASES | DOID:614 | Lymphopenia | 2 | 0.91 | 0.0088 | IL2,IFNG |
| DISEASES | DOID:0060861 | Microphthalmia with limb anomalies | 2 | 0.88 | 0.0101 | OAS1,IFIH1 |
| DISEASES | DOID:11394 | Adult respiratory distress syndrome | 2 | 0.88 | 0.0101 | CXCL10,TNF |
| DISEASES | DOID:1205 | Allergic disease | 3 | 0.8 | 0.0101 | IL2,IFNG,TNF |
| DISEASES | DOID:9008 | Psoriatic arthritis | 2 | 0.86 | 0.0111 | IL12B,TNF |
| DISEASES | DOID:9471 | Meningitis | 2 | 0.83 | 0.0126 | IFNG,TNF |
| DISEASES | DOID:0050161 | Lower respiratory tract disease | 4 | 0.68 | 0.0128 | IL2,IFNG,CXCL10,TNF |
| DISEASES | DOID:0060033 | Autoimmune disease of peripheral nervous system | 2 | 0.81 | 0.0139 | IFNB1,TNF |
| DISEASES | DOID:8577 | Ulcerative colitis | 2 | 0.74 | 0.0197 | IFNG,TNF |

| | | | | | | |
|---|---|---|---|---|---|---|
| DISEASES | DOID:0050338 | Primary bacterial infectious disease | 3 | 0.64 | 0.0238 | IL2,IFNG,TNF |
| DISEASES | DOID:557 | Kidney disease | 4 | 0.57 | 0.024 | IL2,C3,TNFSF13B,TNF |
| DISEASES | DOID:7147 | Ankylosing spondylitis | 2 | 0.68 | 0.027 | IL12B,TNF |
| DISEASES | DOID:0080001 | Bone disease | 5 | 0.5 | 0.0276 | IL2,IFNG,IL12B,CCL8,TNF |
| DISEASES | DOID:848 | Arthritis | 3 | 0.6 | 0.0305 | IL2,IL12B,TNF |
| DISEASES | DOID:381 | Arthropathy | 2 | 0.64 | 0.0329 | CCL8,TNF |
| DISEASES | DOID:9351 | Diabetes mellitus | 3 | 0.58 | 0.0329 | OAS1,TNF,IFIH1 |
| DISEASES | DOID:552 | Pneumonia | 2 | 0.63 | 0.0342 | IFNG,TNF |
| DISEASES | DOID:8778 | Crohns disease | 2 | 0.63 | 0.0342 | IL12B,TNF |
| DISEASES | DOID:0060496 | Respiratory allergy | 2 | 0.63 | 0.0349 | IL2,IFNG |
| DISEASES | DOID:7 | Disease of anatomical entity | 13 | 0.28 | 0.042 | IL2,IFNG,IL12B,C3,CXCL10,STAT2,NFKB2,TNFSF13B,IFNB1,CCL8,OAS1,TNF,IFIH1 |
| TISSUES | BTO:0000570 | Hematopoietic system | 16 | 0.75 | 1.73E-06 | IL2,IFNG,TNFSF10,C3,HAVCR2,LILRB1,NFKBIZ,CXCL9,SLAMF7,NFKB2,TNFSF13B,OAS1,IFI35,TNF,CCL4,IFIH1 |
| TISSUES | BTO:0000878 | Mononuclear cell | 6 | 1.81 | 7.96E-06 | IL2,IFNG,NFKBIZ,CXCL9,TNFSF13B,TNF |
| TISSUES | BTO:0000876 | Monocyte | 5 | 1.62 | 5.17E-05 | IFNG,NFKBIZ,CXCL9,TNFSF13B,TNF |
| TISSUES | BTO:0000089 | Blood | 12 | 0.74 | 5.28E-05 | IL2,IFNG,C3,HAVCR2,LILRB1,NFKBIZ,CXCL9,TNFSF13B,OAS1,IFI35,TNF,CCL4 |
| TISSUES | BTO:0000751 | Leukocyte | 9 | 0.89 | 0.00011 | IL2,IFNG,LILRB1,NFKBIZ,CXCL9,TNFSF13B,OAS1,TNF,CCL4 |
| TISSUES | BTO:0004520 | Regulatory T-lymphocyte | 3 | 1.56 | 0.00024 | IL2,IFNG,TNF |
| TISSUES | BTO:0000753 | Lymphoid tissue | 10 | 0.63 | 0.0007 | IL2,IFNG,LILRB1,SLAMF7,NFKB2,TNFSF13B,OAS1,TNF,CCL4,IFIH1 |
| TISSUES | BTO:0000801 | Macrophage | 4 | 1.21 | 0.0007 | IL2,IFNG,SLAMF7,TNF |
| TISSUES | BTO:0000553 | Peripheral blood | 5 | 1.03 | 0.00088 | IL2,IFNG,NFKBIZ,TNFSF13B,TNF |
| TISSUES | BTO:0000737 | Leukemia cell line | 4 | 1.11 | 0.0011 | IL2,IFNG,TNFSF13B,TNF |
| TISSUES | BTO:0001025 | Peripheral blood mononuclear cell | 3 | 1.19 | 0.0014 | IL2,IFNG,TNF |
| TISSUES | BTO:0000130 | Neutrophil | 3 | 1.1 | 0.0022 | IFNG,TNFSF13B,TNF |
| TISSUES | BTO:0001034 | Peritoneal macrophage | 2 | 1.05 | 0.0042 | IFNG,TNF |
| TISSUES | BTO:0000155 | Bronchoalveolar lavage fluid | 2 | 1.01 | 0.0052 | IFNG,TNF |
| TISSUES | BTO:0005607 | Gamma delta T-lymphocyte | 2 | 1.01 | 0.0052 | IL2,IFNG |
| TISSUES | BTO:0000802 | Alveolar macrophage | 2 | 0.98 | 0.0059 | IFNG,TNF |
| TISSUES | BTO:0004732 | Bone marrow-derived macrophage | 2 | 0.98 | 0.0059 | IFNG,TNF |
| TISSUES | BTO:0006110 | M1 macrophage | 2 | 0.96 | 0.0066 | IFNG,TNF |
| TISSUES | BTO:0001045 | T-lymphocyte cell line | 2 | 0.94 | 0.0073 | IL2,IFNG |
| TISSUES | BTO:0000775 | Lymphocyte | 6 | 0.59 | 0.0083 | IL2,IFNG,LILRB1,OAS1,TNF,CCL4 |
| TISSUES | BTO:0003435 | Memory T-lymphocyte | 2 | 0.89 | 0.009 | IL2,IFNG |
| TISSUES | BTO:0001598 | Splenocyte | 2 | 0.88 | 0.0097 | IL2,IFNG |
| TISSUES | BTO:0005871 | Naive T-lymphocyte | 2 | 0.88 | 0.0097 | IL2,IFNG |
| TISSUES | BTO:0002332 | Monocytic leukemia cell line | 2 | 0.86 | 0.0104 | IFNG,TNF |
| TISSUES | BTO:0003861 | Inflammatory cell | 2 | 0.86 | 0.0104 | IFNG,TNF |
| TISSUES | BTO:0001472 | Peritoneum | 2 | 0.85 | 0.0111 | IFNG,TNF |
| TISSUES | BTO:0002278 | Macrophage cell line | 2 | 0.83 | 0.0121 | IFNG,TNF |
| TISSUES | BTO:0000782 | T-lymphocyte | 4 | 0.62 | 0.0162 | IL2,IFNG,TNF,CCL4 |
| TISSUES | BTO:0001281 | Spleen | 5 | 0.54 | 0.018 | IL2,IFNG,SLAMF7,TNFSF13B,IFIH1 |
| TISSUES | BTO:0001043 | Adult | 2 | 0.75 | 0.0184 | IFNG,TNF |
| TISSUES | BTO:0001522 | B-lymphocyte cell line | 3 | 0.66 | 0.0198 | IFNG,LILRB1,SOCS1 |
| TISSUES | BTO:0000289 | Cytotoxic T-lymphocyte | 2 | 0.69 | 0.0243 | IL2,IFNG |
| TISSUES | BTO:0002417 | Helper T-lymphocyte | 2 | 0.66 | 0.0293 | IL2,IFNG |
| TISSUES | BTO:0002144 | Acute lymphoblastic leukemia cell line | 2 | 0.59 | 0.0413 | IL2,TNFSF13B |
| Subcellular local | GOCC:0043514 | interleukin-12 complex | 5 | 2.74 | 2.85E-07 | IL2,IFNG,IL12B,CXCL10,TNF |
| Subcellular local | GOCC:0070743 | interleukin-23 complex | 4 | 2.27 | 5.12E-06 | IL2,IFNG,IL12B,TNF |
| Subcellular local | GOCC:0071735 | IgG immunoglobulin complex | 3 | 1.44 | 0.00047 | IL2,IFNG,TNF |
| Subcellular local | GOCC:0099126 | Transforming growth factor beta complex | 3 | 1.29 | 0.00097 | IL2,IFNG,TNF |
| Subcellular local | GOCC:0071159 | NF-kappaB complex | 3 | 1.18 | 0.0017 | IFNG,NFKB2,TNF |
| Subcellular local | GOCC:0097072 | Interferon regulatory factor 3 complex | 2 | 0.87 | 0.0109 | IFNB1,IFIH1 |
| Subcellular local | GOCC:1990231 | STING complex | 2 | 0.81 | 0.0149 | IFNB1,IFIH1 |
| Subcellular local | GOCC:0005576 | Extracellular region | 13 | 0.73 | 3.15E-05 | IL2,IFNG,IL12B,TNFSF10,C3,CXCL10,CXCL9,TNFSF13B,IFNB1,CCL8,IFI35,TNF,CCL4 |
| Subcellular local | GOCC:0042101 | T cell receptor complex | 2 | 0.66 | 0.0307 | IL2,IFNG |
| Subcellular local | GOCC:0005615 | Extracellular space | 8 | 0.63 | 0.003 | IL2,IFNG,IL12B,TNFSF10,C3,CXCL10,IFI35,TNF |
| Subcellular local | GOCC:0098802 | Plasma membrane signaling receptor complex | 4 | 0.55 | 0.0307 | IL2,IFNG,IL12B,IFNB1 |
| Subcellular local | GOCC:0005887 | Integral component of plasma membrane | 6 | 0.46 | 0.0307 | IL2,IL12B,TNFSF10,C3,IFNB1,TNF |
| Subcellular local | GOCC:0016020 | Membrane | 15 | 0.28 | 0.0307 | IL2,IFNG,IL12B,TNFSF10,C3,HAVCR2,STAT2,LILRB1,SLAMF7,TNFSF13B,IFNB1,IFI35,TNF,SOCS1,IFIH1 |
| UniProt Keywor | KW-0202 | Cytokine | 11 | 3.88 | 3.40E-14 | IL2,IFNG,IL12B,TNFSF10,CXCL10,CXCL9,TNFSF13B,IFNB1,CCL8,TNF,CCL4 |
| UniProt Keywor | KW-0964 | Secreted | 14 | 0.95 | 2.07E-07 | IL2,IFNG,IL12B,TNFSF10,C3,CXCL10,LILRB1,CXCL9,TNFSF13B,IFNB1,CCL8,OAS1,TNF,CCL4 |
| UniProt Keywor | KW-0395 | Inflammatory response | 6 | 1.76 | 5.40E-06 | C3,CXCL10,HAVCR2,CXCL9,CCL8,CCL4 |
| UniProt Keywor | KW-0391 | Immunity | 8 | 1.22 | 1.23E-05 | IL2,C3,HAVCR2,LILRB1,SLAMF7,TNFSF13B,OAS1,IFIH1 |
| UniProt Keywor | KW-0051 | Antiviral defense | 5 | 1.55 | 4.02E-05 | IFNG,STAT2,IFNB1,OAS1,IFIH1 |
| UniProt Keywor | KW-1015 | Disulfide bond | 13 | 0.46 | 0.0012 | IL2,IL12B,C3,CXCL10,HAVCR2,LILRB1,CXCL9,SLAMF7,TNFSF13B,IFNB1,CCL8,TNF,CCL4 |
| UniProt Keywor | KW-0399 | Innate immunity | 5 | 0.82 | 0.0027 | C3,HAVCR2,SLAMF7,OAS1,IFIH1 |
| UniProt Keywor | KW-1064 | Adaptive immunity | 4 | 0.9 | 0.003 | IL2,HAVCR2,LILRB1,SLAMF7 |
| UniProt Keywor | KW-0732 | Signal | 12 | 0.41 | 0.0042 | IL2,IFNG,IL12B,C3,CXCL10,HAVCR2,LILRB1,CXCL9,SLAMF7,IFNB1,CCL8,CCL4 |

| | | | | | | |
|---|---|---|---|---|---|---|
| UniProt Keywor | KW-0145 | Chemotaxis | 3 | 0.76 | 0.0112 | CXCL10,CCL8,CCL4 |
| InterPro | IPR001811 | Chemokine interleukin-8-like domain | 4 | 1.17 | 0.0016 | CXCL10,CXCL9,CCL8,CCL4 |
| InterPro | IPR036048 | Chemokine interleukin-8-like superfamily | 4 | 1.17 | 0.0016 | CXCL10,CXCL9,CCL8,CCL4 |
| InterPro | IPR006052 | Tumour necrosis factor domain | 3 | 1.05 | 0.0038 | TNFSF10,TNFSF13B,TNF |
| SMART | SM00199 | Intercrine alpha family (small cytokine C-X-C) (chemokine CXC), | 4 | 1.57 | 0.00015 | CXCL10,CXCL9,CCL8,CCL4 |
| SMART | SM00207 | Tumour necrosis factor family, | 3 | 1.37 | 0.00065 | TNFSF10,TNFSF13B,TNF |
