## Supplementary material for "Functional alterations of immune gene expression in ICU and non-ICU patients with Legionnaires’ disease, a prospective observational study": Figure S1

**Table S4: Details of all enriched terms and pathways found in ICU-LD patients**

See dedicated PDF file

**Table S5: Details of all enriched terms and pathways found in non-ICU-LD patients**

See dedicated PDF file

**Figure S1: Comparison of DEG between ICU-LD and LD-unrelated-SS patients** **after** **LPS stimulation.** Comparison of the number of less-and more-expressed DEGs in ICU-LD and LD-unrelated-SS patients (A). Venn diagrams of less-expressed and more-expressed DEGs in ICU-LD and non-LD-unrelated-SS patients (B). List of less-and more-expressed DEGs in ICU-LD and LD-unrelated-SS patients (C). The blue to red gradient colour indicates lowest to highest log2(FC) values. White boxes with black crosses indicate genes that were not differentially expressed between ICU-LD patients and HVs or LD-unrelated-SS patients and HVs. Median gene expressions boxplot of the 44 less-expressed genes in ICU-LD compared with LD-unrelated-SS patients (D). ns: not significant
