## Supplementary figures and images for "Functional alterations of immune gene expression in ICU and non-ICU patients with Legionnaires’ disease, a prospective observational study"

### Figure S1

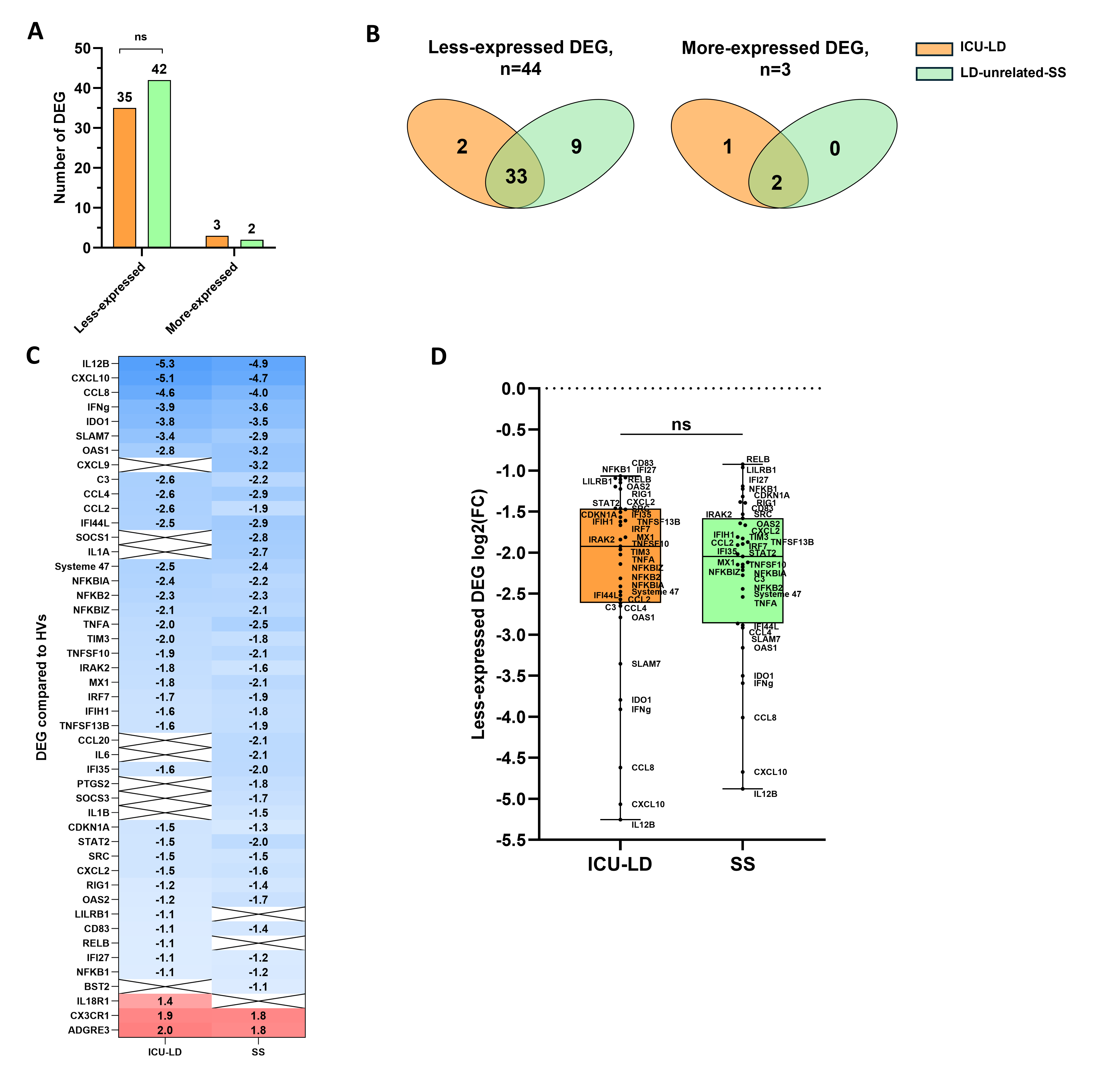
